# Nationwide genomic biobank in Mexico unravels demographic history and complex trait architecture from 6,057 individuals

**DOI:** 10.1101/2022.07.11.499652

**Authors:** Mashaal Sohail, Amanda Y. Chong, Consuelo D. Quinto-Cortes, María J. Palma-Martínez, Aaron Ragsdale, Santiago G. Medina-Muñoz, Carmina Barberena-Jonas, Guadalupe Delgado-Sánchez, Luis Pablo Cruz-Hervert, Leticia Ferreyra-Reyes, Elizabeth Ferreira-Guerrero, Norma Mongua-Rodríguez, Andrés Jimenez-Kaufmann, Hortensia Moreno-Macías, Carlos A. Aguilar-Salinas, Kathryn Auckland, Adrián Cortés, Víctor Acuña-Alonzo, Alexander G. Ioannidis, Christopher R. Gignoux, Genevieve L. Wojcik, Selene L. Fernández-Valverde, Adrian V.S. Hill, María Teresa Tusié-Luna, Alexander J. Mentzer, John Novembre, Lourdes García-García, Andrés Moreno-Estrada

**Affiliations:** Laboratorio Nacional de Genómica para la Biodiversidad (LANGEBIO), Unidad de Genómica Avanzada (UGA), CINVESTAV, Irapuato, Guanajuato, México; Department of Human Genetics, University of Chicago, Chicago, IL, USA; Centro de Ciencias Genómicas (CCG), Universidad Nacional Autónoma de México (UNAM), Cuernavaca, Morelos, México; The Wellcome Centre for Human Genetics, University of Oxford, Oxford, UK; Instituto Nacional de Salud Pública (INSP), Cuernavaca, Morelos, México; Facultad de Odontología, Universidad Nacional Autónoma de México (UNAM), Ciudad de México, México; Unidad de Biología Molecular y Medicina Genómica, UNAM-INCMNSZ, México City, México; Universidad Autónoma Metropolitana, México City, México; Division de Nutrición, Instituto Nacional de Ciencias Médicas y Nutrición Salvador Zubirán, Mexico City, México; Big Data Institute, Li Ka Shing Centre for Health Information and Discovery, University of Oxford, Oxford, UK; Escuela Nacional de Antropología e Historia (ENAH), Mexico City, México; Department of Biomedical Data Science, Stanford University, Stanford, CA, USA; Colorado Center for Personalized Medicine, University of Colorado Anschutz Medical Campus, Aurora, CO, USA; Department of Epidemiology, Johns Hopkins Bloomberg School of Public Health, Baltimore, MD, USA; The Jenner Institute, University of Oxford, Oxford, UK; Department of Ecology and Evolution, University of Chicago, Chicago, IL, USA

## Abstract

Latin America continues to be severely underrepresented in genomics research, and fine-scale genetic histories as well as complex trait architectures remain hidden due to the lack of Big Data. To fill this gap, the Mexican Biobank project genotyped 1.8 million markers in 6,057 individuals from 32 states and 898 sampling localities across Mexico with linked complex trait and disease information creating a valuable nationwide genotype-phenotype database. Through a suite of state-of-the-art methods for ancestry deconvolution and inference of identity-by-descent (IBD) segments, we inferred detailed ancestral histories for the last 200 generations in different Mesoamerican regions, unraveling native and colonial/post-colonial demographic dynamics. We observed large variations in runs of homozygosity (ROH) among genomic regions with different ancestral origins reflecting their demographic histories, which also affect the distribution of rare deleterious variants across Mexico. We analyzed a range of biomedical complex traits and identified significant genetic and environmental factors explaining their variation, such as ROH found to be significant predictors for trait variation in BMI and triglycerides.

---

The genetic architecture of complex traits in admixed genomes cannot be understood outside the context of their underlying histories. Present-day Mexico covers seven Mesoamerican regions with rich civilizational histories^1^. Archaeology and anthropology regionalize Mexico into the north of Mexico, the north of Mesoamerica, the center, occident and gulf of Mexico, Oaxaca and the Mayan region^2^(Fig. 1a). These are based on noting specific pre-Hispanic civilizations and cultures, which began flourishing very early in the Mayan region, in Oaxaca, in the occident and in the gulf of Mexico^3^, and later in the center and north of Mesoamerica. Such histories have also been used to classify Mesoamerican chronology into preclassical, classical, postclassical, colonial, and postcolonial periods.

**Figure 1.**
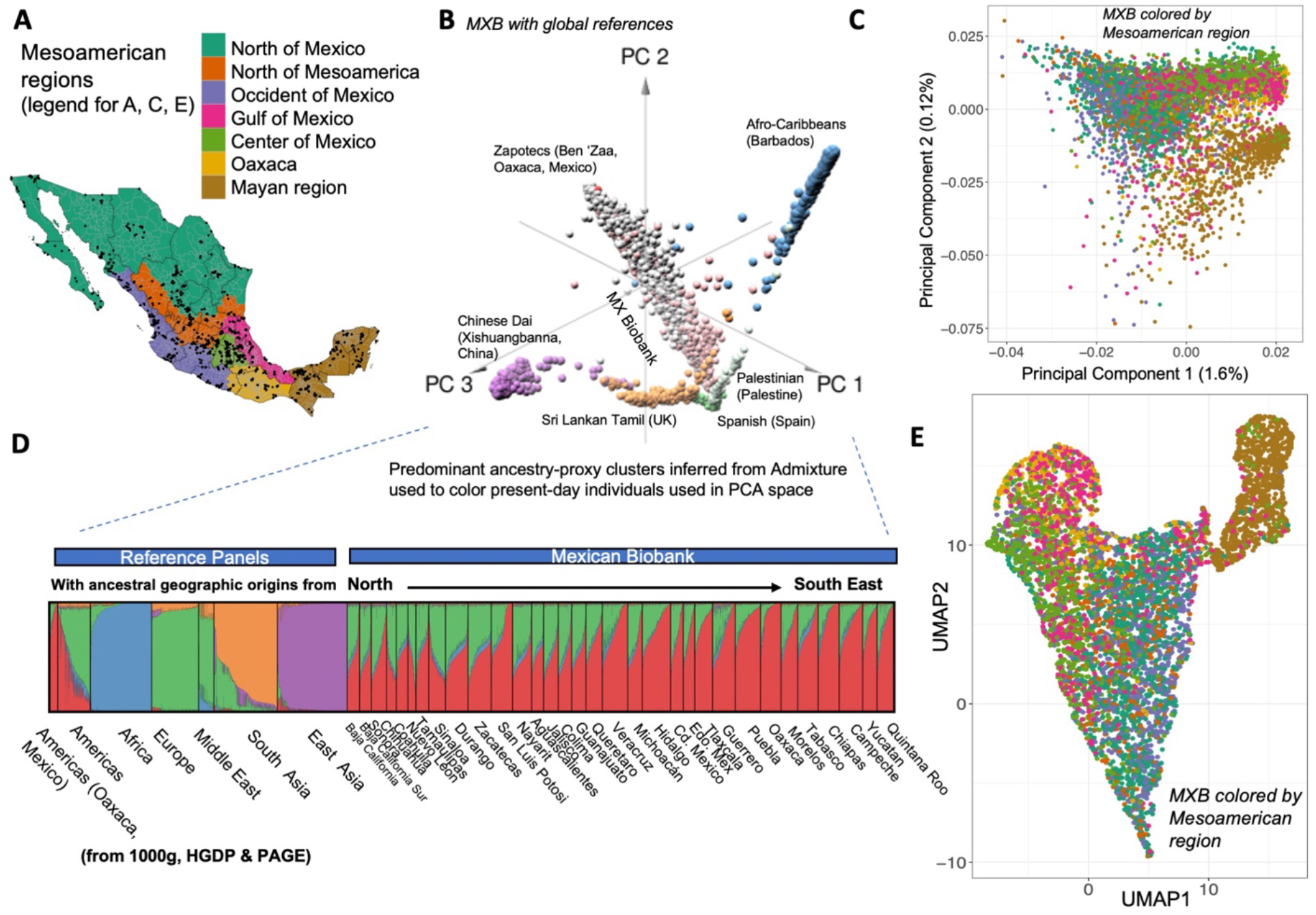
Visualizing genetic structure across the geography of the Mexican Biobank as inferred by dimensionality reduction and unsupervised clustering. A) Mexico regionalized into Mesoamerican regions according to anthropological and archaeological context. B) Principal components analysis (PCA) of MXB with reference global data from the 1000G project, HGDP and PAGE. Some specific sampled cohorts are labelled across the plot to orient the reader. C) PCA with only MXB colored and regionalized into Mesoamerican regions. D) Unsupervised clustering using Admixture and global reference panels (same as in B). E) UMAP analysis of MXB colored by Mesoamerican regions.

In the last five hundred years, Spanish colonization has left an indelible mark on this native tapestry. In a colonial context, ancestries that trace to European, African and Asian sources can be identified in living Mexicans, however they vary in structure and timing between Mesoamerican regions^45-89-11^. The heterogeneity of such a mixture at a genetic level has been characterized, revealing extensive fine-scale population substructure and ancestry sources across Mexico^12–16^. These studies have also identified genes potentially under selection for some traits in different native groups^12,15^.

Further, such varying genetic histories, as captured by ancestry distributions, have been shown to impact variation in complex traits in Mexicans in traits such as lung force capacity^12^, and a number of other complex traits and diseases^17^. Nevertheless, a large gap remains in the representation of Mexicans from across Mexico in cohorts with linked genotypes and phenotypes, which could enable finer-scale studies of genetic history and a better understanding of complex trait architecture among individuals with diverse ancestries from the Americas and those living in rural areas^18^. Past efforts have been limited to studying individuals from the United States and Mexico City and have not simultaneously modelled the influence on complex trait variation of a rich array of genetic and environmental factors as is possible with a nationwide Biobank.

To bridge this gap, we launched the Mexican Biobank (MXB) project, densely genotyping 6,057 individuals from all 32 states across Mexico (Fig. S1-S2) recruited as part of the National Health Survey in 2000 (ENSA2000), which sampled more than 40,000 participants nationwide. To select the samples for genomic and biochemical characterization, we enriched for those individuals that can speak an indigenous language in each state while maximizing the representation of rural localities (~70% of the MXB out of a total of 898 localities, Fig. S2-S5) to increase the representation of indigenous ancestries. The MXB is 70% female and comprised of individuals born between 1910 and 1980, all sampled in the year 2000^18^(Table S1). These individuals were genotyped at ~1.8 million SNPs and have linked information for traits such as height, BMI, triglycerides, glucose, cholesterol, blood pressure and various socioeconomic and biogeographical markers (Table S2).

Here, we leverage rich archaeological and anthropological information to guide a regionalized analysis of Mexico, and harness the power of local ancestry estimation genome-wide and segments of identity-by-descent (IBD) to decipher fine-scale genetic histories using ancestry-specific approaches to denote origins and historical population size changes^19,20^. We reveal a very heterogeneous landscape of both, painting a genetically informed picture of varying demographic trajectories of Mesoamerican civilizations, as well as colonial migrations and dynamics in different regions of Mexico. We further investigate the role of these evolutionary histories as captured by proxies of genetic ancestries in shaping genetic variation and complex traits patterns in Mexico today. We show that these histories result in marked geographic and ancestry-specific patterns in the distributions of runs of homozygosity (ROH) and of the genomic burden of rare deleterious mutations.

Lastly, we study the impact of these histories which could associate certain trait-relevant genotypes with certain genetic backgrounds, along with portions of the genome in ROH and other sociocultural and biogeographical factors capturing environmental context, on creating trait variation in complex and medically-relevant traits such as height, BMI, triglycerides, glucose levels, and others in Mexico. Our results can help guide sampling and design for future genetic mapping efforts by determining which environmental and genetic axes maximize trait variation, to help increase power in genome-wide association studies. They can also help determine cases where environmental interventions are more likely to bring a desired improvement in public health.

## Genetic structure across Mexico is shaped by native diversity and historical migrations

We begin by excavating the population structure in the MXB at different geographic resolutions and time-scales. Principal components analysis (PCA)^21^ captures predominant axes of genetic similarity. Further, a proxy for genetic ancestries from different regions can be quantified using ADMIXTURE^22^ (see note on genetic ancestries in methods). When we visualize the Mexican biobank samples using PCA with individuals from around the world (1000 Genomes^23^, HGDP^24^, and PAGE^25^), we find that most Mexican individuals lie on a cline between living Europeans and indigenous Americans, which we interpret as reflecting the history of admixture in Mexico since Spanish colonization (Fig. 1B, Figs. S6). We also observe a “pull” towards present-day Africans, likely reflecting the genetic impact of the trans-Atlantic slave trade during the colonial period and subsequent migrations that brought many Africans to Mexico. When analyzed alone, the MXB individuals show a striking population substructure delineation between the Mayan region and the rest of the country (Fig. 1C, Fig. 1E, Figs. S7-S17). In the rest of the regions, only a subtle genetic substructure mirroring Mesoamerican geography is visible in the MXB, likely reflecting the effects of movement and mating among the different regions sampled in the year 2000.

We infer ancestry proxies at different geographic resolutions reflecting mating dynamics in the colonial and post-colonial periods by analyzing MXB with global individuals as well as with only native individuals using the software ADMIXTURE (Fig. 1D, Table S3, Fig. S11). We observe that individuals in the Mexican Biobank are inferred to be admixed with varying degrees of ancestries that are found most abundantly in individuals of the Americas (“American ancestries”) and Europe (“European ancestries”). Higher levels of ancestries from the Americas were inferred in the central and southern states of Mexico, compared to the northern states. We observe the largest genetic differentiation as measured using Fst along a north to southeast cline (Fig. S14-17). We observe some ancestries from Africa in individuals found in every single state (Table S3). We observe that only 3 states in the north (Chihuahua, Nuevo Leon and Sinaloa) have more ancestries on average from Europe than ancestries from the Americas, with ancestries from the Americas being the majority ancestries in every other state (Table S3). Lastly, we note the presence of a small but significant proportion of ancestries from East Asia in almost every state (0-2.3%), the highest in the state of Guerrero (2.3%), and an even more modest but significant amount of ancestries from South Asia in the majority of states as well (0-0.8%).

We use an ancestry-specific PCA or MDS approach to pinpoint the origins of the ancestries from the Americas, Africa, and Asia observed in the Mexican Biobank within those regions. For ancestries from the Americas, we observe that such ancestries tend to originate from indigenous cultures predominant in the region an individual is from (Fig. S18, Table S4). For example, such ancestries in the Yucatan peninsula originate from the Maya and Tzotzil. We observe that most ancestries from Africa in Mexico originate from West Africa^26^, in agreement with historical records of shipping voyages from the trans-Atlantic slave trade (Fig. S19)^4^. For the ancestries from East Asia, we find for individuals in Guerrero, such segments projecting to East Asian regions that were linked to the Manila Galleon trade, as reported in preliminary findings^16^. In contrast, for individuals from northern states such as Chihuahua, such segments project to China and Japan, likely reflecting later migrations from East Asia to Mexico (Fig. S20). Similarly, for ancestries from South Asia, as illustrated for individuals from Guerrero, the landing state of the Manila Galleon, we observe diverse roots in South Asia (Fig. S21-S22).

The identification of individuals with genetic ancestries from East Asia in Mexico dating to the Manila Galleon trade agrees with one other recent study^16^. These genetic observations are plausibly explained by the poorly appreciated history of Mexico’s Asian population^5–8^. Using voyage records, slaving documents, and other sources, historians have documented arrivals from the Manila port in the Spanish Philippines to the Acapulco port in Mexico through the 16^th^ and 17^th^ centuries, with origins as diverse as the Philippines, Indonesia, Malaysia, India, Bengal, and Sri Lanka (though they were collectively referred to as “Chinos” by the colonists). They entered from the Acapulco port, but moved through most of Mexico, with even what was called the “China Road” existing between Acapulco and Mexico City. There have also been later 19^th^ and 20^th^ century migrations from China and Japan, especially to the north of Mexico, and inheritance from these ancestors likely explains part of the ancestries from East Asia we observe in the northern states of the MXB today^9–11^.

## Ancestry-specific IBD tracts recover 200 generations of genetic history within Mexico

Apart from the small amount of recent ancestry from Asia discussed above, the ancestry of contemporary Mexicans arises predominantly from lineages that would have been found in Central America, Western Europe, and West Africa prior to 15^th^ century. Each of these sources had different demographic histories prior to and after their arrival in present-day Mexico. To reveal the more recent history of population sizes of these three ancestries, we analyze identity-by-descent segments^27^ overlapped with their corresponding local ancestry inference^19^. We use this approach to estimate effective population size (*N_e_*) trajectories 200 generations into the past for these ancestries in the Mexican Biobank as a whole, as well as in specific Mesoamerican regions (Figure 2). This analysis helps us reveal genetic histories of present-day Mexicans, which is of anthropological interest, as well as relevant for patterns of genetic and complex trait variation as shown in later sections.

**Figure 2.**
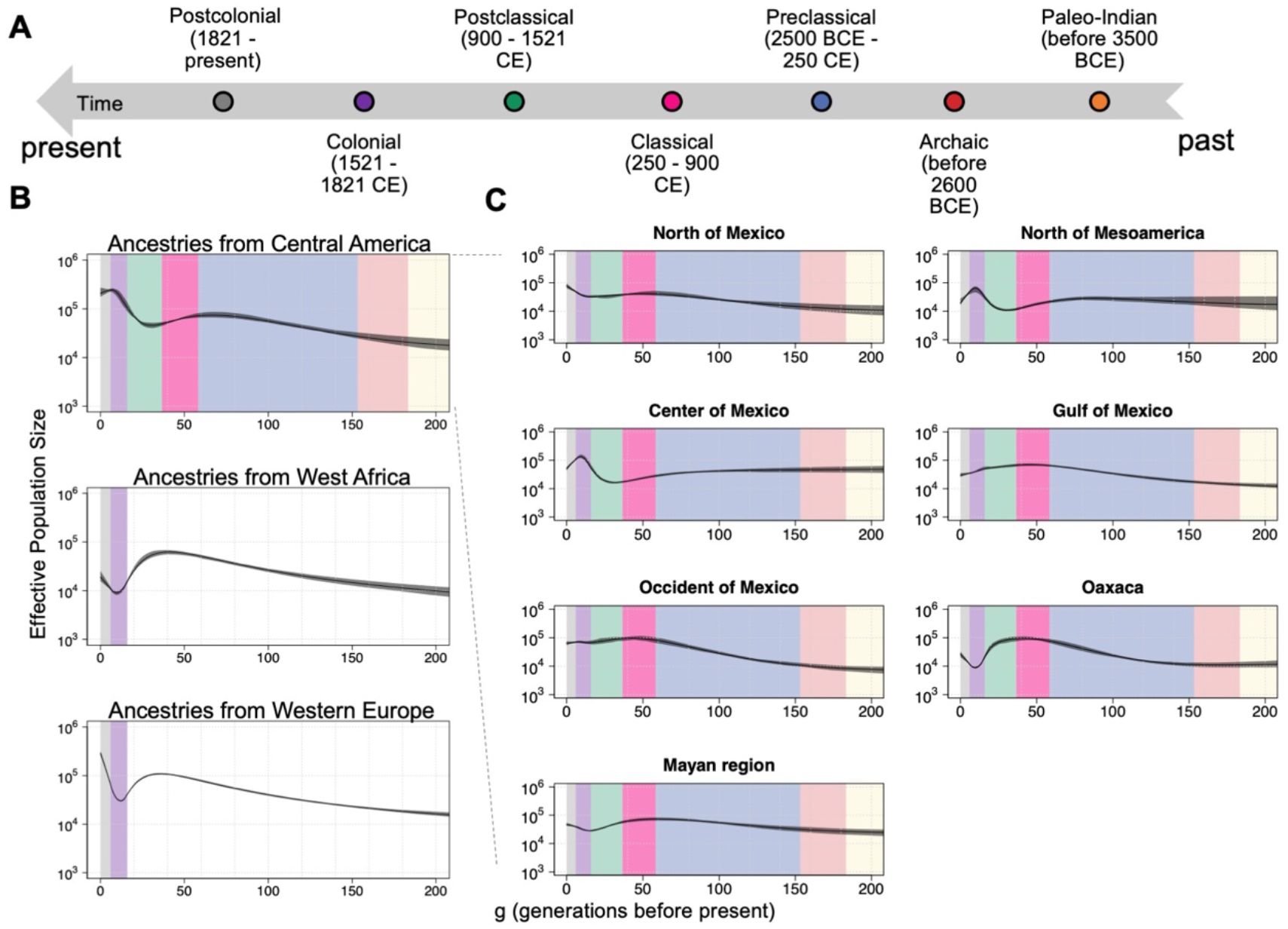
Effective population size (*N_e_*) changes inferred using identity-by-descent (IBD) tracts across ancestries and geographies reveal the different histories present within Mexico. A) Mesoamerican chronology coloring different periods in Mesoamerican history using anthropological and archaeological context B) Ancestry-specific effective population size changes over past 200 generations across Mexico (colored by chronology from A assuming 30 years per generation (see Figs. S23-27 for other generation intervals and ancestries). C) Ancestry-specific effective population changes over time for ancestries from Central America in different Mesoamerican regions of Mexico.

In the entire MXB, contextualized using Mesoamerican chronology (Fig. 2A), we find that, for indigenous lineages, (i.e., those present in the area before the arrival of the Spaniards), the effective population size went through a slow and steady decline in the classical period (250 – 900 CE). This decline was followed by an increase in the postclassical period (900 – 1521 CE), right before the arrival of the Spaniards, and then a decline later in the colonial period (1521 – 1821 CE) and in the post-colonial period (1821 – present) (Fig. 2B).

Further, we observe fine-scale structure in *N_e_* trajectories for indigenous lineages which we interpret in the context of the different cultural histories of Mesoamerican regions (Fig. 2C)^1^. Starting chronologically, archaeologists document that Mesoamerican civilizations flourished very early in the Mayan region, in Oaxaca, in the occident and in the gulf of Mexico, where we also observe large *N_e_* already in the classical period^3^. For example, in the gulf, where we observe high *N_e_* since the pre-classical period (2500 BCE – 250 CE), there is archaeological evidence, among a myriad of other groups, of the Olmecs in the pre-classical period, the Totonacs in the classical period, and the Huastecs in the post-classical period^28^. In Oaxaca, we observe *N_e_* rapidly growing in the pre-classical to the classical period, in line with archaeological inferences that the Zapotecs were already starting to create sedentary settlements in the pre-classical period followed by a rise in social and political structures in the classical period. This was followed by a more militaristic period in the post-classical causing warfare^29^, and our genetic evidence suggests a significant population decline toward the end of the post-classical period. In the Yucatan peninsula, the Maya had prominent civilizational spread in the classical period (peak *N_e_* observed), and started going through a slow decline only in the post-classical period due to what archaeologists have inferred as a combination of different political and ecological factors, and this trajectory is supported in the *N_e_* trend^3^. We further observe that native groups in both Oaxaca and the Mayan peninsula started to increase in population size again through the colonial and post-colonial (1821 – present) periods, after the arrival of the Spaniards.

This is in contrast with the center and north of Mesoamerica, where the Aztec empire had a strong-hold most recently and where we see increasing *N_e_* in the post-classical right before the arrival of the Spaniards and into part of the colonial period, after which we start to see a population decline in *N_e_*. Thus, the decline in *N_e_* after the arrival of the Spaniards is most prominent in the center and north of Mesoamerica, and is actually followed by an increase in Oaxaca and the Mayan region, where native ancestries from Central America are most prevalent today as evidenced by the Admixture analysis (Table S3). As generational time can vary, we present our analysis at two extremes of 20 and 30 years per generation^30^ (Fig. S23 and 2C, respectively).

We observe that ancestries from Western Europe that entered the contemporary Mexican gene pool went through a sharp decline in effective population size during the colonial period. The extent of the founder effect varied by region, with the strongest effect seen in Oaxaca and the Mayan region (Fig. S24, S25). Similarly, ancestries from Western Africa in Mexico revealed stronger founder effects that varied by region with *N_e_* ranging between 10^3^ and 10^4^ in the colonial period. The population size in the post-colonial period continued to grow in some regions such as the occident and north of Mexico and the Mayan region, compared to others (Fig. S26, S27).

## Demographic histories impact patterns of genetic variation in Mexico

### Small ROH prevalence is correlated with ancestry proxies

We next analyze the patterns of ROH in the MXB and their relationship with geography and ancestry proxies. ROH patterns help further illuminate demographic and mating histories of Mexicans^31^, and are relevant for variation in complex traits if trait-relevant variation is partially recessive^32^. We identify ROH (≥ 1 Mb) in the Mexican Biobank and observe that both the number of ROHs and the total length of ROH per individual increases as we move from north to southeast in the country (Fig. S28-29). We confirm that this is primarily due to individuals with more genetic ancestries from Central America also having more ROH in their genomes (Fig. 3A, Figs. S30-31, Table S5).

**Figure 3.**
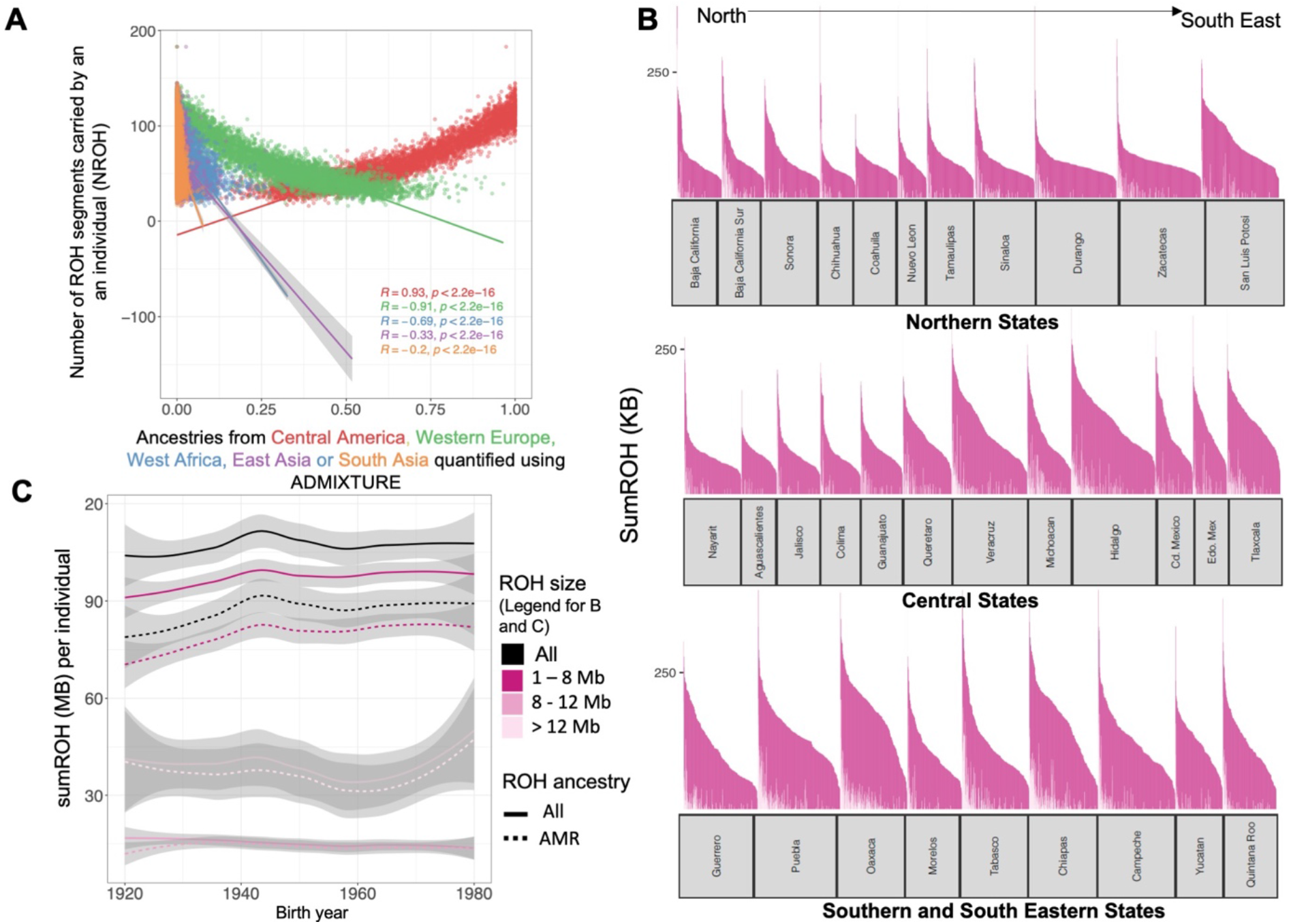
Analysis of runs of homozygosity (ROH) in each individual in MXB across ancestries, geographies and birth year reveals the role of ancient and recent demographic movements to and within Mexico. A) ROH are correlated with ancestries from global region in each individual reflecting the impact of varied and shared demographic histories and bottlenecks. B) Distribution of ROH segments of different sizes for each Northern, Central, Southern and Southeastern state. The y-axis was truncated to aid visualization, truncating the first bar for some states. C) ROH as a function of birth year. Solid lines show ROH overall, and dashed lines indicate ROH overlapping ancestries from Central America. ROH are divided into small, medium and large ROH same as in (B).

Next, we asked whether this signal of higher ROH associated with higher ancestry from Central America is due to historical bottlenecks and small population sizes, or due to consanguinity. A bottleneck event or a long-term small population size will result in a large number of small ROH^33^. Consanguinity or marriage between relatives would instead result in fewer but longer ROH^33^. To answer this question, we analyze the total length of ROH per individual in each state, after first partitioning ROH by size (Fig. 3B). We observe that there are more ROH per individual moving southwards in Mexico in large part due to small ROH (smaller than those expected from recent consanguinity e.g., < 8 Mb), implying that bottlenecks and small population sizes rather than consanguinity have been largely responsible for more ROH in individuals with higher ancestries from Central America.

Lastly, we observe that ROH that are found on segments of the genome from Central America are more frequently found in younger individuals compared to older individuals (Spearman’s rho = 0.31, p = 0.016) (Fig. 3C). We also verified that this correlation with birth year primarily derives from small ROH (rho = 0.35, p = 0.006), and small ROH found on genomic segments from the Americas (rho = 0.39, p = 0.002) (Fig. 3C). This result is at least partly due to younger individuals carrying more ancestries from Central America compared to older individuals, especially in the rural localities (Fig. S51), and agrees with recent observations about ancestry and ROH made in Mexican-Americans^17^. The observation of higher ancestries from Central America in younger individuals in rural areas may be due to either individuals in rural areas procreating at a higher rate, or individuals with other ancestries moving out from rural to urban areas. The observation of a larger number of small ROH in younger individuals in the MXB is relevant for parsing the genetic architecture of complex traits and diseases, especially those with a recessive component.

### Rare deleterious variant burden is correlated with ancestry proxies

We also investigated the effects of demographic history on the frequency distribution of genetic variants. If such an effect exists for variants that contribute to trait variation, it would imply varying genetic architectures for some traits that may be captured by ancestry proxies. This analysis is motivated by previous theoretical and empirical work showing that undergoing a bottleneck changes the allele frequency distribution in the group that experienced the bottleneck^23,34,35^. In particular, rare variants are lost or increase in frequency after the bottleneck.

We can evaluate this effect within and between genes in the genome by calculating the genome load of genetic variants, or mutation burden. We calculated the genome-wide mutation burden by summing the derived alleles at each SNP in the genome for different types of variants (intergenic, synonymous, putatively deleterious). When considering only rare variants (DAF<= 5%), we observe that the total number of rare variants is negatively correlated with ancestries from Central America, while it is positively correlated with ancestries from Western Europe and West Africa (Fig. 4). Thus, individuals with higher ancestries from Central America carry fewer rare variants likely due to a history of more bottlenecking events compared to individuals with higher ancestries from Western Europe and West Africa. We observed the same general pattern for different types of variants. However, this effect of varying demographic histories on variant frequencies is strongest for putatively neutral variants (Fig. 4). We have verified these observations for rare variants with whole-genome sequences from the subset of Mexicans living in Los Angeles (MXL) from the 1000 Genomes Project to rule out ascertainment biases due to the array genotyping as the source of this effect (Fig. S50).

**Figure 4.**
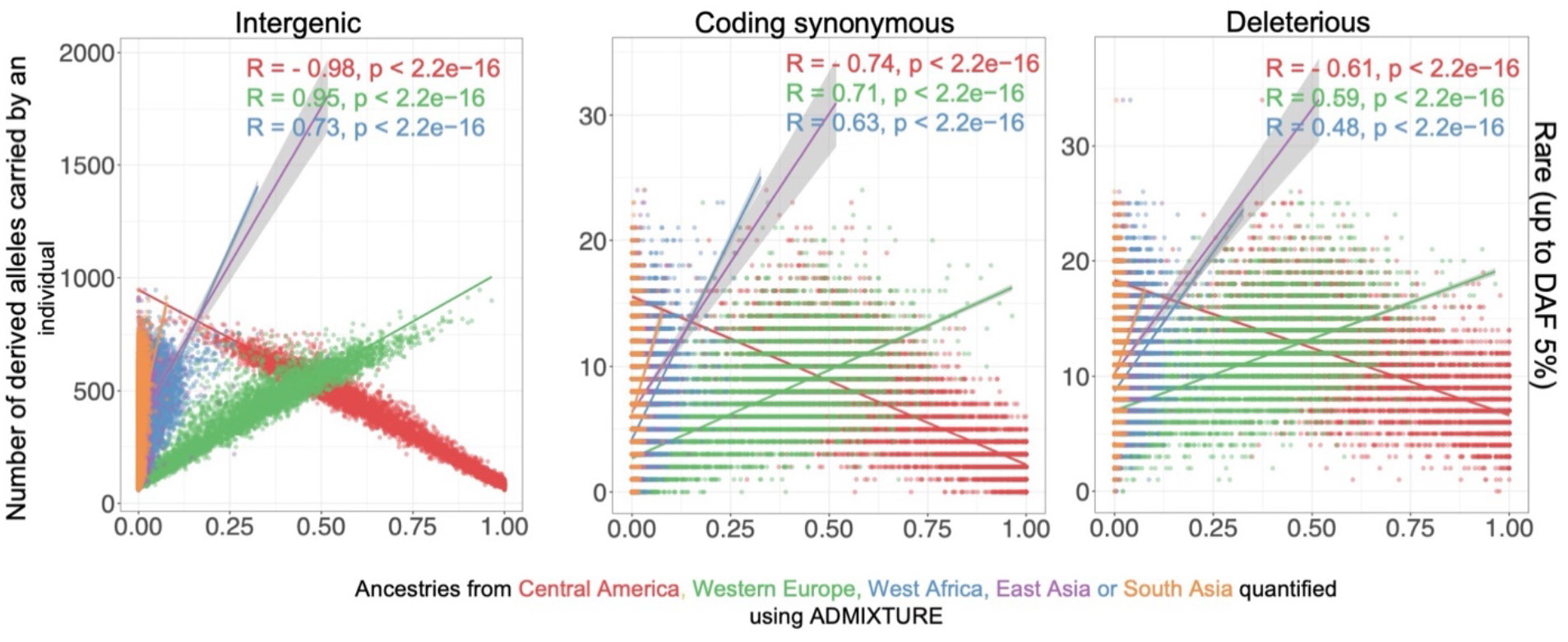
Mutation burden in different ancestries show effects of bottleneck in causing loss of rare variants. Rare variants are correlated with levels of ancestries from Central America, Western Europe or West Africa for rare variants (DAF < 5%), and common variants. Analysis of WGS from 1000 genomes MXL shows that the rare mutation burden result is robust while the full mutation burden correlation is caused by ascertainment bias of the MEGA array (Fig. S50). Variants were annotated using VEP, and deleterious variants are a combined set of missense variants predicted to be damaging by polyphen2 along with splice, stoploss and stopgain variants.

As shown in previous studies, the effect of demographic history on the total mutation burden is minimal in MXB^35–37^. When we consider all frequencies (DAF <= 100%), we see only a small correlation between mutation burden and ancestries from different regions likely due to ascertainment bias as this correlation does not persist in whole genome sequence data from 1000 Genomes (Fig. 4, Fig. S50). This is because while some rare variants are lost, some increase in frequency, compensating the total mutation burden, which overall remains unchanged.

## Complex traits display varying roles of genetics and environment

Lastly, we assessed the contribution of genetic variation towards impacting variation in complex traits or disease in Mexico (Fig. S32). For example, ROH have been previously shown to have associations with a broad range of complex traits, and estimated to be negatively associated with height, weight, and cholesterol^32^. Such associations, if due to genetic factors, point towards a recessive architecture of the traits. Further, genetic ancestry proxies can also be associated with complex traits due to genetic factors, or due to differential experience of discrimination and other socioeconomic factors (Fig. S32). Such genetic factors can be different distributions of ROH or other differential patterns of genetic variation caused by demographic and environmental histories that vary among ancestries, and that can lead to the association of particular causal genotypes with ancestry proxies. Indeed, genetic ancestry proxies in Mexico are correlated with the number and length of ROH (Fig. 3A). We therefore model the association of genetic factors such as ancestry proxies and ROH with trait variation in the same model to disentangle these effects. As genetic ancestry proxies can also reflect differential environmental exposures, we consider in our model several environmental factors, to further disentangle the role of genetic factors reflected in ancestry proxies compared to environmental factors. Our model therefore also includes variables available in the MXB related to discrimination, socioeconomic opportunities, and living environment (collectively called sociocultural and biogeographical factors). We also account for cryptic relatedness and potential unmodelled environmental factors using a genetic relationship matrix and city or town of origin as random effects in a mixed model framework. Significant associations would give insight into the architecture, including the genetic architecture, of the trait in Mexico to help guide future efforts in genetic mapping, and in considering other interventions towards improving public health.

Aiming to first understand how the traits are distributed geographically and relative to single model covariates, we first visualize average trait values by units of our biogeographical and sociocultural factors to understand the dimensions of trait variation (Figs. 5A, S33-41). Next, we use a mixed model to estimate the contribution of genetic factors to trait variation jointly modelled with the environmental factors (Figs. 5B, D-F, S42-49). Finally, we visualize trait values by birth year to assess their changes over time. We focus on several quantitative traits: height, BMI, triglycerides, cholesterol, glucose, blood pressure, and others. Our test predictive variables in the full model include genetic factors (genetic ancestry proxies from ADMIXTURE and ancestry-specific MDS analyses, and ROH in kb in each genome), life history factors (age, sex), sociocultural factors (educational attainment as a proxy for income levels (Fig. S42), whether they speak an indigenous language or not as a proxy for differential experience of discrimination/other cultural factors such as diet, whether they live in an urban or rural environment), and biogeographical factors (altitude, latitude and longitude). For height and other traits analyzed, a significant association with ancestry proxies could reflect the association of particular causal genotypes with those ancestries or associated unmodelled environmental factors such as nutrition. If the association is due to genetic factors, it should still not be interpreted as deterministic of a trait value, but makes a case for inclusion of diverse individuals in genetic studies of complex traits.

**Figure 5.**
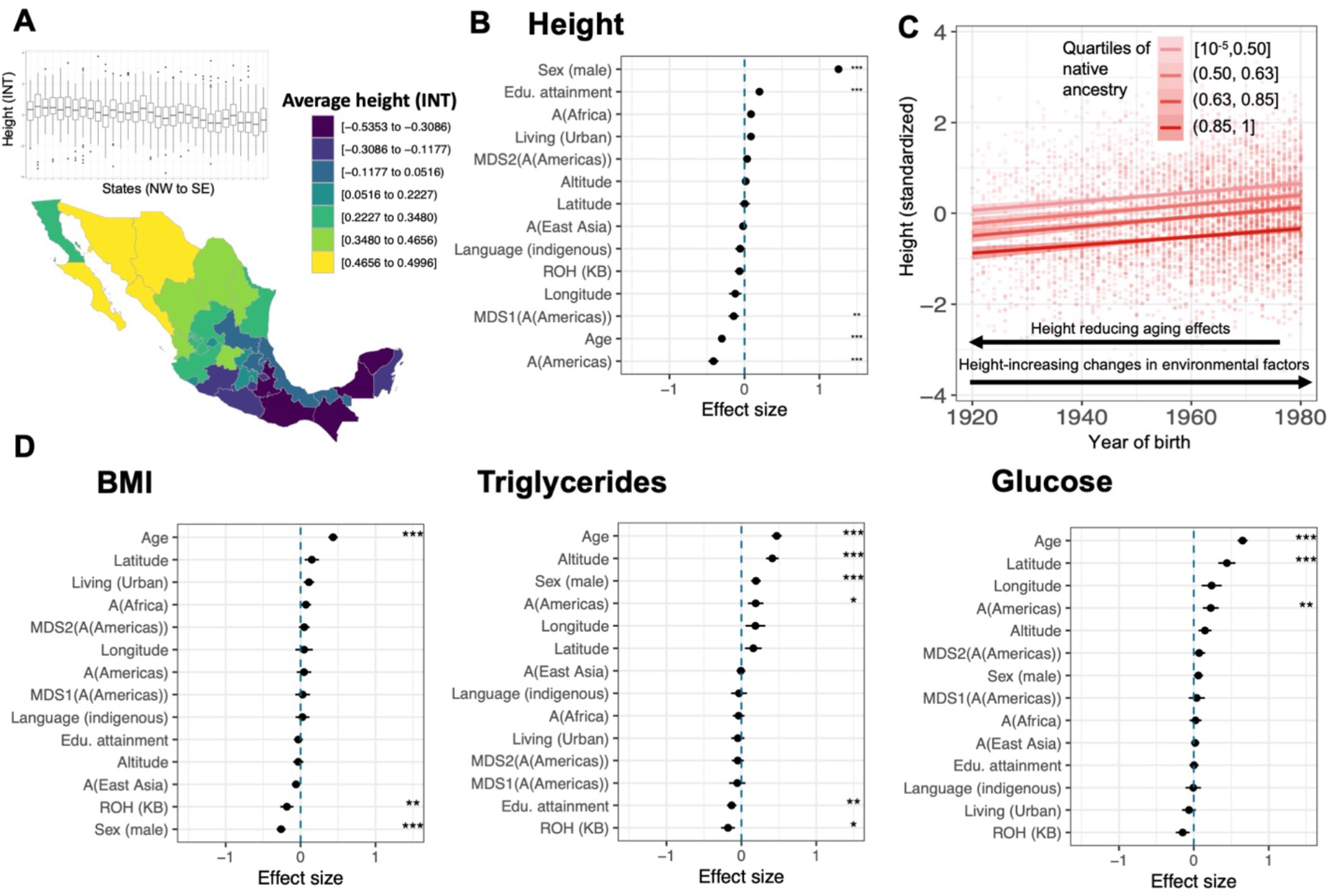
Height variation over space and birth year, and analysis of the factors influencing height and other complex trait variation. A) Map of average height in Mexico (inset shows boxplots of height variation in each state from North-west to South-east). B) Explanatory model for height variation implicates the role of genetics and environment. The plot shows effect size estimates and confidence intervals from a mixed model analysis. All quantitative predictors are centered and scaled by 2 standard deviations. Asterisks indicate significance of the effect of a predictor after Bonferroni correction (** P < 10^-5^, *** P < 10^-6^) across traits and predictors analyzed. C) Height as a function of birth year in quantiles of ancestries from Central America. D) Trait profiles for BMI, Triglycerides, and Glucose. Results of mixed model analysis same as in (B). Educational attainment is on a scale from 0-8 (low to high educational attainment), and altitude is measured in meters (low to high).

### Height

When viewed as averages per state, height values show a clear increasing pattern from southeast to northwest in the MXB (Fig. 5A). Even though every state shows a large variance (Fig. 5A), height shows a significant correlation with both latitude and longitude univariately (Figs. S34-35). With our explanatory mixed model, we can explain 66.33% of the variance for height. We find that individuals with higher ancestries from Central America are significantly shorter (*β* = −0.42, *p* < 2.2 × 10^-16^) (Figure 5b). Further, considering ancestries at a finer resolution, we observe decreased height with a change in ancestries from the North of Mexico (Huichol, Tarahumara) to the Mayan region (Tojolabal, Maya) (Fig. S48). Notably, individuals having higher educational attainment are also estimated to be taller. After Bonferroni correction across traits and predictors, the relationship between ROH and height is not statistically significant (*β* = −0.07,*p* = 0.03). Nevertheless, younger individuals with any range of ancestries from Central America are taller than older individuals with the same ancestries (Fig. 5C). As the positive correlation between birth year and height for all individuals regardless of their ancestries demonstrates, height can also vary due to environmental factors or aging.

### Body mass index (BMI)

BMI similarly shows a significant correlation with latitude and longitude univariately (Figs. S34-35). Our full mixed model explains 30.10% of the variation in BMI. In the full model, ROH remain significantly associated with lower BMI (*β* = −0.18, *p*= 3.95 × 10^-5^), while ancestry does not (Fig. 5D). BMI is also significantly correlated with birth year, increasing with older age, as well as with being female (Fig. 5D).

### Triglycerides

There are clear differences in levels by region that are more striking for cholesterol (see below) than for triglycerides (Fig. S33). Our full model explains 37.23% of the variation in triglyceride levels. Ancestries from Central America are significantly associated with higher triglyceride levels (*β* = 0.19, *p* = 3.17 × 10^-4^) while the ROH carried by an individual are significantly associated with lower triglyceride levels (*β* = −0.18, *p* = 1.4 × 10^-4^) (Fig. 5D). Notably, age, being male, lower educational attainment and high altitude are also associated with higher triglyceride levels (Fig. 5D).

### Cholesterol

Cholesterol levels show a significant correlation with latitude and longitude univariately (Fig. S33-35). In the full model (explaining 29.49% of trait variation), we do not see any correlation with genetic ancestry proxies or ROH, but we estimate significantly lower cholesterol in individuals who speak an indigenous language (*β* = −0.26,*p* = 6.93 × 10^-7^). We also estimate higher cholesterol in those living in an urban environment, at high altitude or of a higher age (Fig. S43). For HDL and LDL levels, we similarly find a significantly lower cholesterol in those who speak an indigenous language but not related to ancestry (Fig. S43) likely indicating that cultural/diet factors are stronger than the genetic factors tested here. Living in in an urban environment is also significantly associated with high HDL and LDL levels (Fig. S43).

### Glucose

Glucose levels show a significant correlation with latitude in univariate analysis (Fig. S34). In the full model (explaining 28.62% of the trait variation), we estimate ancestries from Central America to be significantly correlated with higher glucose levels (*β* = 0.23, *p* = 2.452 × 10^-5^). Glucose also remains significantly associated with latitude (increasing northwards) and higher age (Figure 5d). For fasting glucose, where we reduce sample size by about four-fifths, ancestry from Central America still has a positive estimated coefficient (Fig. S45), but it is not significant with the smaller sample size.

### Other traits

We also analyze creatinine, and systolic and diastolic blood pressure (Fig. S44). Individuals that speak an indigenous language have significantly lower creatinine (*β* = −0.18, *p* = 1.32 × 10^-3^). Living in an urban environment and altitude are significantly associated with higher creatinine. Only age and sex are significantly associated with diastolic and systolic blood pressure (adjusted by medication status) (Fig. S44).

Overall, ancestries from Central America are significantly associated with trait variation for height, triglycerides, and glucose levels (Fig. S46), the ROH present in a genome with BMI and triglycerides (Fig. S47) levels, and native ancestry variation within Central America with height variation (Fig. S48-49). In contrast, cholesterol levels, creatinine levels, and blood pressure are significantly associated with environmental, but not genome-wide genetic factors. This does not rule out the effect that specific gene variants may have in the variation of these traits, as illustrated before by candidate gene approaches discovering functional variants exclusive to the Americas^38^.

## Conclusion

Our work is a demonstration of the value of generating genotype-phenotype data on underrepresented populations to reveal lesser-known genetic histories and generate findings of biomedical relevance. It is also an illustration of the joint modeling of genetic and environmental effects to reveal the etiology of complex traits and disease. In this project, we ensured diverse indigenous and rural presence in our sampling strategy, considered the fluidity of ancestries from different local and global regions in our analyses and evaluated their reflection in genetic and disease-relevant complex trait variation. By leveraging the largest nationwide genomic biobank in Mexico, we find diverse sources of ancestries in Mexico in light of its unique history, infer population size changes and runs of homozygosity using ancestry-specific haplotype identity that reveal an elaborate fine-scale structure in the country. We also show that demographic history affects the frequency distribution of genetic variants thus changing how many rare variants individuals with different ancestries carry. We observe a significant impact of genetic ancestries, ROH, as well as socioeconomic and biogeographic variables on a variety of complex traits implicating the importance of both genetic and environmental factors in explaining complex trait variation and in considerations of potential public health interventions. Our results can inform the design of future studies in other admixed populations and the MXB will hopefully motivate additional efforts to strengthen local research capacity across Latin America and benefit underserved populations globally.

## Funding

This work was supported by “The Mexican Biobank Project: Building Capacity for Big Data Science in Medical Genomics in Admixed Populations”, a binational initiative between Mexico and the UK co-funded equally by CONACYT (Grant number FONCICYT/50/2016), and The Newton Fund through The Medical Research Council (Grant number MR/N028937/1) awarded to AME and AH. MS was also supported by the Chicago Fellows program of the University of Chicago. Training activities in Mexico were hosted by CINVESTAV and supported in part by CABANA, a capacity strengthening project for bioinformatics in Latin America, funded by the Global Challenges Research Fund (GCRF) of the UK.

## Data availability

The dataset for the 6,057 newly genotyped individuals from the MX biobank project are available at the European Genome-phenome Archive (EGA) through a Data Access Agreement with the Data Access Committee (EGA accession number in process).

## Code availability

All custom scripts/approaches described in Methods will be made available from https://github.com/msohail88/MXB_popstruct_complextraits on publication and all existing software packages and versions used are noted in Methods

## Acknowledgments

We thank the participants of the *Encuesta Nacional de Salud, 2000* (2000 National Health Survey, ENSA 2000), conducted in Mexico nationwide by the *Secretaría de Salud (*Health Secretariat) and the *Instituto Nacional de Salud Pública* (National Institute of Public Health, INSP). We are grateful to Mauricio Hernández and Celia Alpuche-Aranda for Institutional support from INSP, Mitzi Flores, Rocío Nájera, and Adriana Garmendia for project management support, and to Carlos Conde, Victor Guerrero Lemus, Armando Mendez Herrera, Cruz Portugal García, Rosario Rodriguez, and Manuel Velazquez Mesa for biobank maintenance and sample preparation. We thank Mary Ortega, Cecilia Gutiérrez, and Sara García for technical assistance,

Jacob Cervantes for IT support, Harald Ringbauer for useful advice with the ROH analysis, Juan Esteban Rodriguez for helpful conversations about population structure in Mexico, and Arslan Zaidi for useful comments on an earlier draft of this manuscript.

## Inclusion and Ethics statement

Samples were collected as part of the 2000 National Health Survey (ENSA 2000) conducted by the National Institute of Public Health (INSP), and informed consent was obtained from all participants. The ENSA 2000 was carried out following the strictest ethical principles and in accordance with the Helsinki Declaration of Human Studies. Extracted DNA has been stored and maintained at the National Institute of Public Health (Cuernavaca, Mexico), and samples were genotyped at the Advanced Genomics Unit of CINVESTAV (Irapuato, Mexico) through a collaboration agreement. The data has been jointly analyzed promoting local leadership and participation of Mexican researchers and trainees. The project was reviewed and approved by the Research Ethics Committee and the Biosafety Committee of the National Institute of Public Health (IRB approvals CI: 1479 and CB: 1470). For the present project, personally identifiable data was removed from the data set.

## Notes

### Competing Interest Statement

The authors have declared no competing interest.

### Summary of Updates

Supplemental file updated

