## Supplementary Material for "Nationwide genomic biobank in Mexico unravels demographic history and complex trait architecture from 6,057 individuals"

### List of Supplementary Figures

**Fig. S1. Map of Mexico with state labels.**
**Fig. S2. Number of individuals per state.**
**Fig. S3. Distribution of rural and urban samples in the MXB.**
**Fig. S4. Number of MXB individuals that speak Spanish, and indigenous language or both.**
**Fig. S5. Number of MXB individuals from a rural or urban locality.**
**Fig. S6. PCA analysis of MXB with global reference panels.**
**Fig. S7. PCA analysis of MXB.**
**Fig. S8. PCA analysis of MXB and NMDP grouped by Mesoamerican regions.**
**Fig. S9. PCA analysis of MXB and NMDP grouped by indigenous culture.**
**Fig. S10. Admixture analysis of MXB in principal component space of MXB and NMDP.**
**Fig. S11. Admixture analysis of MXB and NMDP.**
**Fig. S12. UMAP analysis of MXB samples with 10 principal components.**
**Fig. S13. UMAP analysis of MXB samples with global reference panels.**
**Fig. S14. Fst analysis of autosomes of MXB individuals.**
**Fig. S15. Fst analysis of X chromosomes of MXB individuals.**
**Fig. S16. Fst analysis of autosomes of MXB individuals with high ancestries from Central America.**
**Fig. S17. Fst analysis of X chromosomes of MXB individuals with high ancestries from Central America.**
**Fig. S18. Origins of ancestries from the Central America in MXB.**
**Fig. S19. Origins of ancestries from West Africa in MXB.**
**Fig. S20. Origins of ancestries from East Asia in MXB.**
**Fig. S21. Origins of ancestries from South Asia in MXB for all states.**
**Fig. S22. Origins of ancestries from South Asia in MXB for Guerrero.**
**Fig. S23. asIBNe analysis results for ancestries from Central America are visualized for 30 or 20 years per generation.**
**Fig. S24. Estimated effective population size for ancestries from Western Europe by Mesoamerican region in Mexico (30 years per generation).**
**Fig. S25. Estimated effective population size for ancestries from Western Europe by Mesoamerican region in Mexico (20 years per generation).**
**Fig. S26. Estimated effective population size for ancestries from West Africa by Mesoamerican region in Mexico (30 years per generation).**
**Fig. S27. Estimated effective population size for ancestries from West Africa by Mesoamerican region in Mexico (20 years per generation).**
**Fig. S28. NROH distribution by state.**
**Fig. S29. SROH distribution by state.**
**Fig. S30. NROH are correlated with ancestries in MXB.**
**Fig. S31. SROH are correlated with ancestries in MXB.**
**Fig. S32. A framework for the role of genetics and environment in creating genetic variation and variation in complex traits and disease risk used in this study.**
**Fig. S33. Average trait values per state visualized on the map of Mexico.**
**Fig. S34. Complex trait variation by latitude.**
**Fig. S35. Complex trait variation by longitude.**
**Fig. S36. Complex trait variation by altitude.**

**Fig. S37. Complex trait variation by sex.**
**Fig. S38. Complex trait variation by birth year.**
**Fig. S39. Complex trait variation by indigenous heritage.**
**Fig. S40. Complex trait variation by rural or urban living environment.**
**Fig. S41. Variograms of complex traits.**
**Fig. S42. Correlation of educational attainment and income levels.**
**Fig. S43. Mixed model results for Cholesterol related traits.**
**Fig. S44. Mixed model results for Creatinine and blood pressure.**
**Fig. S45. Mixed model results for fasting glucose.**
**Fig. S46. Mixed model results for ancestries from Central America.**
**Fig. S47. Mixed model results for sROH carried by an individual.**
**Fig. S48. Mixed model results for two MDS axes reflecting genetic variation within Central America.**
**Fig. S49. Mixed model results for two MDS axes reflecting genetic variation within Central America (using MAAS-MDS run on MXB and NMDP excluding Tojolabal individuals).**
**Fig. S50. Mutation burden in 1000G MXL individuals as a function of their ancestries from the Central America, Western Europe, and West Africa.**
**Fig. S51. Ancestries from Central America as a function of birth year in rural and urban localities.**
**Fig. S52. Proportion of individuals that speak Spanish, an indigenous language or both, as a function of birth year.**

**List of Supplementary tables**

**Table S1. Age and sex distribution in the Mexican Biobank.**

**Table S2. List of anthropometric, disease and lifestyle variables in the Mexican Biobank**

**Table S3. Estimation of admixture proportions by state in MXB.**

**Table S4. NMDP cultural groups and their designation into Mesoamerican regions.**

**Table S5. Number of ROH by local ancestry tracts.**

### Supplementary Text

#### Strengths and limitations of MXB

We summarize the strengths and limitations of the the MXB. Strengths include:1) large sample size; 2) the initial study population was representative of the Mexican civilian population at state and national levels in 2000; 3) we purposefully over selected indigenous populations 4) collection of socio-demographic, geographic, anthropometric, clinical, biochemical, and genetic data that allow for adjustment of models; 5) some of the clinical and biochemical markers have become public health problems in the present day (diabetes, obesity)[1] and thus this study may contribute to better understanding of the epidemiology and causal factors; 6) representativeness of the data set allows correlation with diseases that were unknown at the time of the survey and that have now been characterized (e.g. COVID-19). Limitations include: 1) the study is cross-sectional, thus limiting causal inferences (e.g. age and glucose); 2) even though the survey was conducted in 2000, it is unlikely that the genetic composition of the Mexican population has varied much;  3) the sample size limits detection of phenotypes that are rare; 4) there are many characteristics, factors, or variables that were not studied or are unknown that undoubtedly modify the significance of our observations; however, we are able to estimate the extent to which our models account for the variation in the observed data.

#### Gene-culture co-evolution in MXB for ancestries and languages from the Americas

We observe in the MXB that ancestries from the Americas are more frequently observed in younger individuals than in older individuals in the biobank (Fig. S51), and that this signal is largest in rural areas. We also observe that the proportion of individuals that speak an indigenous language is lower in younger individuals compared to older individuals, providing an example of anti-correlated gene-culture co-evolution (Fig. S52). Many linguists, artists, and human rights activists have been warning that Mexico is becoming increasingly monotone, and this is not because of the lack of indigenous presence today, but due to discrimination and repressive assimilation policies, whereby many indigenous people believe it is better not to be heard speaking their native language in Mexico[2]. Now, geneticists such as ourselves join this chorus as we observe an increasing presence of ancestry from the Americas going hand in hand with a decline in the passage of indigenous language.

### Materials and Methods

#### ENSA2000

Since 1987, Mexico has established periodical health surveys (ENSA) for surveillance of Mexican population-based health metrics. In this study, we use data and samples collected from the survey performed in 2000, the ENSA 2000. This survey was a probabilistic, multi-stage, stratified, cluster household survey conducted by the Mexican Secretariat of Health from November 1999 to June 2000. Research design and methods have been described elsewhere[3]. Participants were randomly selected in order to be representative of the civilian, non-institutionalized Mexican population, at the state and national levels. Trained personnel conducted the interviews. Information was collected on household and sociodemographic characteristics, current health status, health care service usage, and behavioral aspects of participants. Sera and buffy coats were obtained from 43,085 individuals aged 20 years or older. More than fifty publications have arisen from this survey providing critical insights into the status of national health alongside some genetic traits of the sampled population[4]. In particular, the inclusion of individuals from remote and rural locations in Mexico makes this survey unique. Given its large volume, sophisticated sampling design, breadth of demographic sampling and extensive trait data, the ENSA 2000 represents a valuable untapped genetic resource to link genetic markers and health outcomes.

#### The Mexican Biobank Project: phenotype, lifestyle and environmental data

For each individual, we have access to a range of anthropometric, disease, lifestyle and environmental information. These variables are summarized in Table S2. Serum samples were further used to measure a number of biochemical traits analyzed in this study. All traits analyzed in the complex trait analysis were pre-processed as follows.

Biometric data were filtered to remove outliers with apparent errors in data entry. Outliers were identified based on distribution density over the complete 6k dataset, resulting in height (between 100-200 cm) and weight (between 25-300 kg).

Biochemical traits were similarly curated to remove extremes and negative values (<0). Glucose was also checked against finger prick tests taken at time of survey and values that were greatly discordant were also removed. Glucose measurements were further stratified by random or fasting glucose samples based on participant questionnaire responses.

Blood pressure was manually curated for individuals where values differed by more than 20 units for the two readings taken, where diastolic pressure was higher than systolic, or were unusually high/low (<30 or >300). In these cases, both readings were manually checked, and discordant readings discarded. These updated values were then merged with the remaining samples. A set of adjusted blood pressure phenotypes was also generated, adjusting for treatment for hypertension. In those individuals who were reported to be receiving some form of hypertension treatment, 15 units were added to SBP and 10 to DBP (SBP_adj and DBJ_adj)[5,6].

Quantitative traits were inverse normalized and transformed traits were used for the complex trait analyses.

For each individual, we have access to variables for various socio-cultural factors such as access to healthcare and clean water, yearly income, educational attainment, whether they speak an indigenous language or not, and whether they live in a rural or urban environment.

Localities were assigned values of latitude, longitude, and altitude (meters) by the National Institute of Statistics and Geography (INEGI) in Mexico.

**The Mexican Biobank Project: genetic data**

Our samples were genotyped on the Illumina's Expanded Multi-Ethnic Genotyping Array (MEGAex). The design of this array was previously led by Christopher Gignoux and Genevieve Wojcik, co-authors on this manuscript. Several properties place the MEGA array as the ideal choice for biobank genotyping. It captures 1,748,250 SNPs derived from admixed population studies, making it broadly applicable in diverse populations. The array has boosted SNP coverage in both the MHC and KIR loci, a marker set of over 30,000 SNPs for ancestry estimation, and the inclusion of over 17,000 medically relevant genetic variants from prior GWAS and clinical studies. Such breadth of coverage of genomic diversity provides a comprehensive quantitative resource of the genetic variability in this cohort.

**Generation and quality control of MXB genetic data**

Genome Studio was used to convert raw image files to plink files with raw genotype information. All SNPs were flipped to the forward strand, and duplicate SNPs were removed. For sites with missing chromosome number, physical position or both, we updated the map using the information in the SNP name or by mapping their rsID using dbSNP Build 151.

We removed all individuals with more than 5% missing genotype data, and all genotypes with more than 5% missing individuals. We restricted the analyses to only autosomes and removed all monomorphic SNPs. We restricted the analysis to only biallelic SNPs and removed all SNPs with ambiguous strand for all downstream analyses. All related individuals were detected using plink (--Z-genome --min 0.5) after pruning for LD (--indep-pairwise 50 5 0.5). A script was written to iteratively find nodes and remove related individuals to obtain the final qc-ed dataset.

**Reference panels: sources and quality control**

Reference genetic panels were used for various analyses of population structure. We used global populations from the 1000 Genomes Project[7] and Human Genome Diversity Project[8], Zapotec individuals from Oaxaca from the PAGE study[9], and native individuals from across Mexico from the NMDP[10] for the analyses of population structure and ancestry.

For each reference panel, we restricted to autosomes, removed all monomorphic SNPs, flipped all SNPs to the forward strand, and removed SNPs with ambiguous strand.

**Anthropological classification**

We used anthropological and archaeological context to delineate different Mesoamerican regions[11]. An individual's locality was used to place them into one of the seven regions: the north of Mexico, the north of Mesoamerica, the center, occident and gulf of Mexico, Oaxaca and the Mayan region in the south east[11]. This classification was used to visualize and regionalize some of the population structure and history analyses, especially those relating to native genetic substructure within Mexico.

#### Note on genetic ancestry

Genetic ancestry represents a path through the ancestral recombination graph[12]. In this study, we obtain proxies for the ARG using ADMIXTURE[13] (see below). As such, we are discretizing a continuous quantity for the purposes of understanding the effects of varying demographic histories on genetic and complex trait variation in MXB. Labelling and use of such discretized ancestry proxies remains a contentious issue[14]. To clarify the point that such proxies are not essentialized entities in the real world, but rather variables we use for the purposes just described, we opt to refer to our ancestry proxies as from the region whose present-day individuals such proxies cluster with. Thus, we use, “ancestry from the Americas”, “ancestry from Europe,” “ancestry from Africa,” “ancestry from South Asia,” and “ancestry from East Asia” in the text, and shorter versions of the same for some figures (A(Americas) etc.).

Such largely continental regions were useful for our analyses only in so much as they reflect demographic and environmental histories, the implications of which on genetic and complex trait variation we are interested in. This is only one arbitrary scale to discretize at, and we also consider origins of and implications of ancestral variations within such regional groupings in several analyses, where we perform dimensionality reduction within such regional groupings (For example, PCA(A(Americas)), MDS(A(Americas)) etc.).

We would like to note that continental groupings are similar to racial categories that were created in the last 500 years and used to justify European superiority and colonization of global regions including present-day Mexico[1415,16]. In Mexico, such categories have a similar history of racism and eugenics as in other parts of the world[17]. Aware of this history, we still believe it of scientific and medical value to assess the patterns of genetic and complex trait variation that result from varying demographic and environmental histories at different scales. We reject continental groupings as a fixed hierarchical categorization of humans, as well as their use to justify the superiority of one group over another. Instead, we use ancestry proxies that are estimated from Admixture using an unsupervised clustering, as well as axes of ancestry that result from dimensionality reduction within these ancestries, capturing variation among groups from the Americas for example. This is to highlight that genetic ancestry in humans is continuous over time and space and should only be considered in that complexity and at different scales.

#### Population structure analyses

For the analyses of population structure, we merged the qc-ed MXB dataset and reference panels using plink. We repeated some of the QC steps on the merged dataset, removing any monomorphic or duplicate SNPs. We also removed individuals with more than 5% missing genotype data, and genotypes with more than 5% missing individuals to obtain the clean merged dataset.

We performed two sets of analyses using principal components (PCA), admixture and Uniform Manifold Approximation and Projection (UMAP). One was performed on the merged dataset including MXB, Zapotecs from the PAGE study and global populations from 1000 Genomes Project and HGDP, and a second analysis was performed on the merged dataset including only

MXB and individuals native to present-day Mexico from the NMDP. Fst analysis was performed on all MXB individuals, as well as on only MXB individuals with 90% ancestry from the Americas as estimated from the admixture analysis.

Smartpca from Eigenstrat[18] was used to perform the principal components analysis. The R package pca3d was used to visualize the first three principal components. ADMIXTURE[13] was used to perform the admixture analysis. Principal components generated by smartpca were used to perform the UMAP analysis[19]. Fst analysis was performed using smartpca.

Given the large loss of SNPs due to admixture linkage disequilibrium in our admixed Mexican individuals, we opted not to prune for linkage disequilibrium for the population structure analyses presented in this study. We repeated the analysis on an LD pruned set of SNPs and obtained similar results (data not shown). Given the admixed nature of the Mexican individuals, we did not remove SNPs due to Hardy-Weinberg equilibrium in the MXB, as many SNPs are expected to be out of Hardy-Weinberg equilibrium due to admixture, and population structure.

**Analyses of sub-continental ancestry**

Analyses were performed to obtain axes of genetic variation or ancestry amongst each continental group. Such analyses also help place the specific origins of an ancestry present in Mexico today. These analyses were performed using rfmix[20] to estimate local ancestry along the genome and pcamask[21] to perform an ancestry-specific PCA for ancestries originating from present-day Africa and Asia. During the course of this study, new and improved methods to estimate local ancestry along the genome (GNOMIX)[22] and to perform ancestry-specific PCA (Multiple Array Ancestry Specific Multidimensional Scaling, MAAS-MDS, an MDS designed for analyzing samples from several different genotyping arrays simultaneously)[23] were published, allowing us to use these tools for the analysis within ancestry from the Americas the results of where were also used for the complex trait analysis.

maas-MDS on ancestry from the Americas. For the MAAS-MDS[23] analyses, local ancestry was inferred with Gnomix[22] using the same three continental references for each array. For European reference we used the populations IBS and GBR from the 1000 Genomes Project (1KGP)[7], for ancestry from Africa, the YRI population from 1KGP, and for ancestry from the Americas, PEL from 1KGP (only those samples with more than 95% ancestry from the Americas) and the 50 genomes of native individuals across Mexico generated as part of the Mexican Biobank Project[24]. The reference genomes were merged with each array resulting in 856,352 SNPs in array one and 967,338 in array two. Array one included eleven Native American groups from NMDP genotyped with the Affymetrix 6.0 array: Tarahumara, Huichol, Purepecha, Nahua, Totonac, Mazatec, Northern Zapotec (from Villa Alta District, Northern Sierra in Oaxaca State), Triqui, Tzotzil and Maya (from Quintana Roo State). Array two included the 6,051 individuals from the MX-Biobank project genotyped with MEGAex. The MAAS-MDS was applied to the Native American ancestry segments, that is, masking intercontinental components of African and European origin, in both arrays one and two. The analysis was run using average pairwise genetic distances and only considering individuals with >20% Native American ancestry, to generate ancestry specific MDS axes for ancestry from the Americas in the MXB.

asPCA on ancestry from Africa. We performed this analysis on all individuals in the MXB with >= 5% ancestry from Africa estimated from the admixture analysis. This resulted in 1965 individuals with ancestry originating from present-day Africa. In this set of individuals, we used populations from the 1000 genomes project (CEU, YRI)[7] and PAGE (Zapotecs from Oaxaca)[9] to estimate local ancestry using rfmix[20]. The MHC region was excluded from the analysis. SNPs out of Hardy-Weinberg equilibrium were removed from each of the reference panels (10^-3) and the MXB AFR (10^-8) subset beforehand. This dataset was merged with a subcontinental reference panel covering a range of groups in present-day Africa[25]. Pcamask[21] was used to mask all ancestries other than ancestry from Africa, and to generate ancestry specific principal components for ancestry from Africa in the MXB.

asPCA on ancestry from East Asia. We performed this analysis on the top 100 individuals in the MXB with ancestry from East Asia estimated from the admixture analysis. This ranged from 3.9% to 52% ancestry originating from present-day East Asia. In this set of individuals, we used populations from the 1000 genomes project (CEU, YRI, KHV, CHS)[7] and PAGE (Zapotecs from Oaxaca)[9] to estimate local ancestry using rfmix[20]. The MHC region was excluded from the analysis. SNPs out of Hardy-Weinberg equilibrium (10^-3) were removed from each of the reference panels and the MXB EAS subset beforehand. This dataset was merged with a subcontinental reference panel (including all 1000 Genomes East Asian populations and East Asian, Mainland Southeast Asian, and Southeast asian groups from the OGVP)[7]. Pcamask[21] was used to mask all ancestries other than ancestry from East Asia, and to generate ancestry specific principal components for ancestry from East Asia in the MXB.

asPCA on ancestry from South Asia. We performed this analysis on the top 100 individuals in the MXB with ancestry from South Asia estimated from the admixture analysis. This ranged from 2.5% to 7.7% ancestry originating from present-day East Asia. In this set of individuals, we used populations from the 1000 genomes project (CEU, YRI, GIH)[7] and PAGE (Zapotecs from Oaxaca)[9] to estimate local ancestry using rfmix[20]. The MHC region was excluded from the analysis. SNPs out of Hardy-Weinberg equilibrium (10^-3) were removed from each of the reference panels and the MXB SAS subset beforehand. This dataset was merged with a subcontinental reference panel (including all 1000 Genomes South Asian populations)[7]. Pcamask[21] was used to mask all ancestries other than ancestry from South Asia, and to generate ancestry specific principal components for ancestry from South Asia in the MXB.

**Population History analyses**

Analyses of population history using an approach that uses ancestry-specific IBD segments (asIBDNe) were performed on the entire MXB dataset, and on individuals belonging to each of the Mesoamerican regions. Identity-by-descent (IBD) segments of the genome can be used to estimate effective population size (Ne) for thousands of years into the past (IBDNe)[26]. These IBD segments can be further overlapped with local ancestry tracts to obtain ancestry-specific IBD tracts to estimate population size in an ancestry-specific manner for an admixed cohort (asIBDNe)[27].

For this analysis, the MXB was merged with populations from the 1000 genomes project (CEU, YRI)[7] and PAGE (Zapotecs from Oaxaca)[9] . SNPs in each population were previously filtered for Hardy-Weinberg equilibrium (10^-5 for reference groups and 10^-10 for the mxb samples).

The MHC region was excluded from the analysis. We repeated some of the QC steps on the merged dataset, removing any monomorphic or duplicate SNPs. We also removed individuals with more than 5% missing genotype data, and genotypes with more than 5% missing individuals to obtain the clean merged dataset.

We followed a computational pipeline recommended by the developers of asIBDNe to call ibd segments and local ancestry along the genome. We used beagle (beagle.25Nov19.28d.jar)[28] to phase the data, refined-ibd (refined-ibd.17Jan20.102.jar)[29] to call IBD, and merge-ibd-segments (merge-ibd-segments.17Jan20.102.jar) to remove breaks and short gaps in IBD segments, removing gaps between IBD segments that have at most one discordant homozygote and that are less than 0.6 cM in length. Local ancestry was estimated using rfmix. The RFMIX output was rephased to match the original phasing. asIBDNe (ibdne.19Sep19.268.jar) was run to estimate ancestry-specific population sizes using a 2 cM IBD length threshold.

**Runs of homozygosity**

The MXB dataset was LD-pruned using plink (--indep-pairwise 50 5 0.9). Runs of homozygosity were estimated using plink ( --homozyg) identifying 349400 ROH. We estimated nROH (number of ROH carried by an individual) and sROH (total sum of ROH in an individual in kb). ROH were divided into small, medium and large according to the theoretical framework in[30]. Python scripts were used to categorize ROH by length, and to overlap ROH with local ancestry calls from rfmix to obtain ancestry-specific ROH summary statistics. Local ancestry calls were the same as used for the asIBNe analysis. 38340 ROH did not overlap a homozygous local ancestry assignment and were removed from this analysis, the remaining 311060 that overlapped a homozygous local ancestry assignment were kept. We used a python script to compute the number of ROH in ancestry switch points as well (58 ROH or 0.00019 of all ROH fall within an ancestry switch and were also excluded from the analysis).
ROH were also correlated with birth year in the MXB and used as a variable in the complex traits mixed model analysis. For the birth year analysis, we removed the first two decades, as each year has below 15 individuals in this period. Birth year was also directly correlated with level of ancestry from the Americas inferred using admixture in rural and urban localities separately. ROH were also correlated with global ancestries per individual estimated from the admixture analysis. An R script was used to analyze distributions of sROH by geography.

**Mutation burden analyses**

Variants were annotated according to whether they were ancestral or derived, and their functional effect depending on their location in a gene or genome. Ancestral alleles for each SNP in the MXB were inferred using the EPO pipeline from the 1000 Genomes Project. Variant Effect Predictor (VEP)[31] was used to annotate the effect of a variant using the humdiv database, and picking one consequence (or transcript) per variant according to a criteria that includes the canonical status of the transcript, APPRIS isoform annotation, transcript support level, biotype of transcript ("protein_coding" preferred) and consequence rank preferring high impact.

Mutation burden is defined as the sum of derived alleles carried by an individual. A computational pipeline using vcftools, python, linux and R was used to compute mutation burden in different classes of variants, and at different derived allele frequency thresholds. We computed either a rare mutation burden (derived allele frequency <= 5%) or an overall mutation burden considering all allele frequencies. Our pipeline used R packages matrixStats, dplyr and ggplot2. We correlated the mutation burden with the global ancestry percentage from different present-day continental origins in all individuals. The ancestry estimates were from the admixture analysis. We computed a spearman's correlation and p-value.

This analysis was repeated in the 1000 Genomes Project MXL population. This was to check if the effect we were observing was due to ascertainment bias in the MEGAex array which covers fewer rare variants predominantly native to the area that is Mexico today. The whole genome sequences from the 1000G Project allowed us to rule this out. Ancestry estimates were generated using Admixture with reference panels from 1000 Genomes (CEU, GBR, YRI, PER) and 50 whole genomes sequences of native individuals across Mexico generated as part of the Mexican Biobank Project[24]. Variant effect predictor was used to annotate SNPs, and mutation burden was computed in the same manner. The deleterious category includes the following consequence terms: splice acceptor variant, splice donor variant, stop gained, stop lost, start lost, and transcript amplification.

**Complex trait analyses**

To assess the factors involved in creating complex trait variation, we performed a mixed model analysis using the lme4qtl R package. Lme4qtl allows flexible model creation with multiple random effects[32].

As fixed predictors of complex trait variation, we considered genetic variables. These included genetic ancestries either estimated from admixture or continuous axes of ancestry such as those estimated using maas-MDS. We also considered runs of homozygosity (amount of ROH carried in an individual genome in kb). We also considered biogeographical variables such as latitude, longitude and altitude (meters). We considered life history variables of age and sex. Lastly, we considered variables that are proxies for socio-economic and cultural differences (educational attainment as a proxy for income levels (Fig. S42), whether they speak an indigenous language or not as a proxy for differential experience of discrimination/culture, whether they live in an urban or rural environment). In order to ease interpretation of the mixed model coefficients for jointly considered numeric and binary predictors, we standardized predictor variables as follows[33]. To make coefficients of numeric predictors comparable to those for untransformed binary predictors, we divide each numeric variable by two times its standard deviation[33]. We centered both the binary and numeric predictors. All the covariates mentioned above are significant when jointly modeled for at least one tested trait, justifying their use in the full model.

We also include two random predictors in our model. These are (1) the covariance structure defined by the genetic relationship matrix (GRM) and (2) the locality where the individual is from to capture any other environmental variation (such as diet) not captured by the fixed predictors.

The GRM was generated using the GENESIS R package using kinship coefficients. As kinship estimates can be inflated under the presence of population structure and admixture, we obtained kinship coefficients for the GRM in the following manner: (1) PC-air[34] was used to obtain principal components that capture ancestry and not relatedness. This procedure used kinship coefficients estimated using KING[35] as input to partition samples into a related (5,562) and unrelated (271) set (using kinship threshold 0.044) and performing PCA on the unrelated set (2) PC-relate[36] was used to obtain kinship coefficients that capture relatedness but not ancestry. This method uses the ancestry-representative principal components from (1) to correct for population structure before calculating the kinship coefficients.

For this analysis, we removed rare variants (MAF < 5 %), regions with known long-range linkage disequilibrium[37] and variants in high linkage disequilibrium (r2> 0.1 in a window of 50 kb and a sliding window of 1 variant).

Bonferroni correction across all traits and predictors analyzed was used to assess the significant effect of a predictor variable on a complex trait.

Maps of Mexico to visualize trait distributions were created using the mxmaps R package.

**Links to software and R packages used:**

EIGENSTRAT: https://github.com/DReichLab/EIG
ADMIXTURE: https://dalexander.github.io/admixture/
UMAP:https://github.com/lmcinnes/umap
GNOMIX: https://github.com/AI-sandbox/gnomix
maas-MDS: https://github.com/AI-sandbox/maasMDS/
RFMIX: https://github.com/slowkoni/rfmix
PCAmask: https://sites.google.com/site/pcamask/home
asIBDNe: https://faculty.washington.edu/sguy/asibdne/
Plink: https://www.cog-genomics.org/plink/1.9/
Variant Effect Predictor: https://www.ensembl.org/info/docs/tools/vep/index.html
KING: https://www.kingrelatedness.com/
Mxmaps: https://www.diegovalle.net/mxmaps/
Genesis: http://bioconductor.org/packages/release/bioc/html/GENESIS.html
lme4qtl: https://github.com/variani/lme4qtl

### Supplementary figures

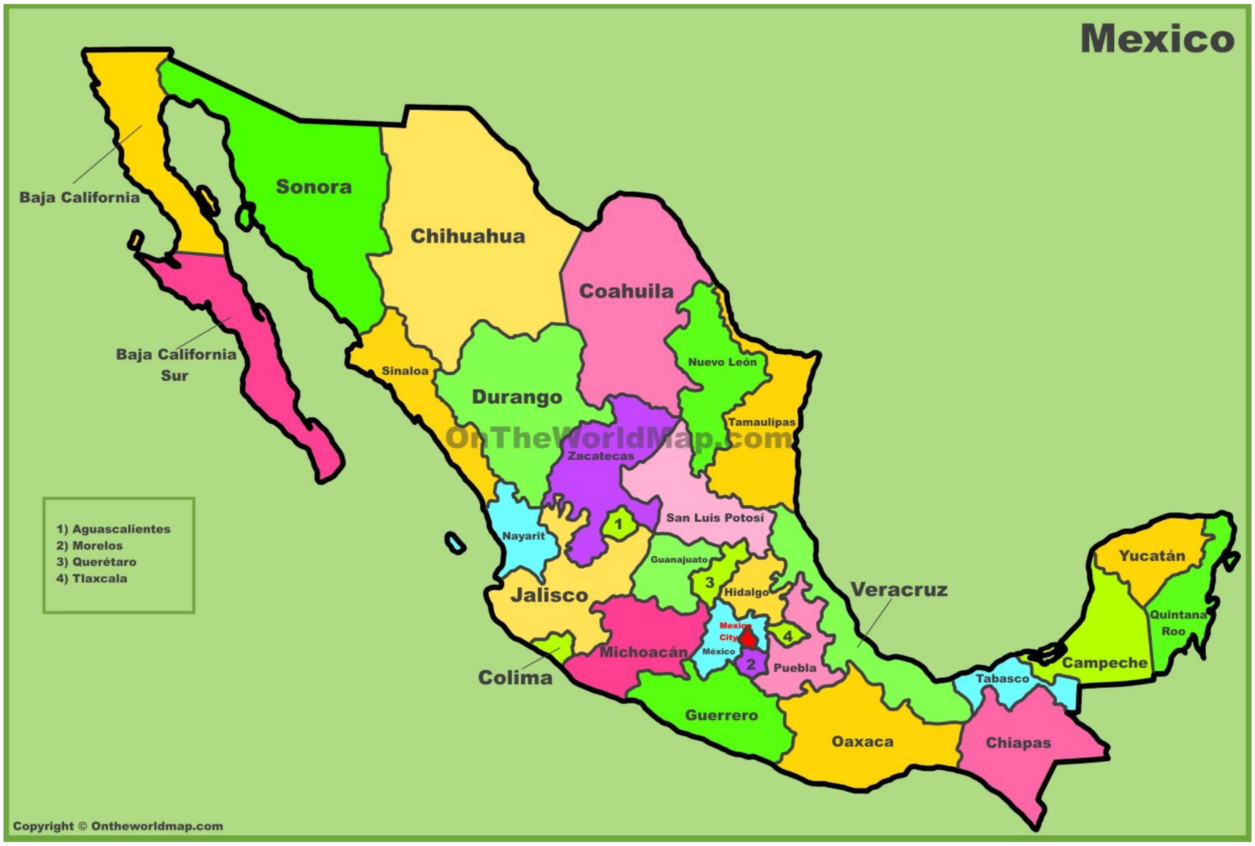


**Fig. S1. Map of Mexico with state labels.** The MXB project collected and genotyped samples from all 32 states of Mexico. Reproduced with permission from https://ontheworldmap.com/mexico/mexico-states-map.html.

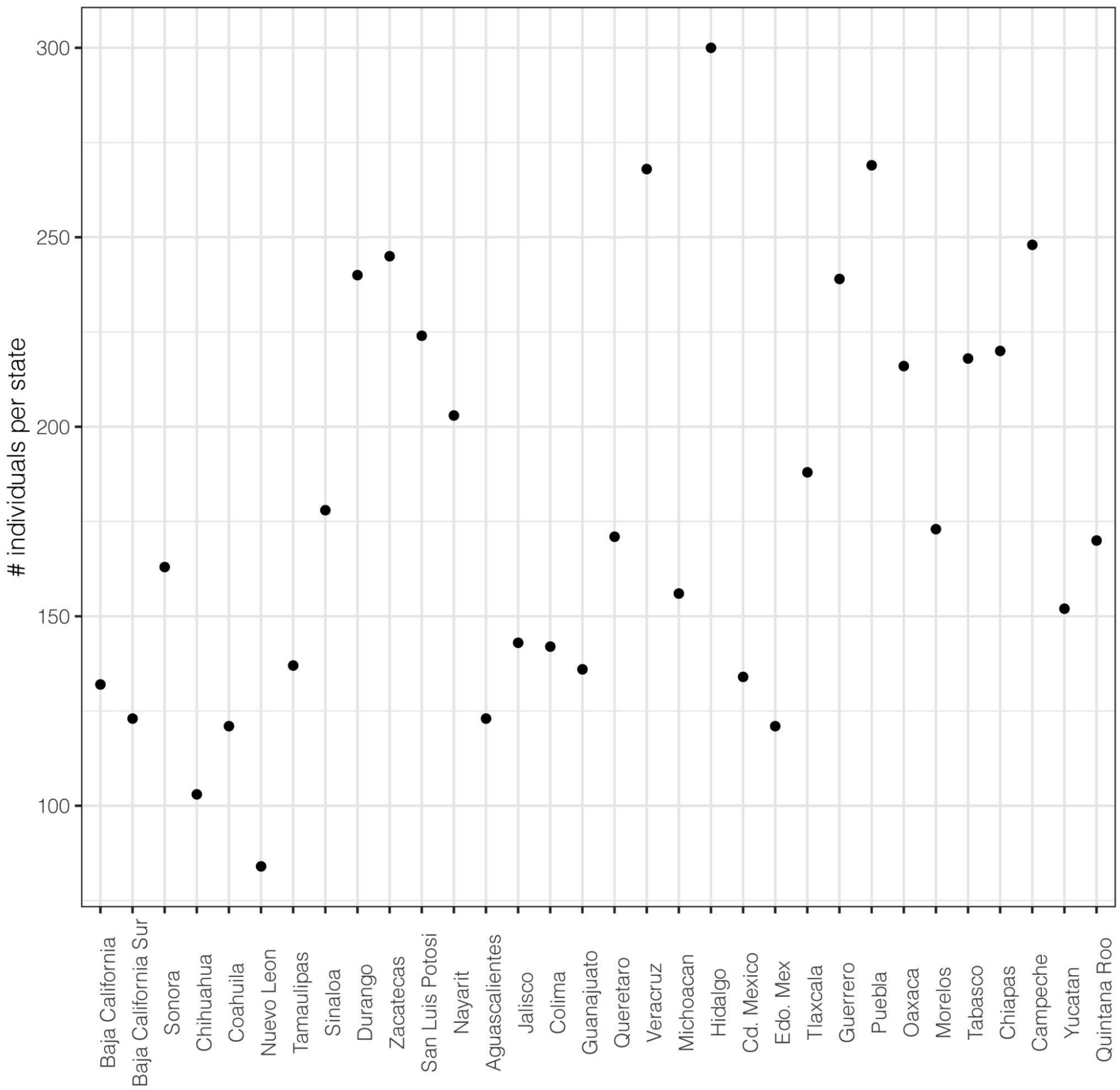


**Fig. S2. Number of individuals per state in the MXB.**

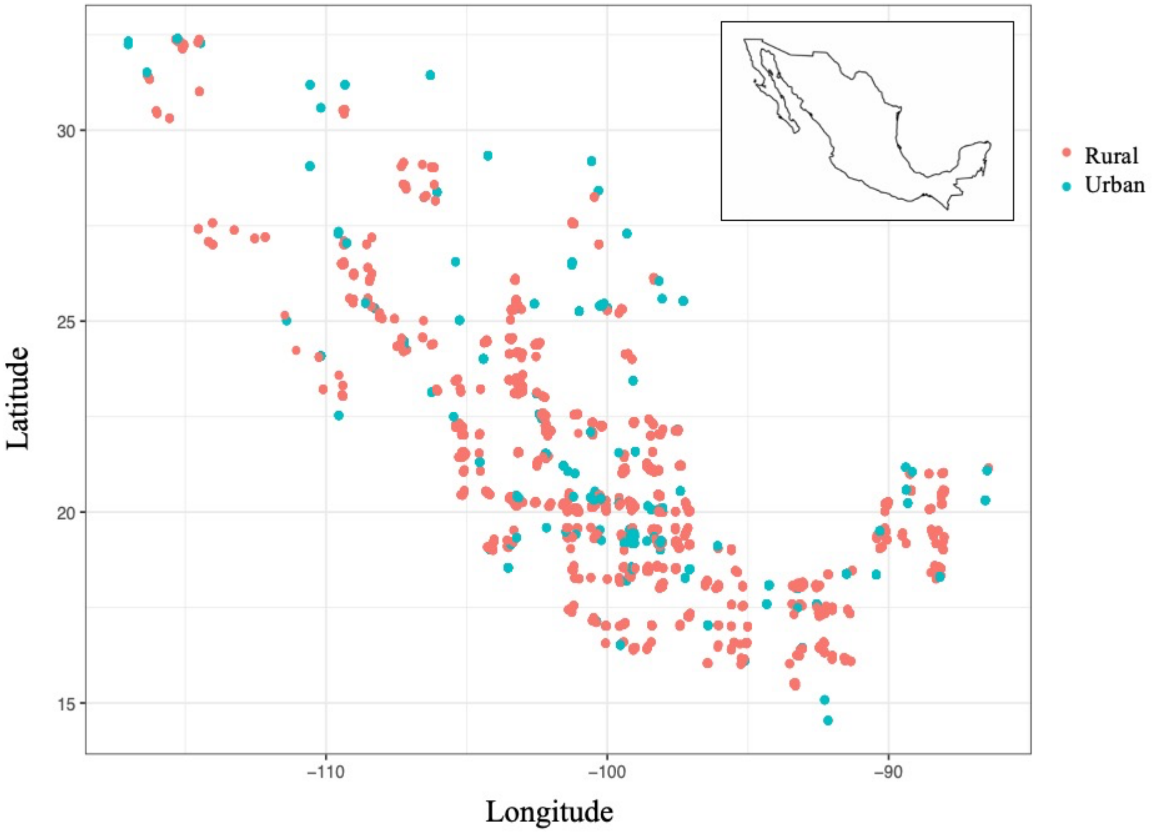


**Fig. S3. Distribution of rural and urban samples in the MXB.** Each dot represents the geographic location of a sampled locality with a minimum sample size of 5 individuals selected for genotyping. In total the MXB covers 898 localities, 321 municipalities, and 32 states across Mexico.

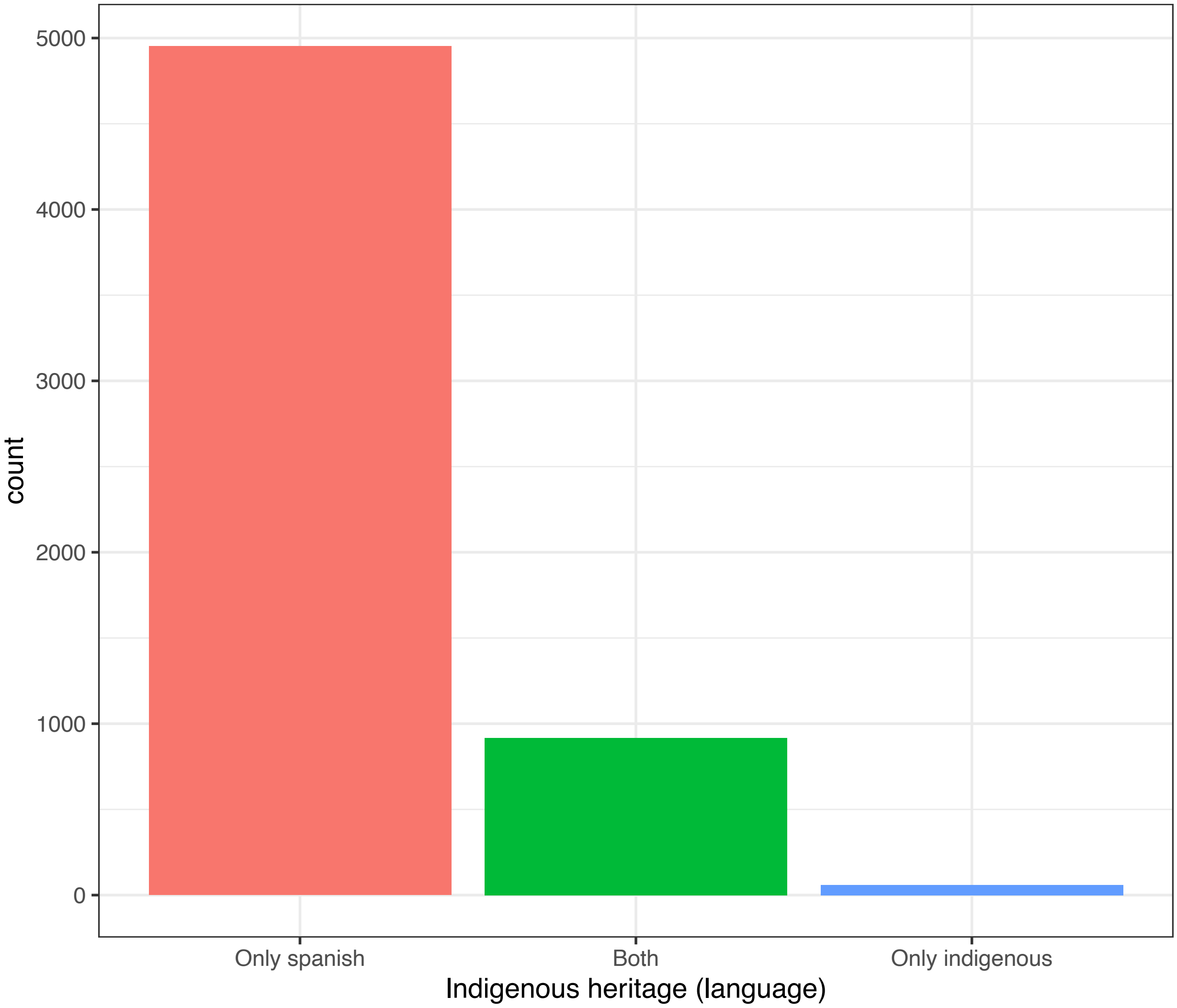


**Fig. S4. Number of MXB individuals that speak Spanish, an indigenous language or both.** Spanish speakers are shown in orange, those that speak both in green, and those that speak an only an indigenous language in blue.

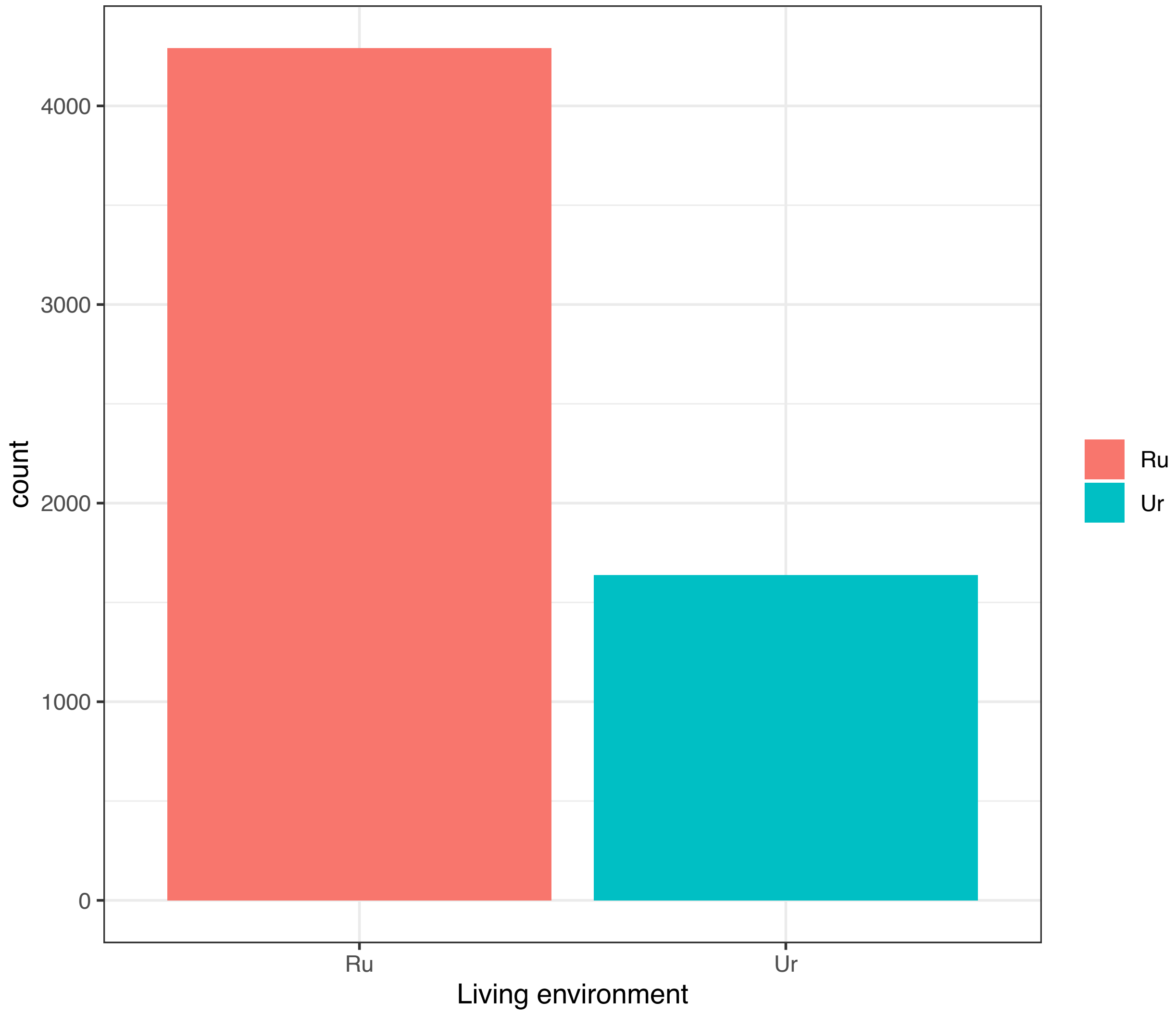


**Fig. S5. Number of MXB individuals from a rural or urban locality.** Rural localities are shown in orange, and urban localities in blue.

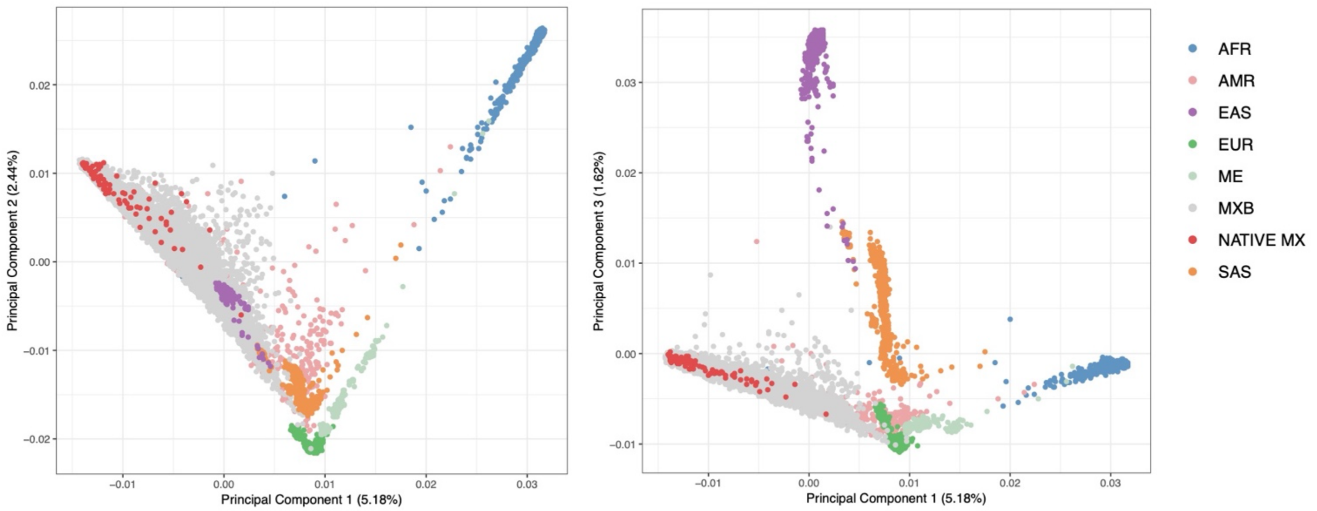


**Fig. S6. PCA analysis of MXB with global reference panels.** Predominant ancestry-proxy clusters inferred from Admixture are used to color present-day individuals used in PCA space solely to help with visualization (in groupings used by the reference datasets).

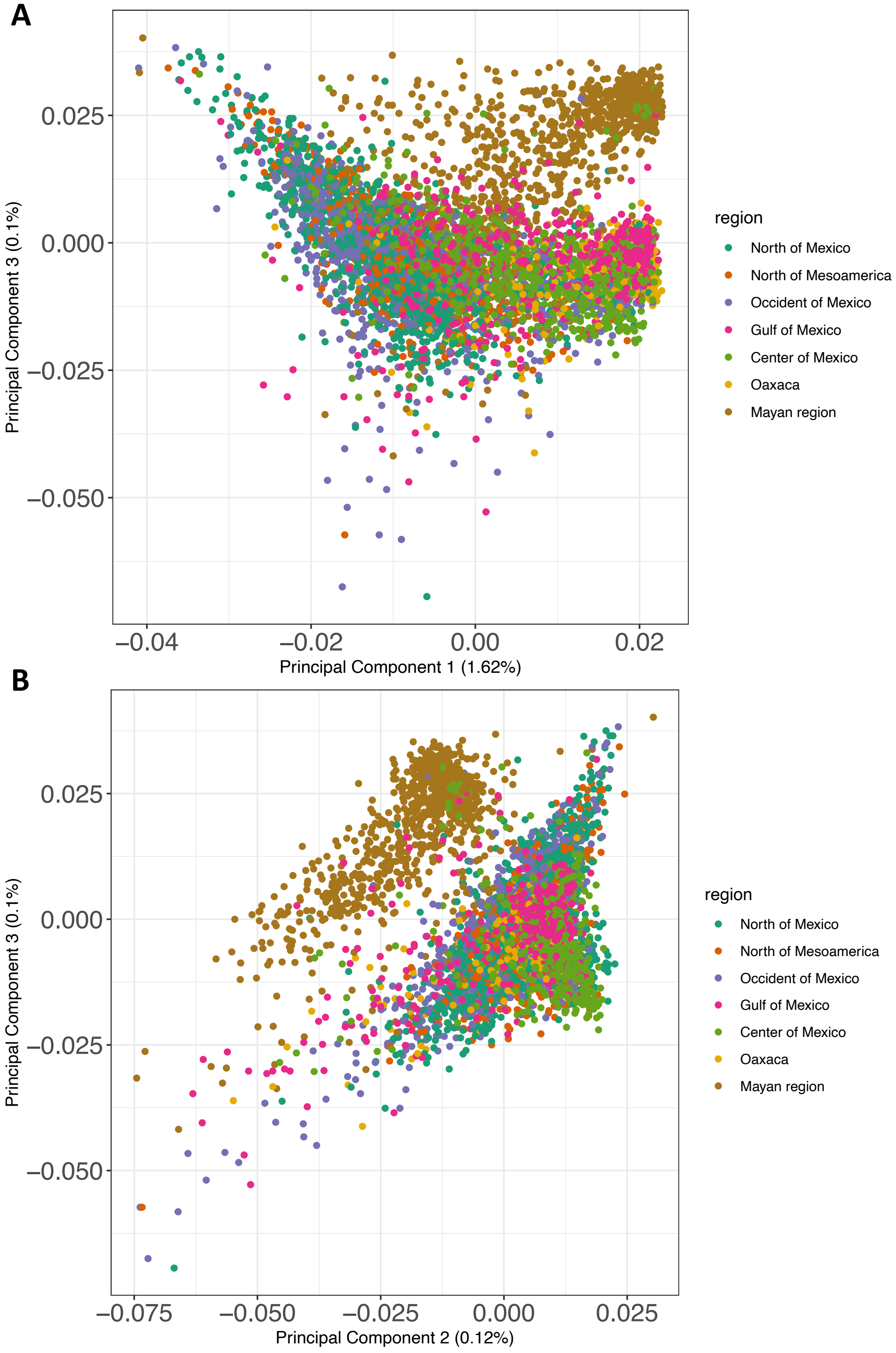


**Fig. S7. PCA analysis of MXB.**

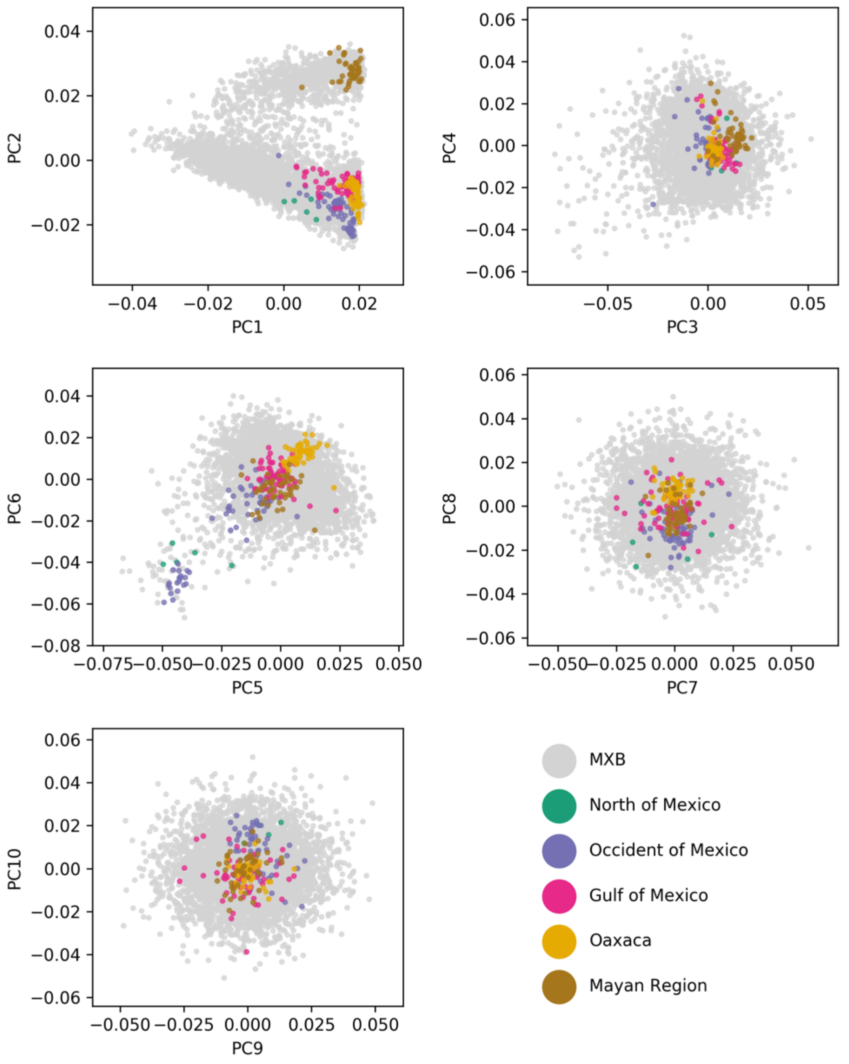


**Fig. S8. PCA analysis of MXB and NMDP grouped by Mesoamerican regions.**

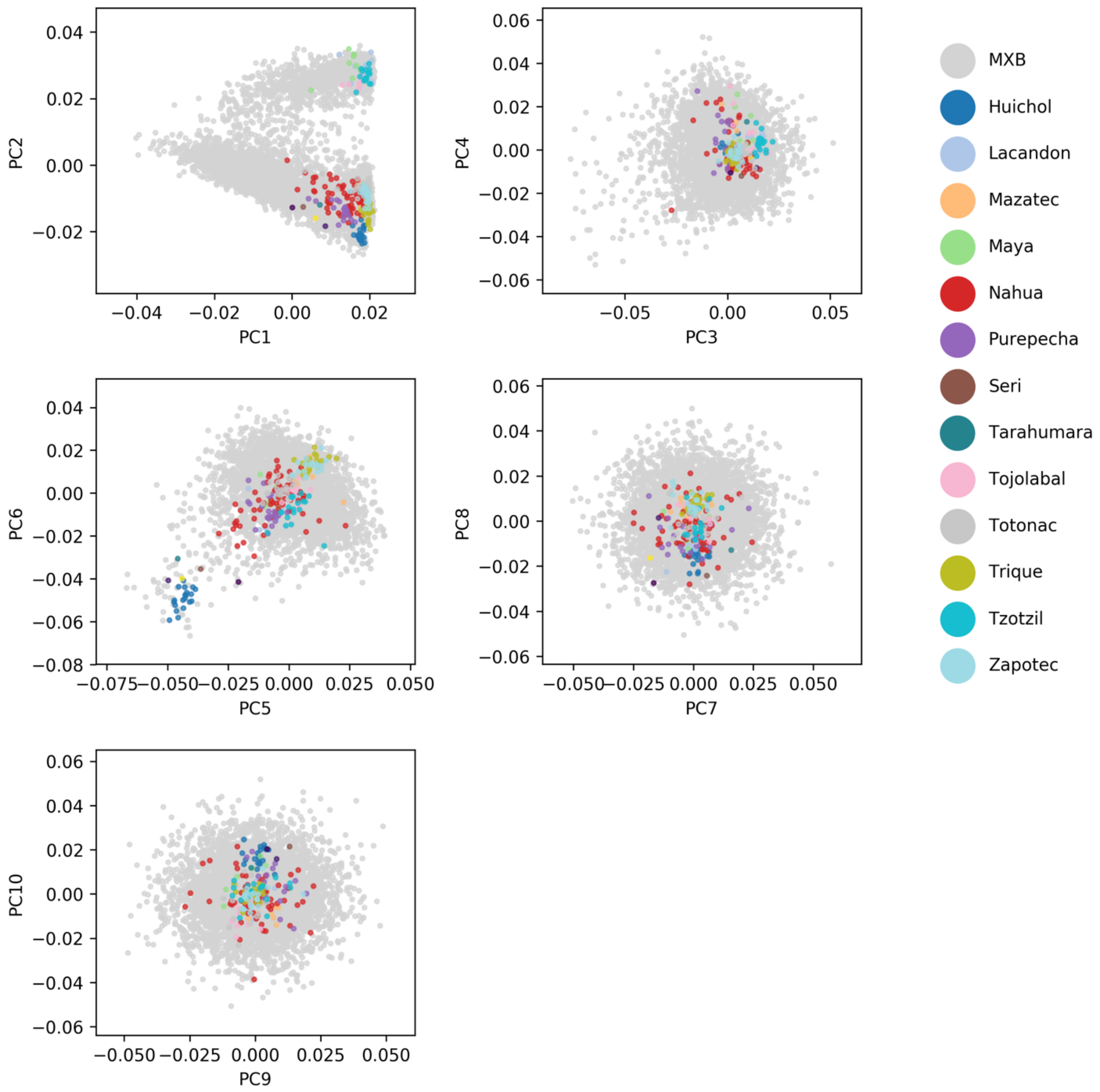


**Fig. S9. PCA analysis of MXB and NMDP grouped by indigenous culture.** NMDP individuals are classified according to the culture the individuals self-identify with[10].

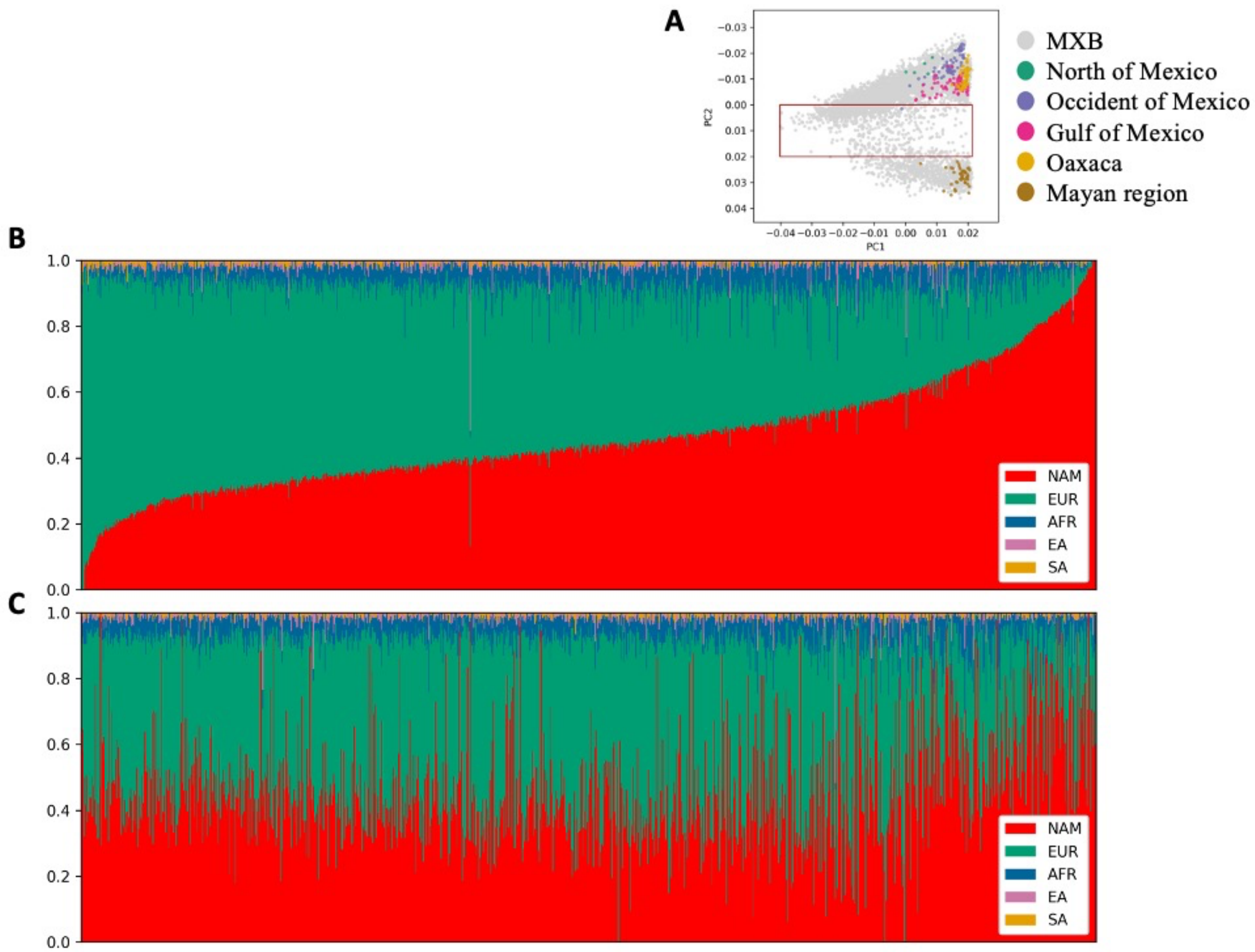


**Fig. S10. Admixture analysis of MXB in a principal component space defined by MXB and indigenous individuals from the NMDP.** (**A**) PCA analysis of MXB samples with indigenous groups from NMDP (2). Indigenous groups were divided into the Mesoamerican regions. A red box indicates the samples selected for parts (B) and (C). (**B**) Admixture analysis of MXB samples sorted by principal component 1. This axis captures the amount of ancestry from the Americas, with more native individuals found moving rightwards. (**C**) Admixture analysis of MXB samples sorted by principal component 2. This axis captures variation within ancestry from the America and does not result in any clear pattern at the scales of ancestries from different continents.

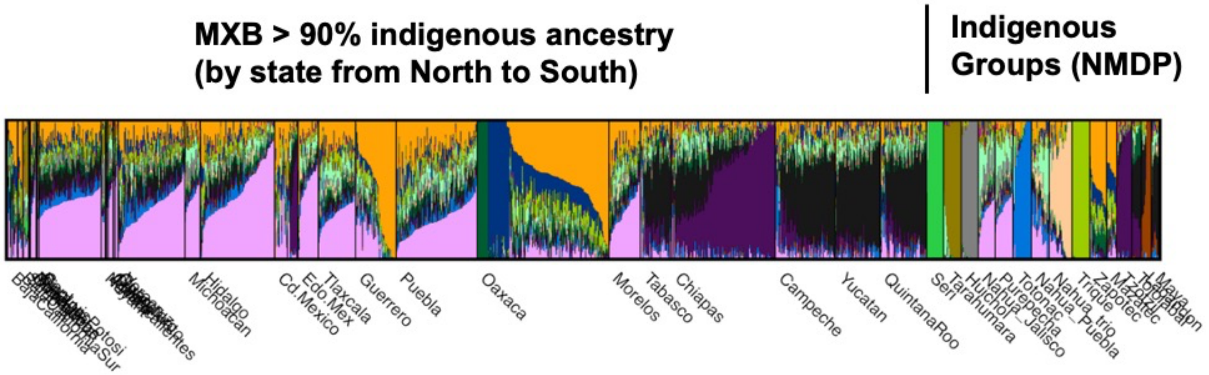


**Fig. S11. Admixture analysis of MXB and NMDP.** Only individuals that had significant ancestries from Central America were chosen for this analysis from the analysis in Figure 1. An admixture analysis was performed with indigenous groups from the NMDP (K=15).

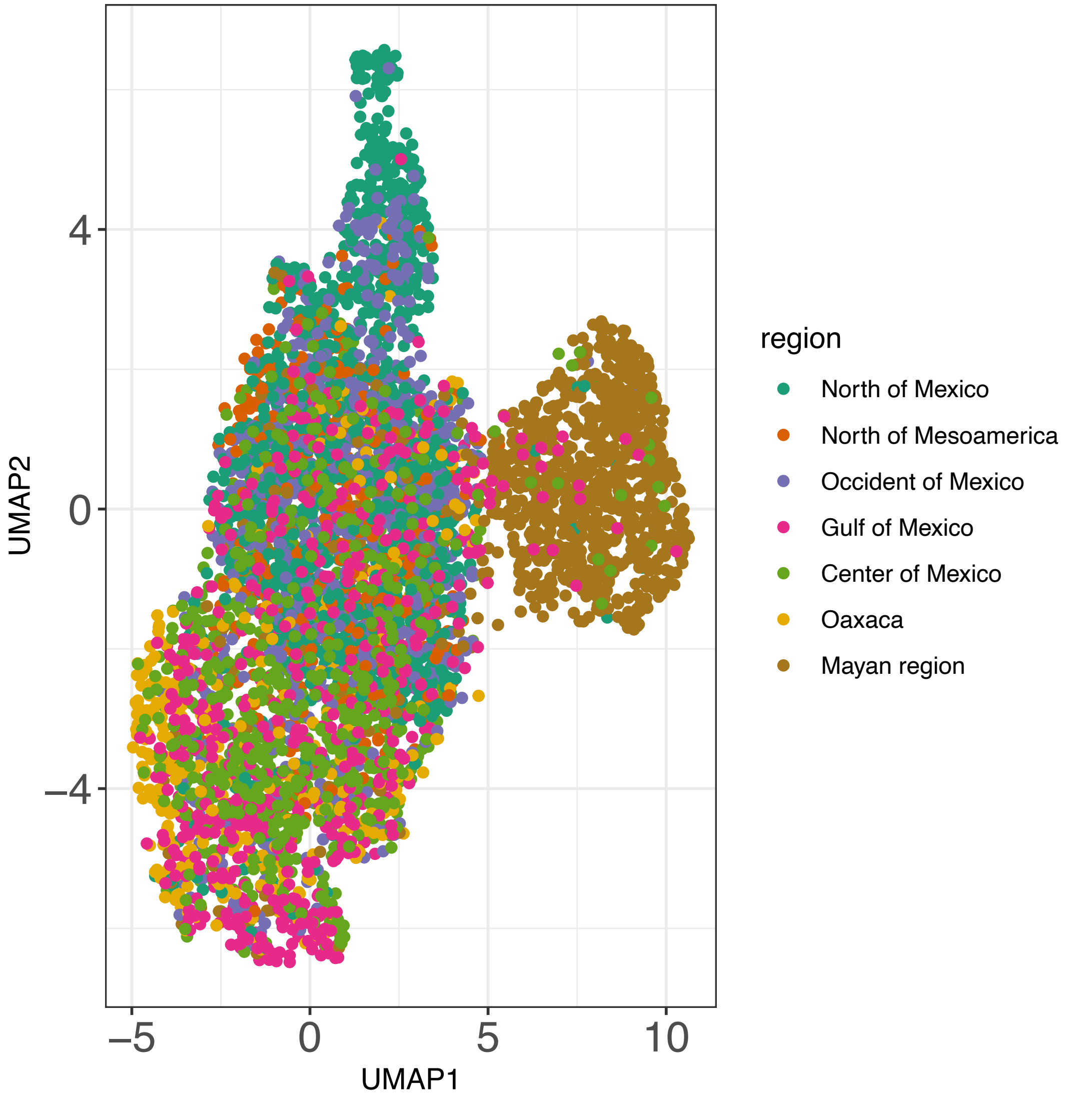


**Fig. S12. UMAP analysis of MXB samples with 10 principal components.**

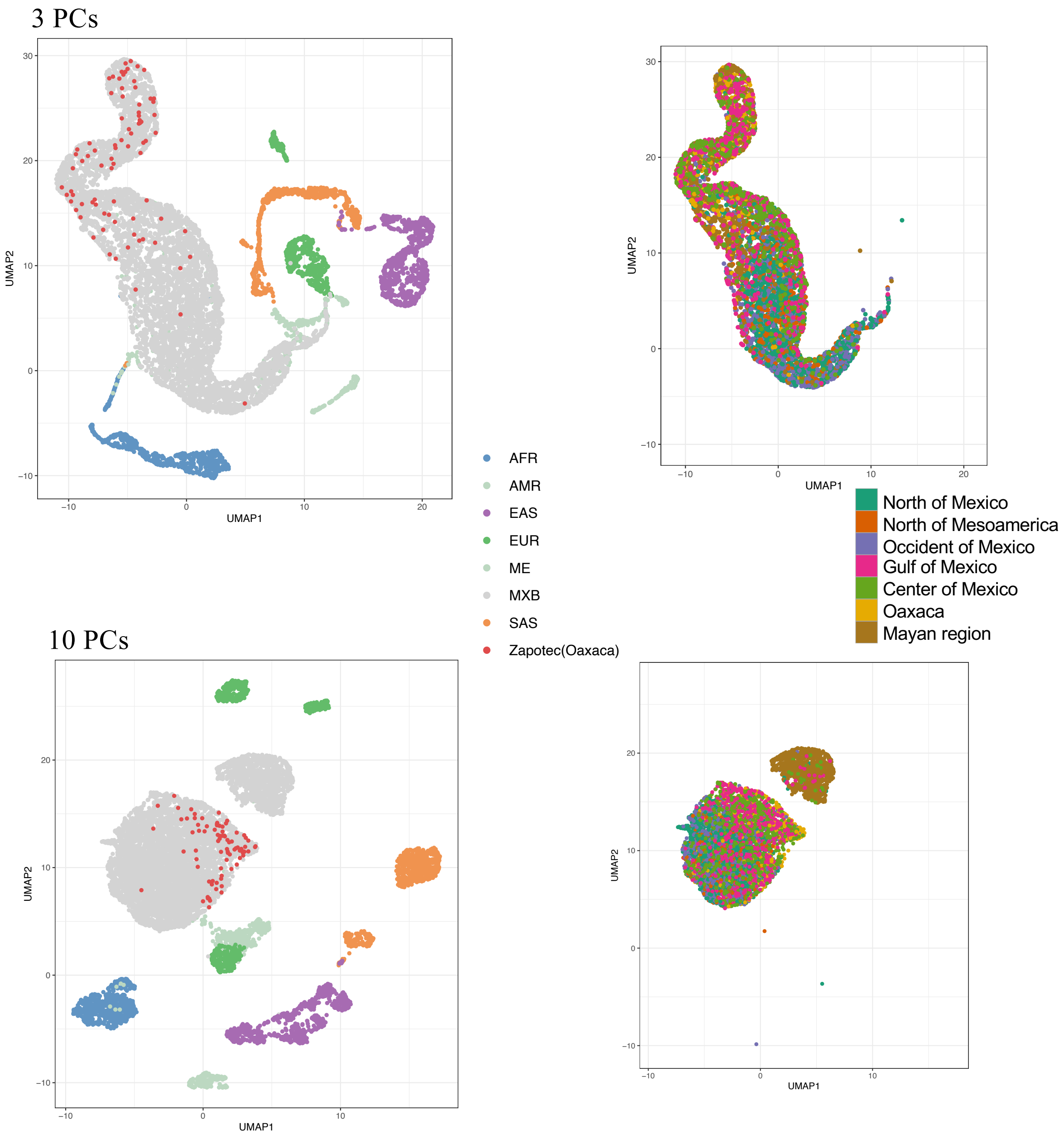


**Fig. S13. UMAP analysis of MXB samples with global reference panels.**

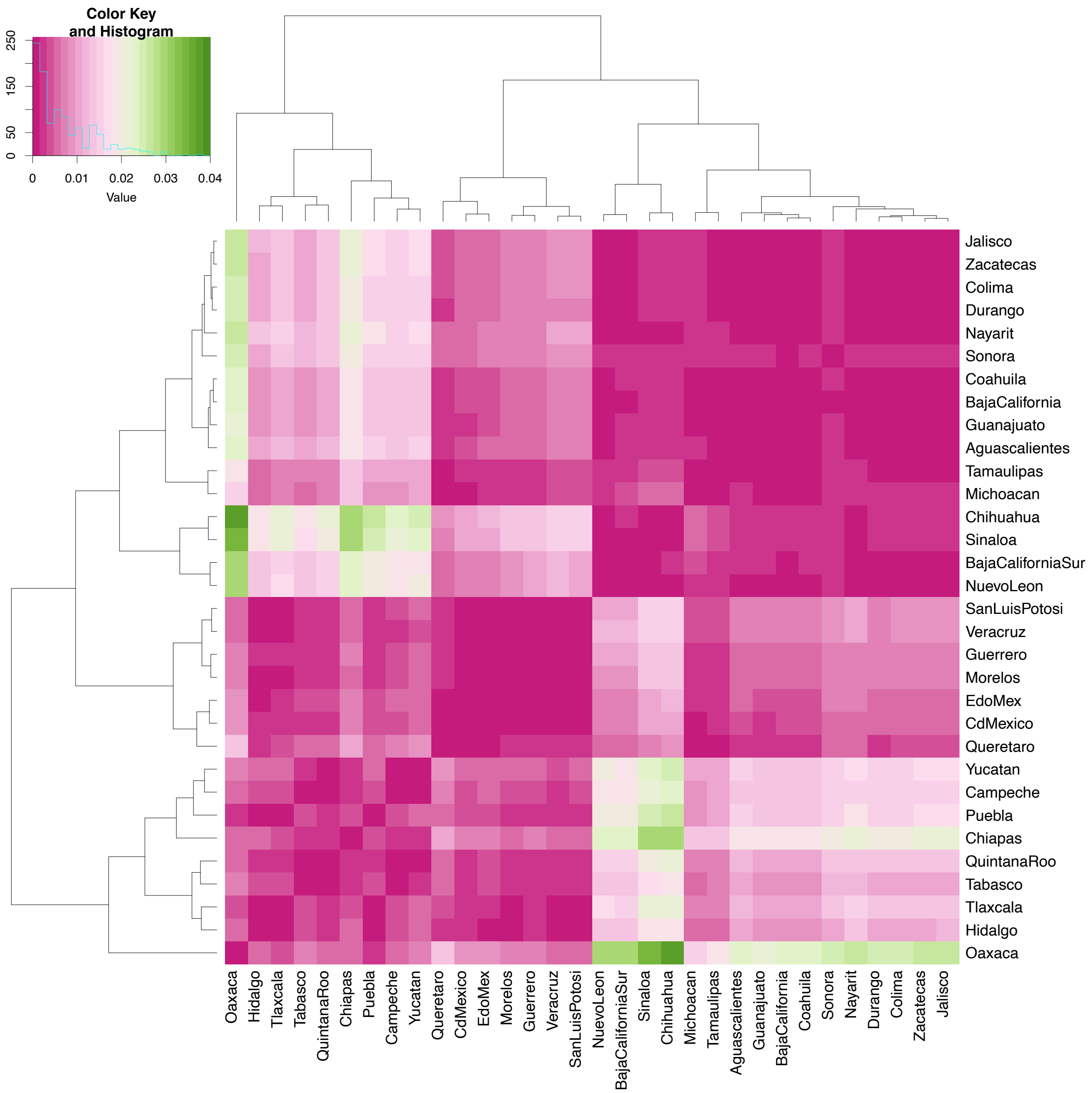


**Fig. S14. Fst analysis of autosomes of MXB individuals.**

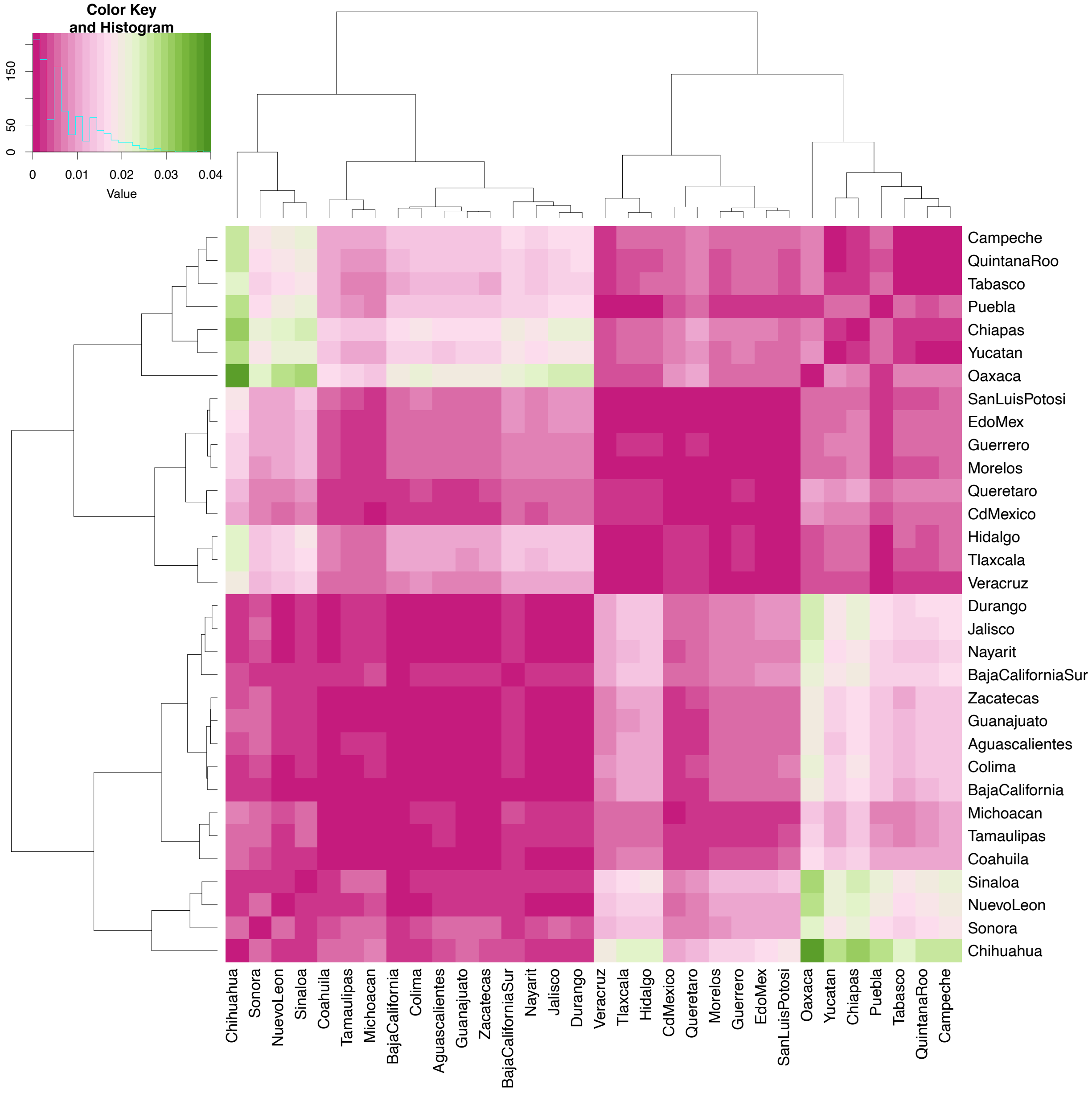


**Fig. S15. Fst analysis of X chromosomes of MXB individuals.**

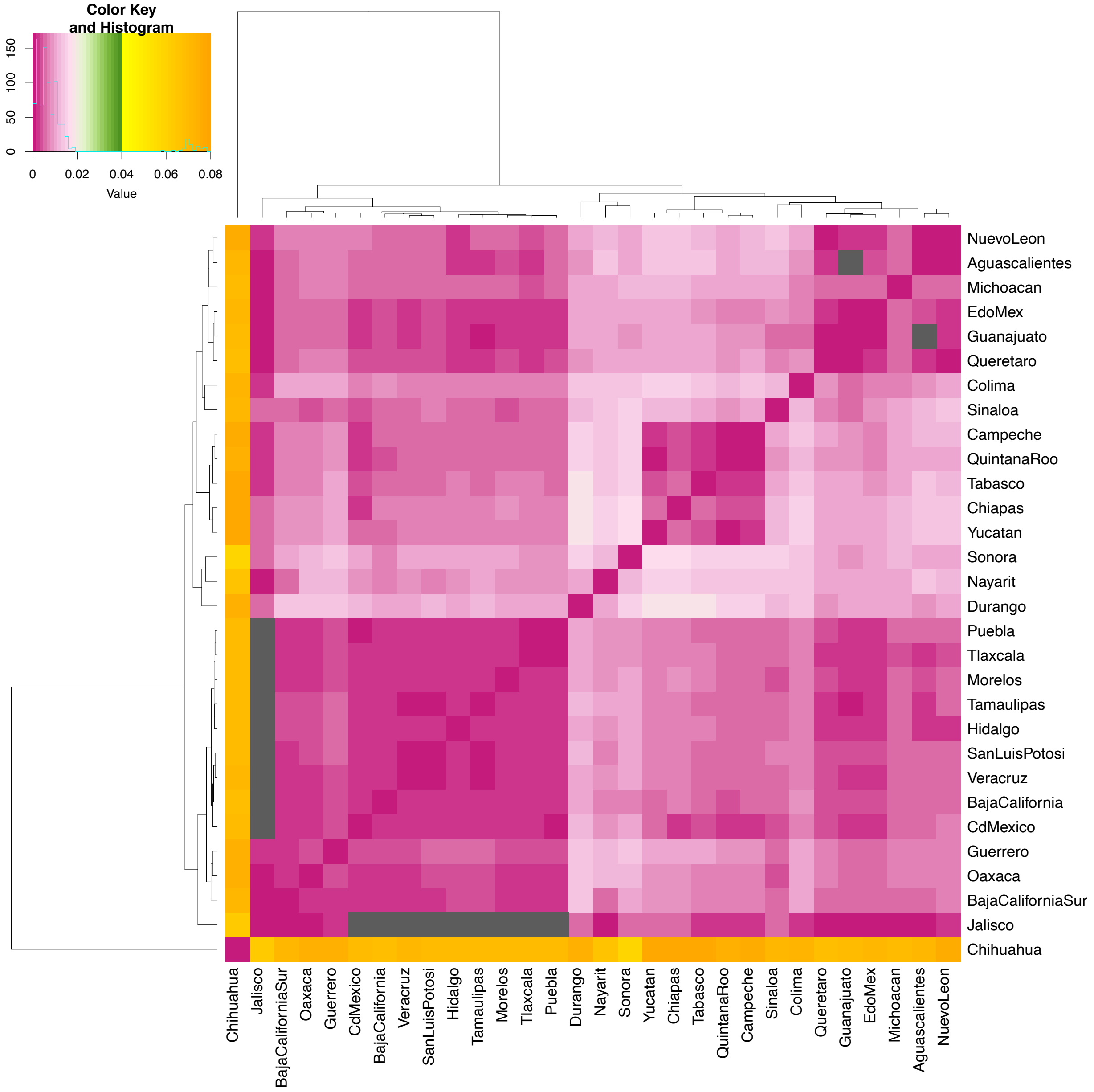


**Fig. S16. Fst analysis of autosomes of MXB individuals with high ancestries from Central America.** Only individuals with ≥ 90% ancestry from Central America (inferred using Admixture) were analyzed. Squares are colored black if a negative Fst value is estimated.

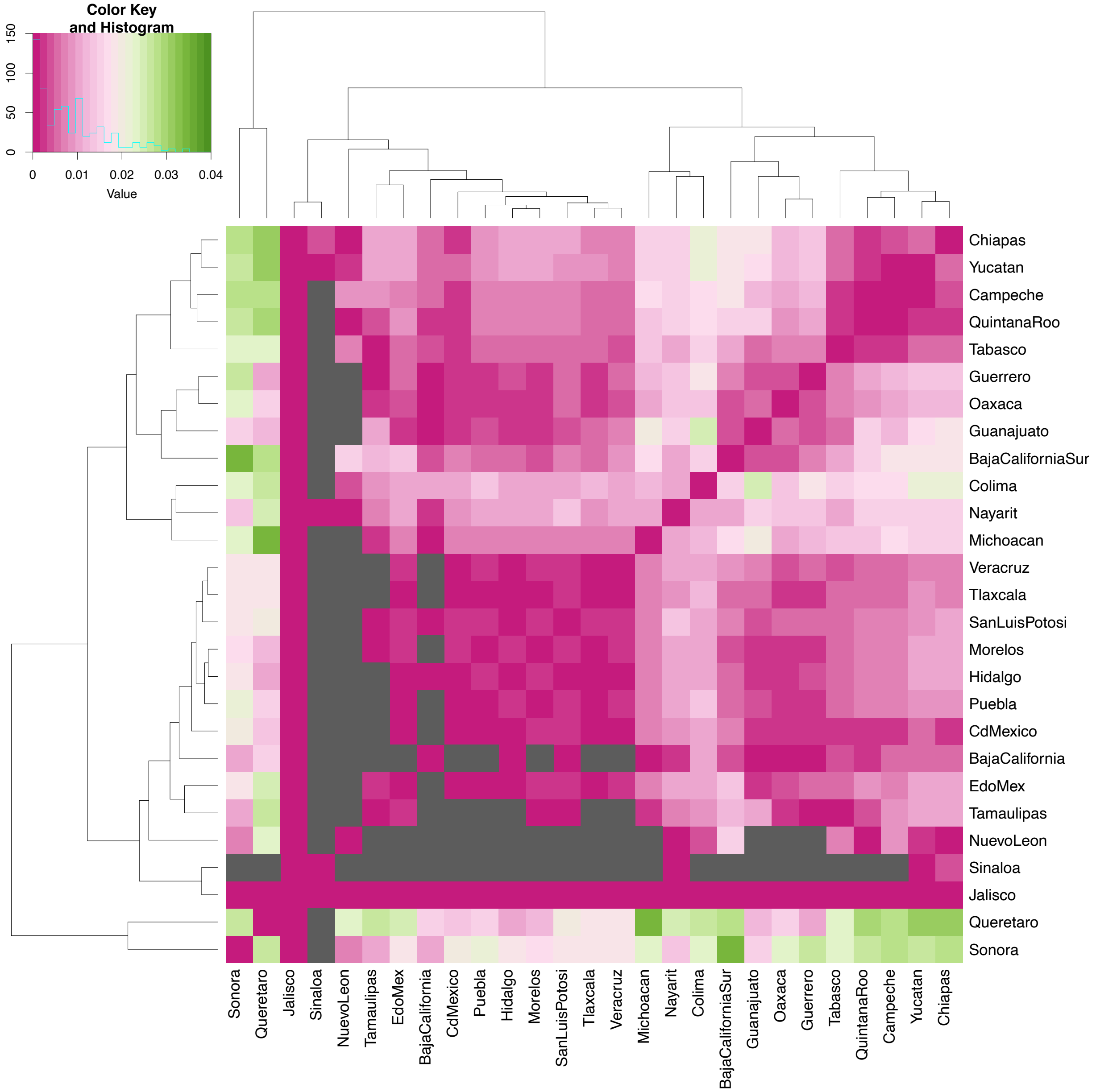


**Fig. S17. Fst analysis of X chromosomes of MXB individuals with high ancestries from Central America.** Only individuals with ≥ 90% ancestry from Central America (inferred using Admixture) were analyzed. Squares are colored black if a negative Fst value is estimated.

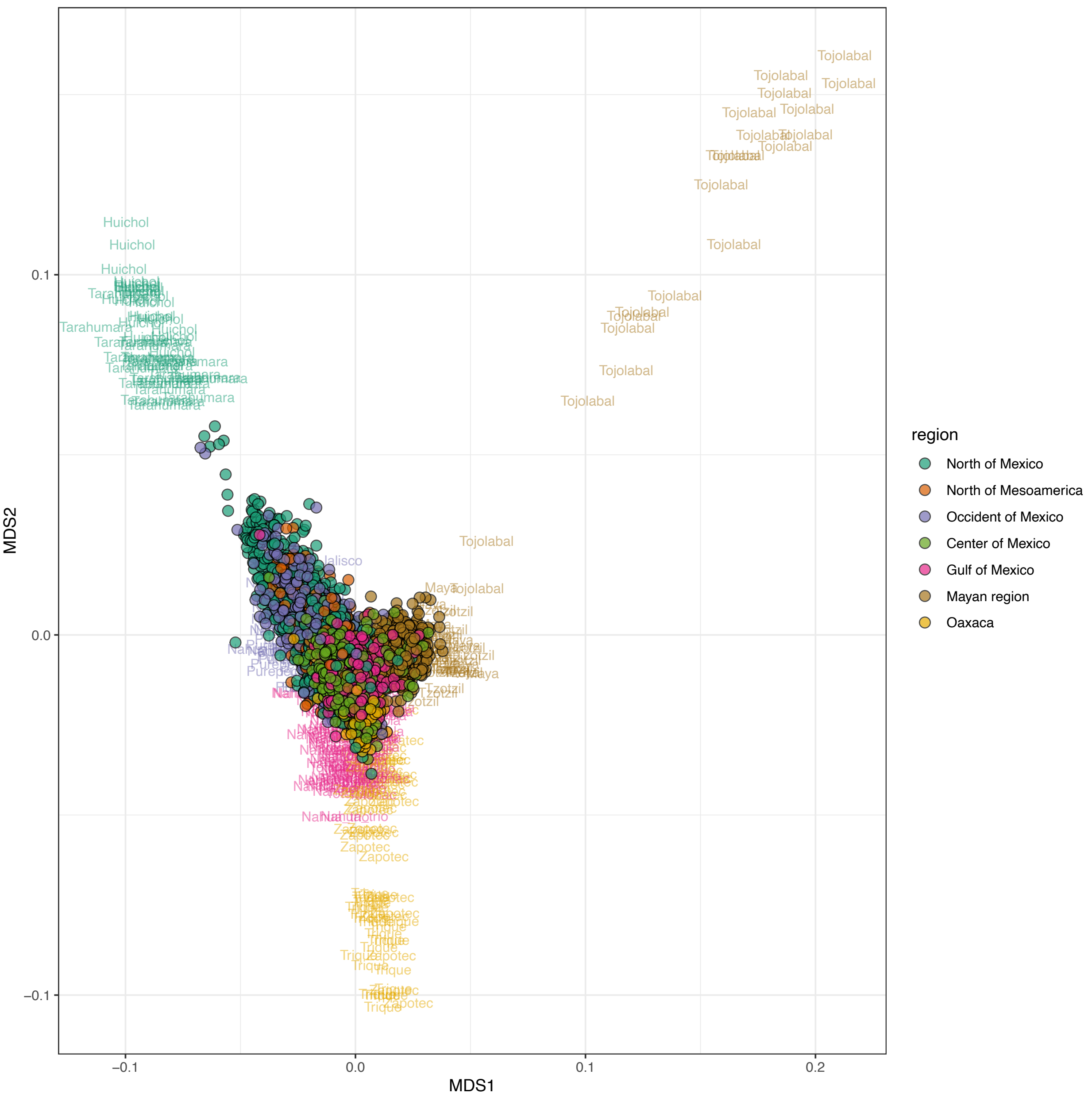


**Fig. S18. Origins of ancestries from Central America in MXB.** MAAS-MDS was used along with NDMP samples for this analysis.

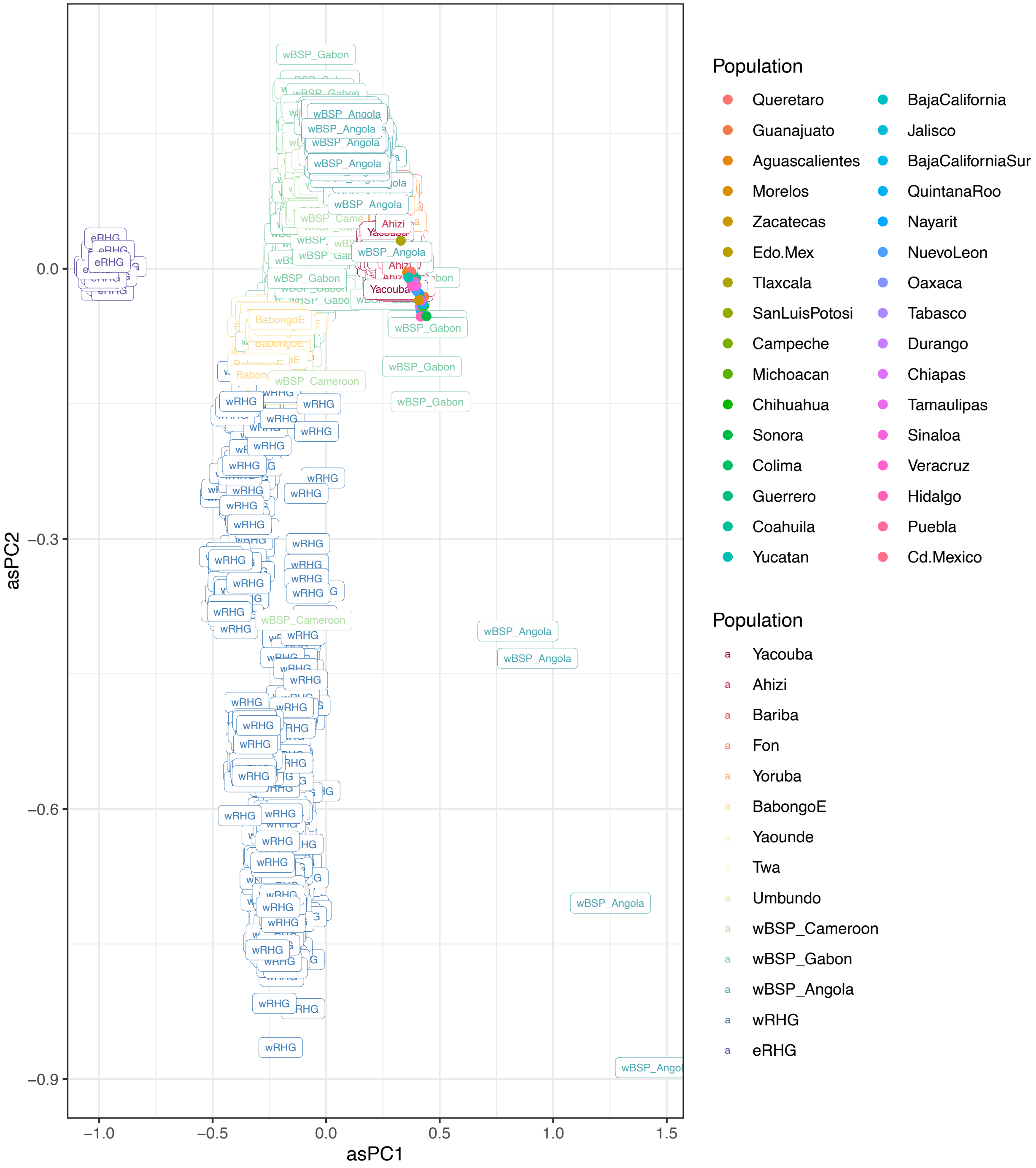


**Fig. S19. Origins of ancestries from West Africa in MXB.** Each state is plotted at the average values for asPC1 and asPC2 of each of its individuals.

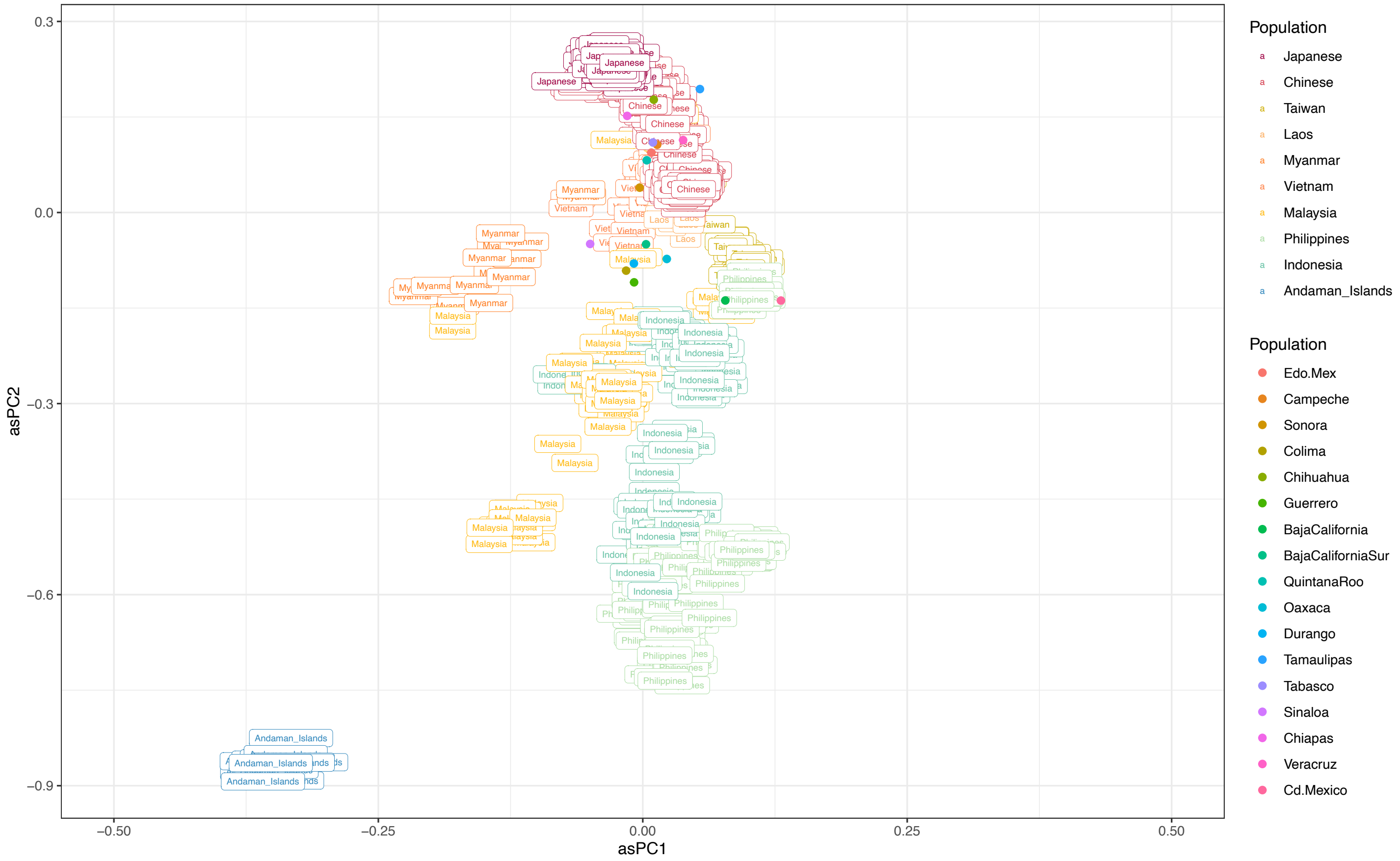


**Fig. S20. Origins of ancestries from East Asia in MXB.** Each state is plotted at the average values for asPC1 and asPC2 of each of its individuals.

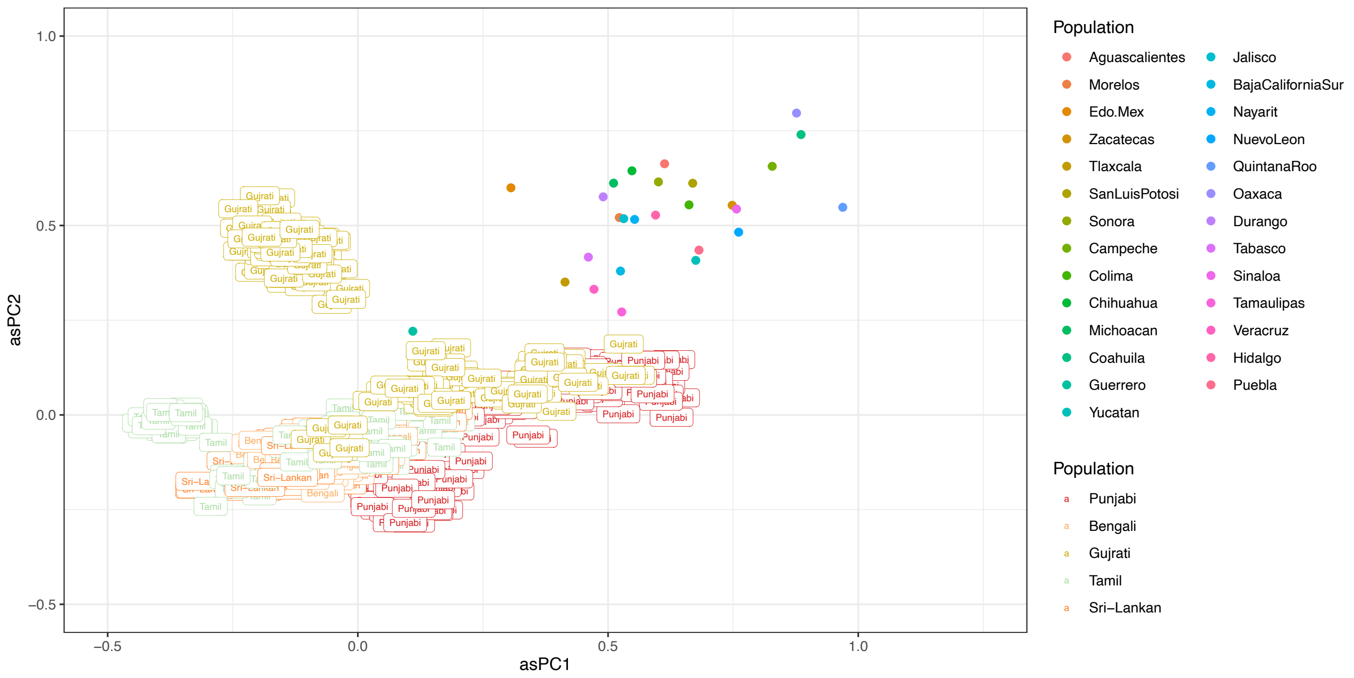


**Fig. S21. Origins of ancestries from South Asia in MXB.** Each state is plotted at the average values for asPC1 and asPC2 of each of its individuals.

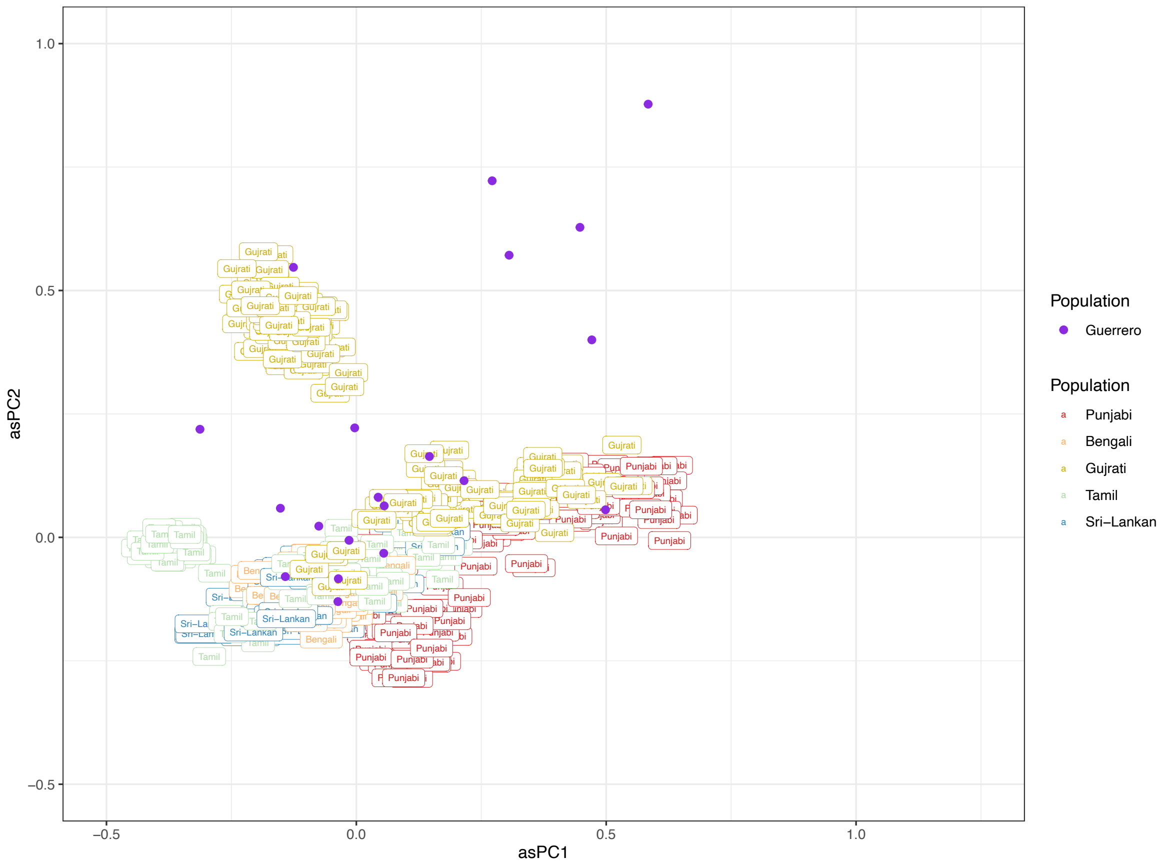


**Fig. S22. Origins of ancestries from South Asia in MXB for Guerrero.**

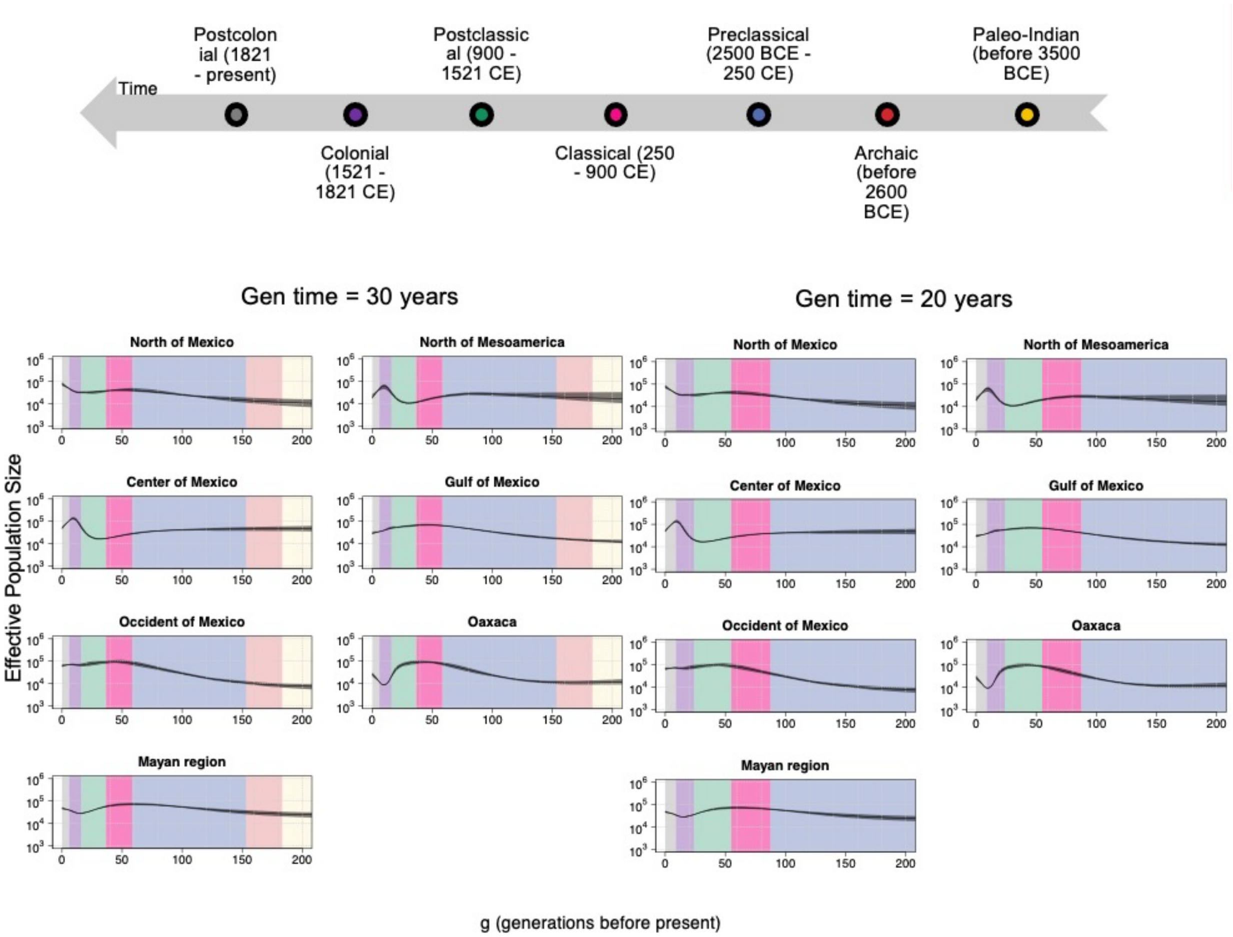


**Fig. S23. asIBDNe analysis results for ancestries from Central America are visualized for 30 or 20 years per generation.**

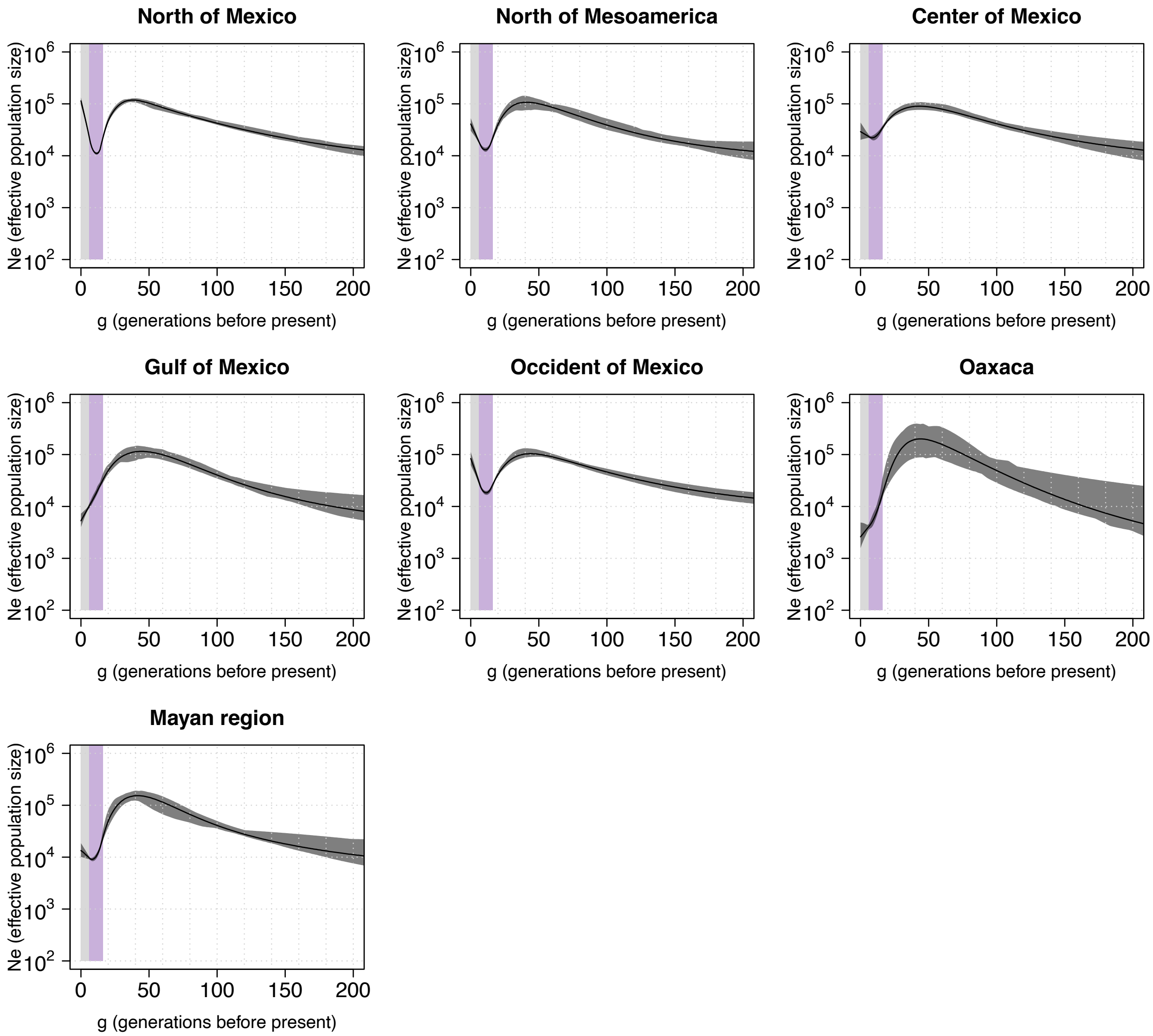


**Fig. S24. Estimated effective population size for ancestries from Western Europe by Mesoamerican region in Mexico (30 years per generation).**

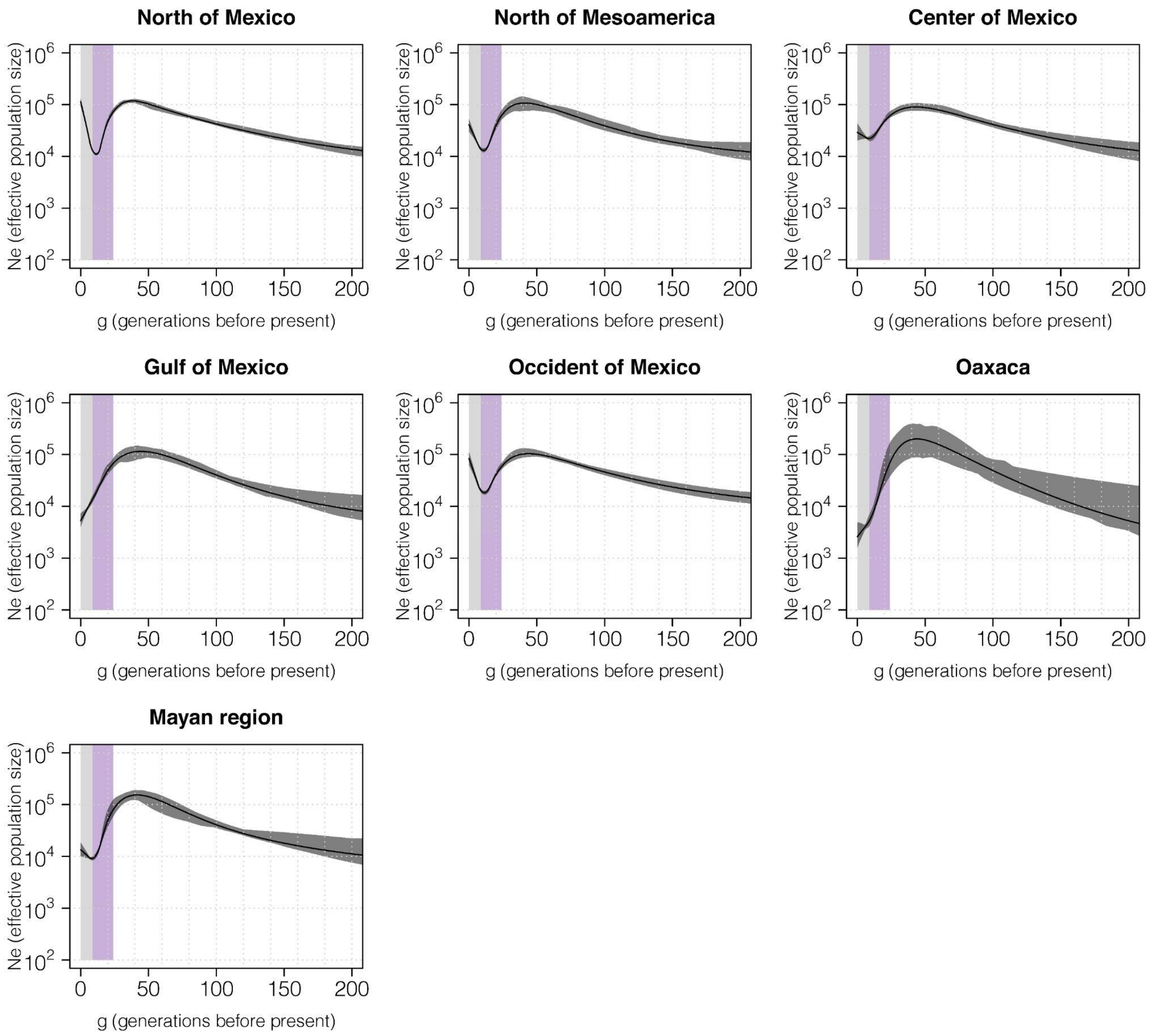


**Fig. S25. Estimated effective population size for ancestries from Western Europe by Mesoamerican region in Mexico (20 years per generation).**

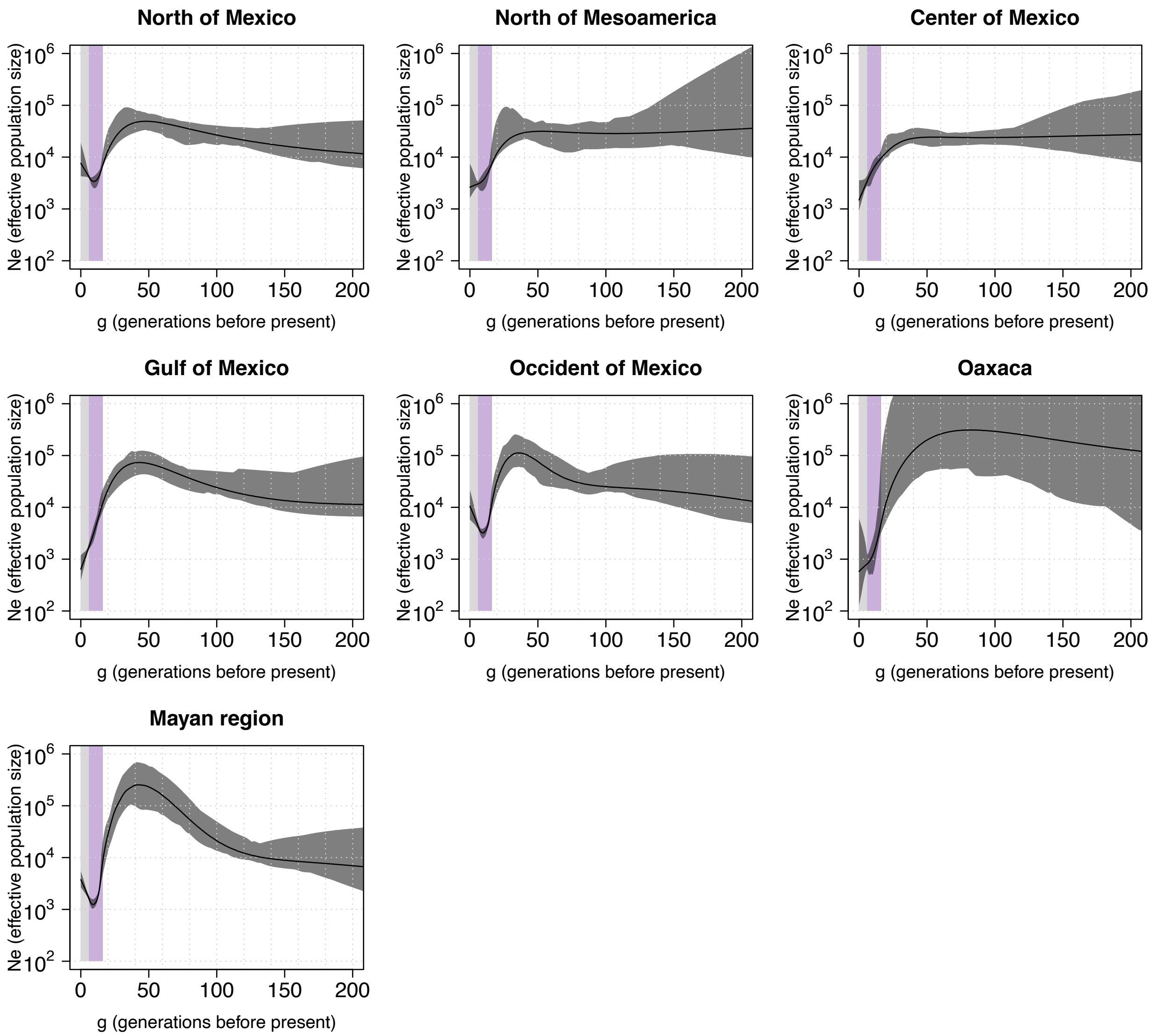


**Fig. S26. Estimated effective population size for ancestries from West Africa by Mesoamerican region in Mexico (30 years per generation).**

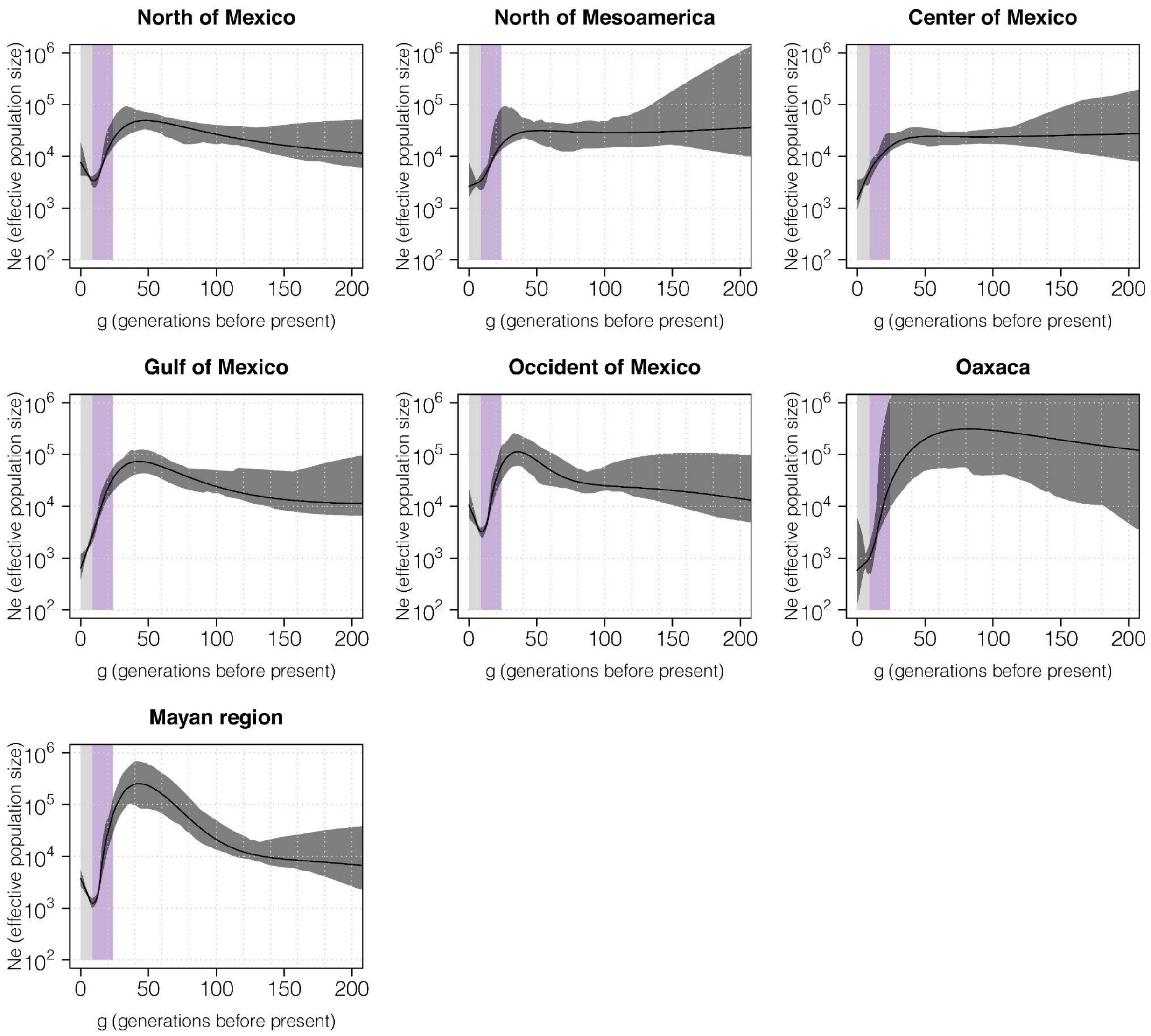


**Fig. S27. Estimated effective population size for ancestries from West Africa by Mesoamerican region in Mexico (20 years per generation).**

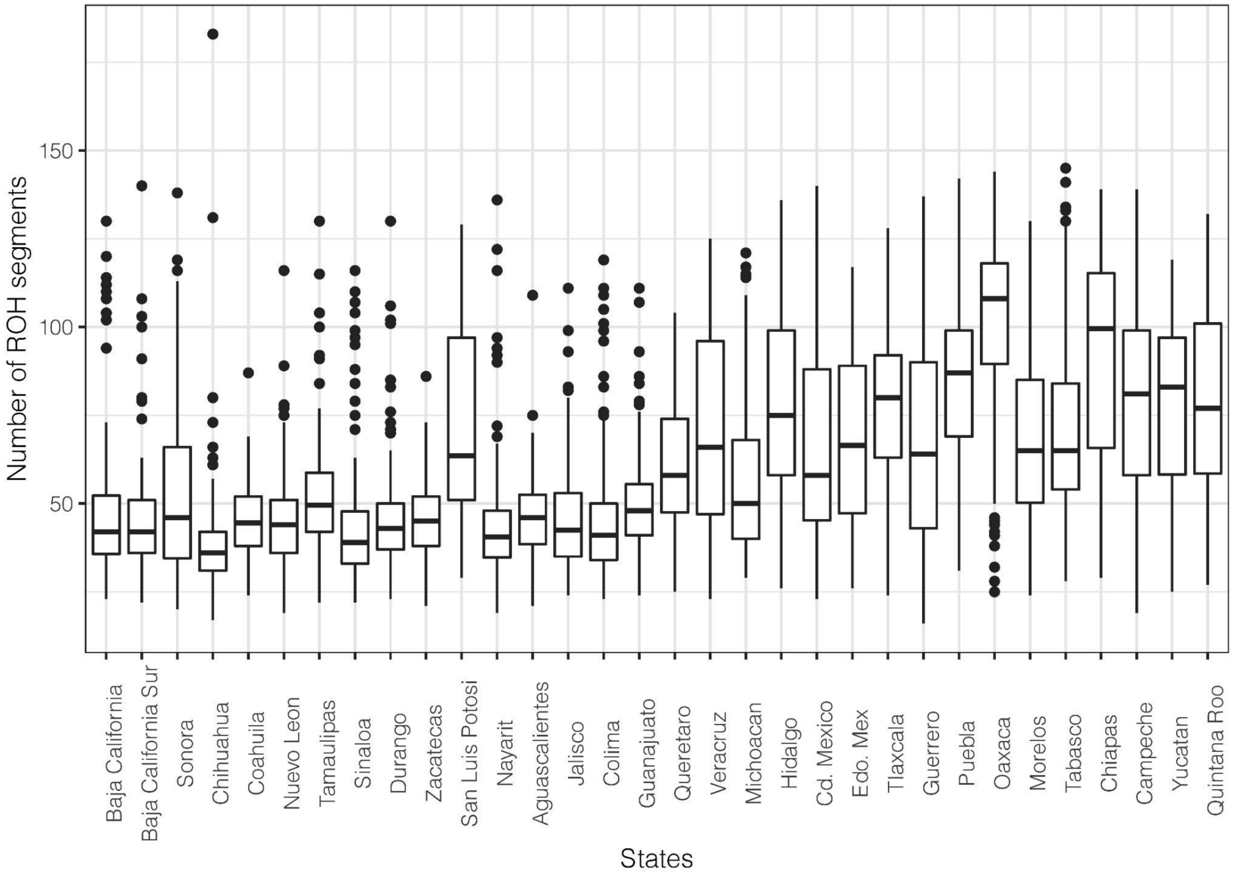


**Fig. S28. NROH distribution by state.**

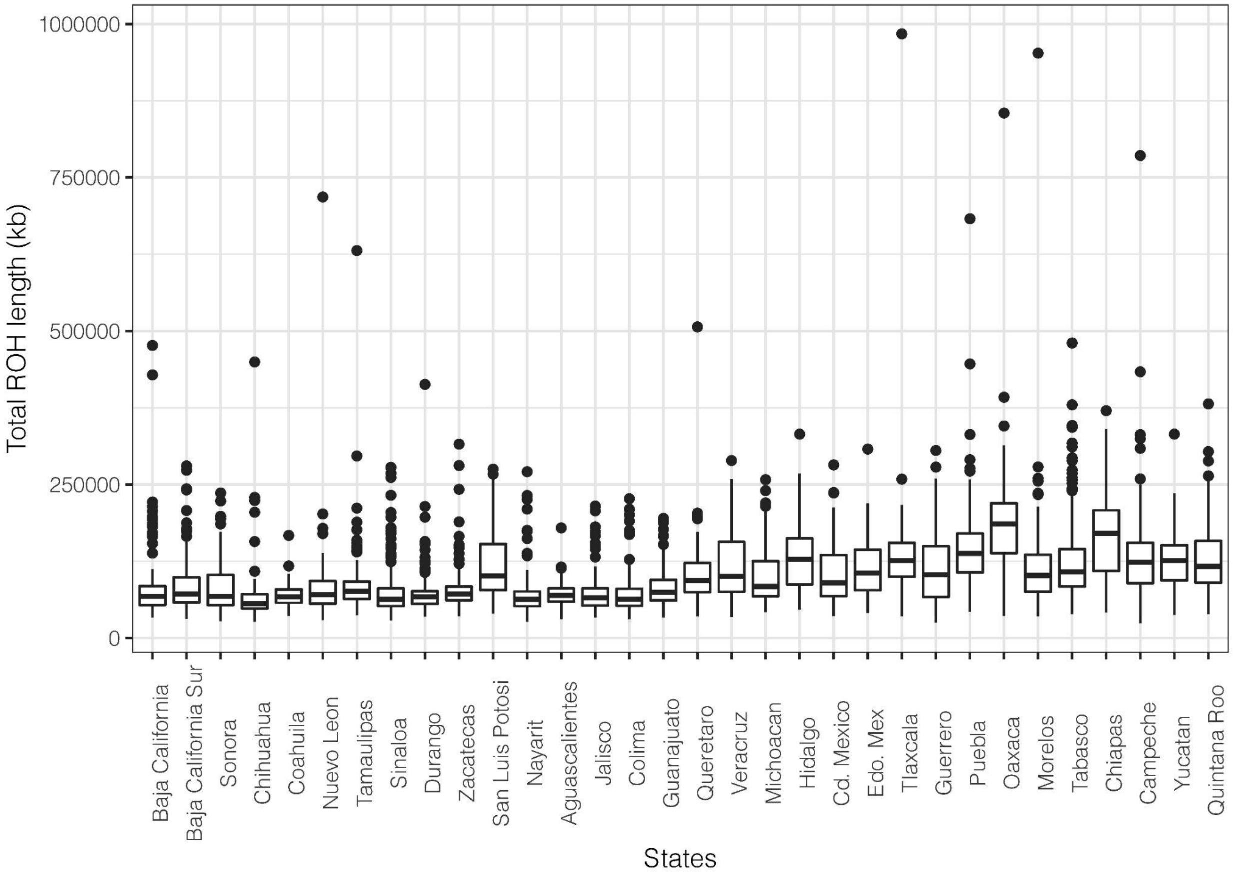


**Fig. S29. SROH distribution by state.**

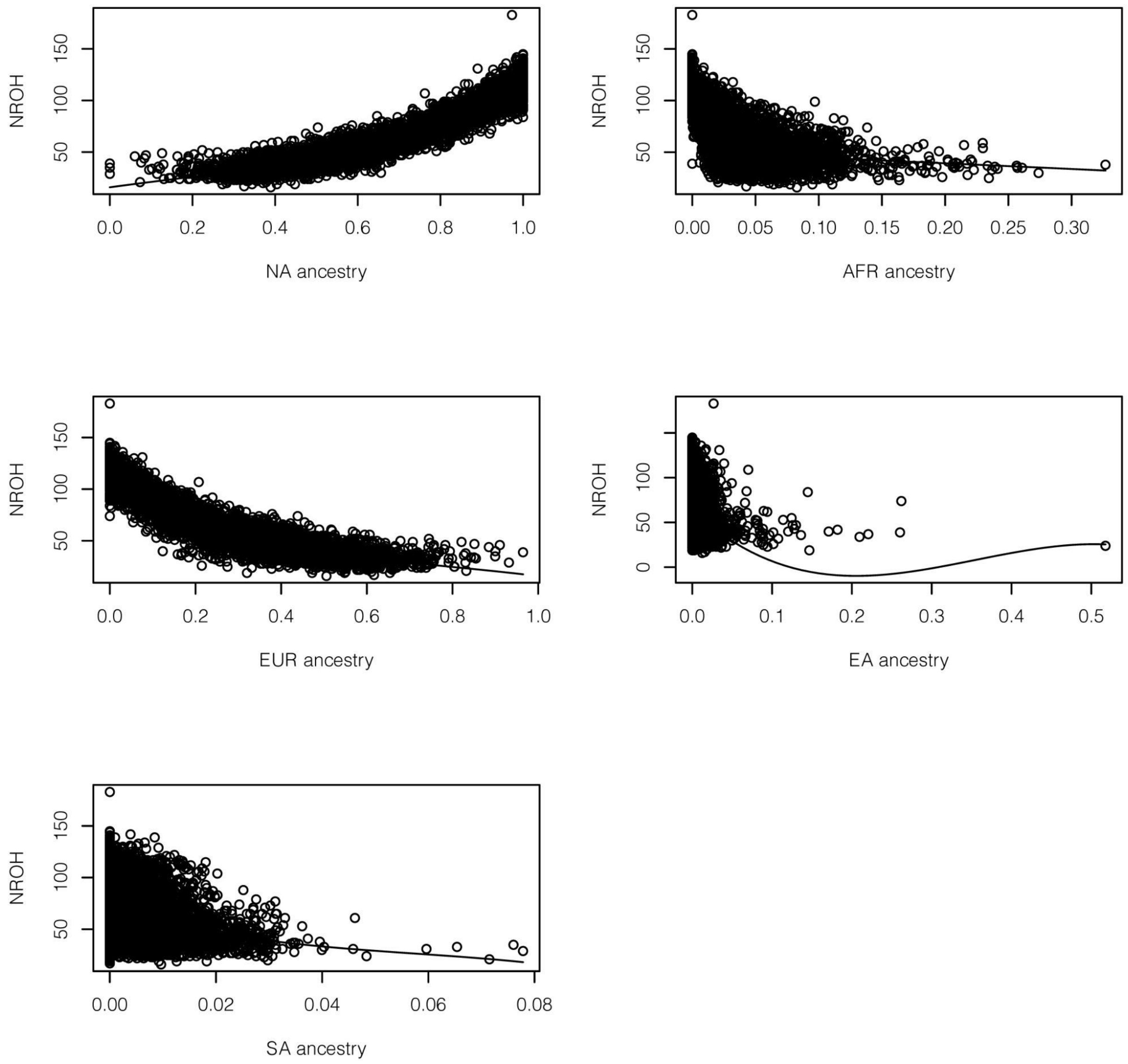


**Fig. S30. NROH are correlated with ancestries in MXB.** Ancestry proxies are inferred from ADMIXTURE.

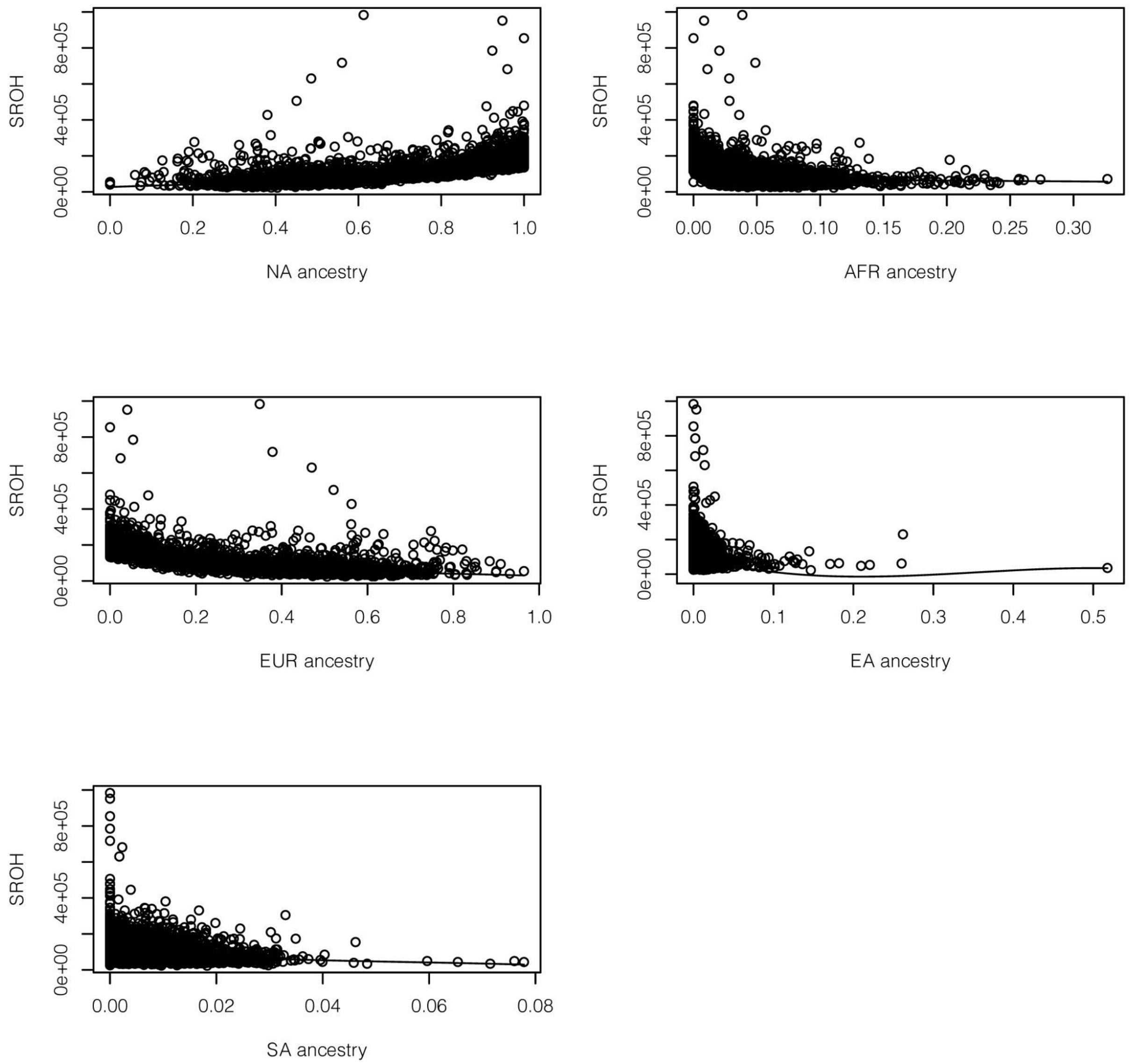


**Fig. S31. SROH are correlated with ancestries in MXB.** Ancestry proxies are inferred from ADMIXTURE.

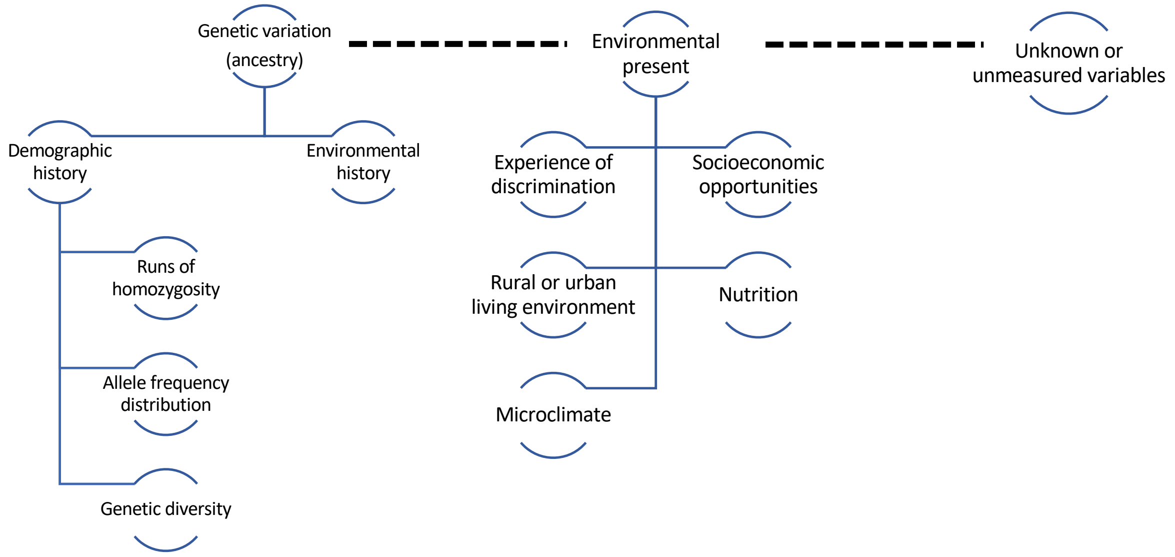


**Fig. S32. A framework for the role of genetics and environment in creating genetic variation and variation in complex traits and disease risk used in this study.** Breakdown of genetic variation (ancestries) as demographic (reflected in patterns of ROH, allele frequencies and genetic diversity) and environmental history, and its cross-talk with environmental present and unknown or unmeasured variables. The interrelation between genetics and environment is very complex and there are likely variables unknown or unmeasured that undoubtedly participate in the framework.

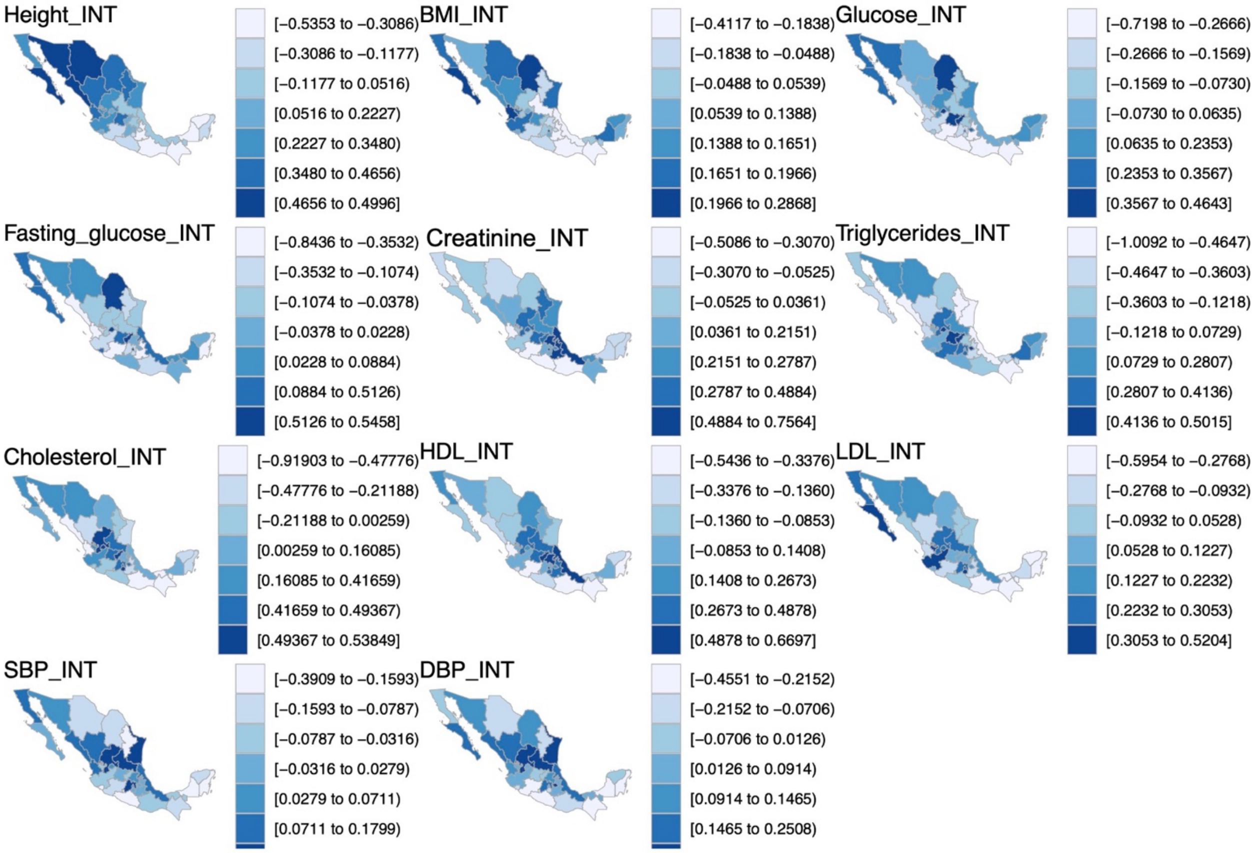


**Fig. S33. Average trait values per state visualized on the map of Mexico.** Quantitative traits were inverse normalized and average values for transformed traits per state are shown here. See methods (The Mexican Biobank Project: phenotype, lifestyle and environmental data) for further details on quantitative traits analyzed.

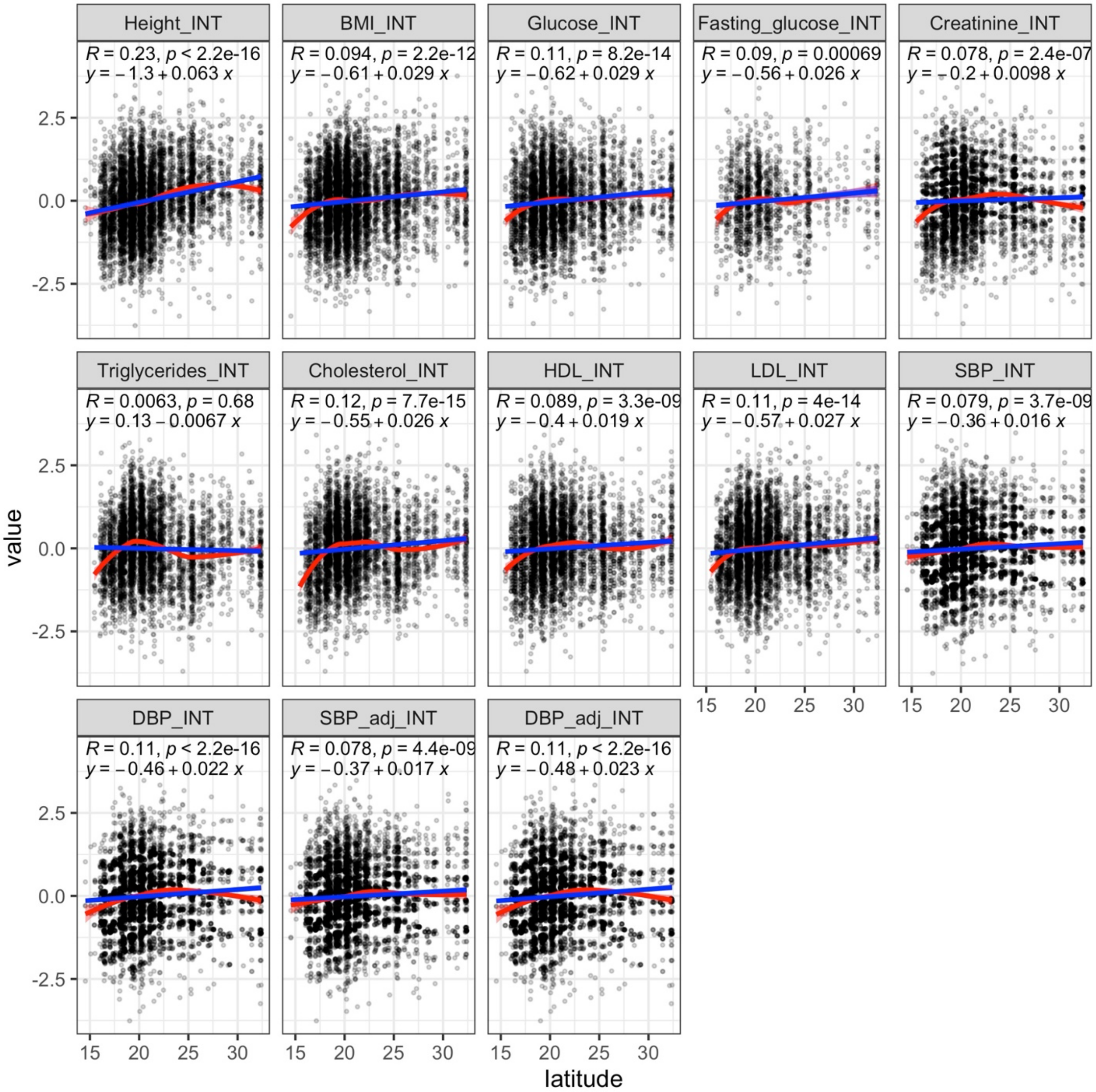


**Fig. S34. Complex trait variation by latitude.** To aid with visualization, a smoothed conditional mean line is shown using smoothing method loess (red) or lm/linear model (blue).

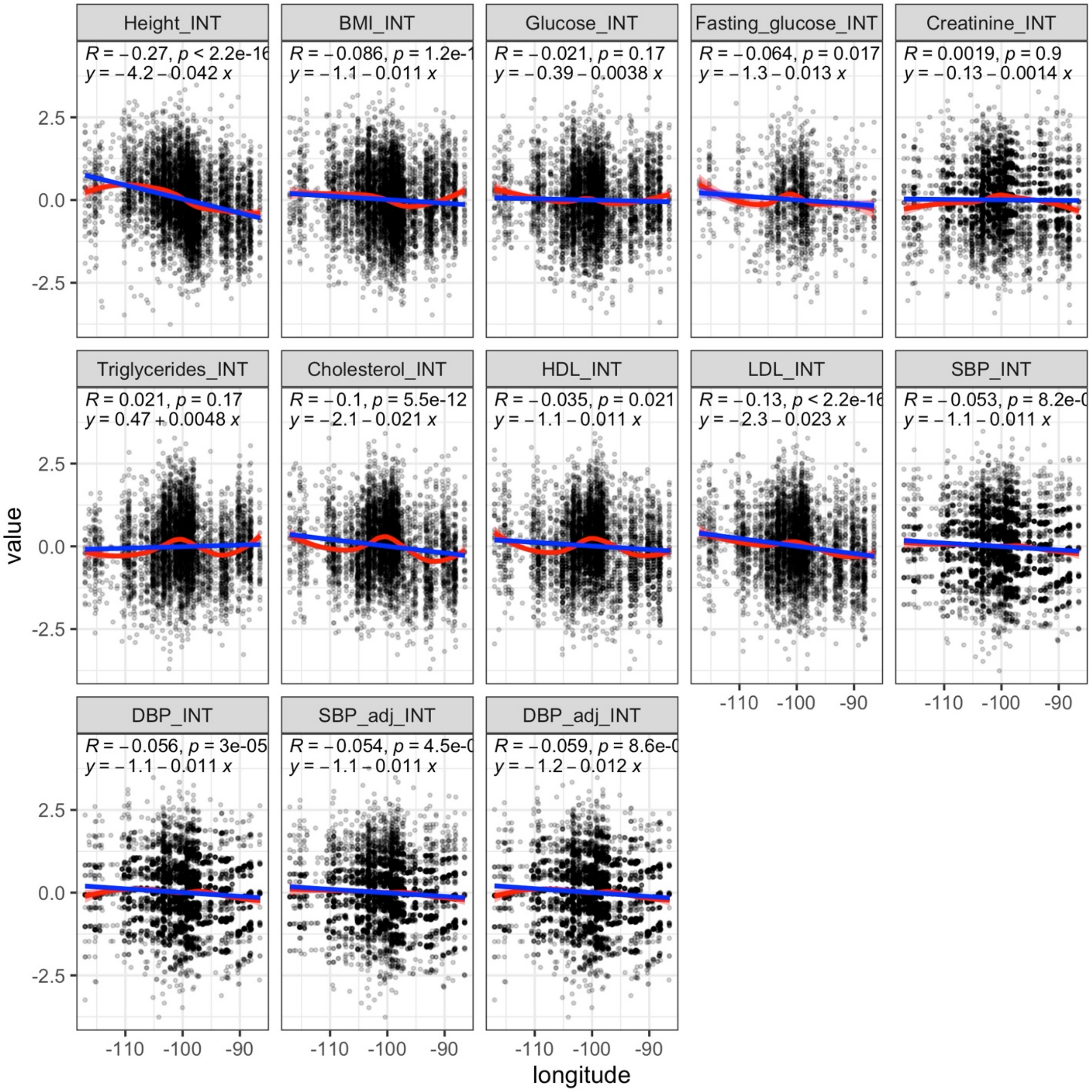


**Fig. S35. Complex trait variation by longitude.** To aid with visualization, a smoothed conditional mean line is shown using smoothing method loess (red) or lm/linear model (blue).

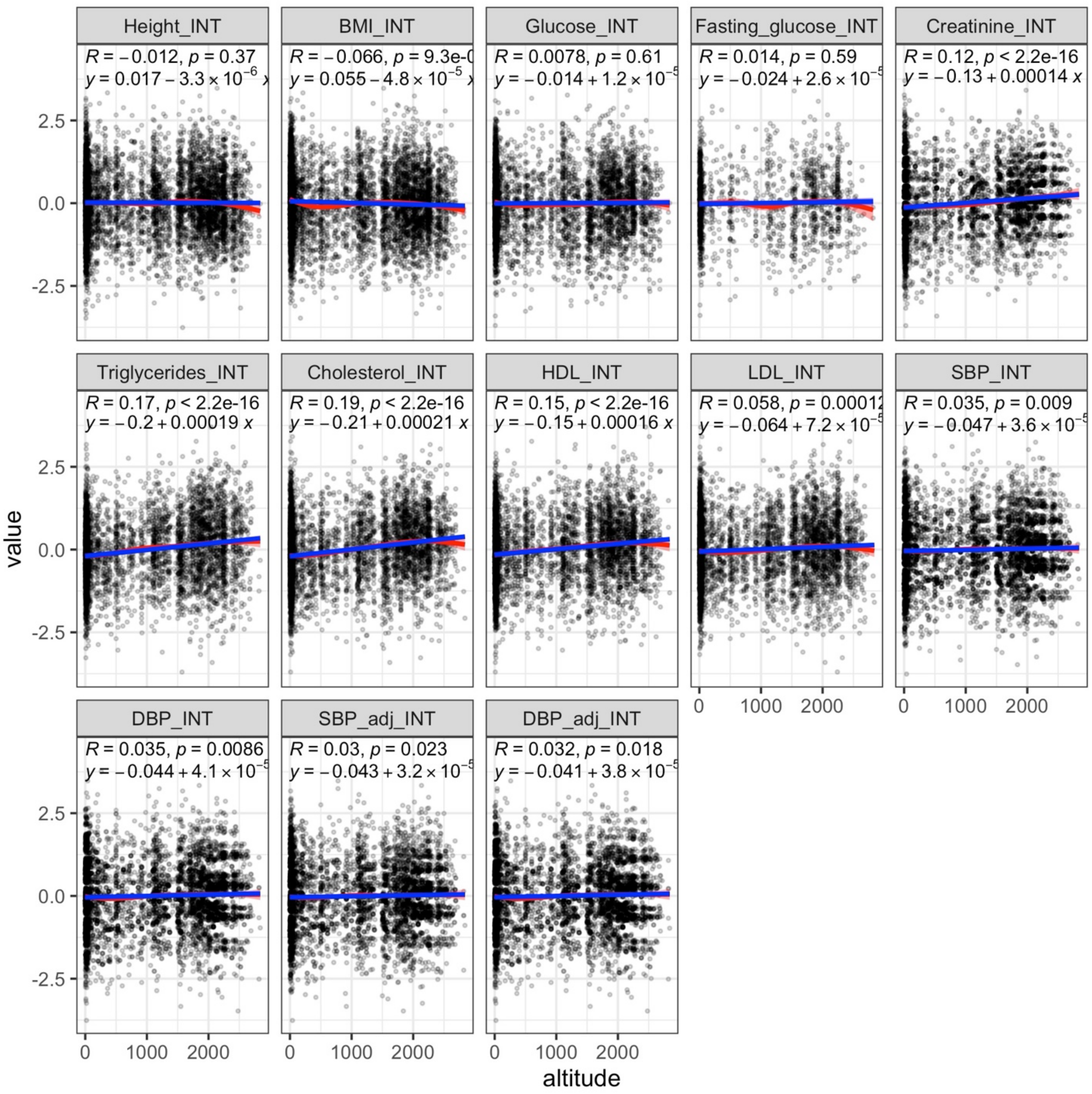


**Fig. S36. Complex trait variation by altitude.** To aid with visualization, a smoothed conditional mean line is shown using smoothing method loess (red) or lm/linear model (blue).

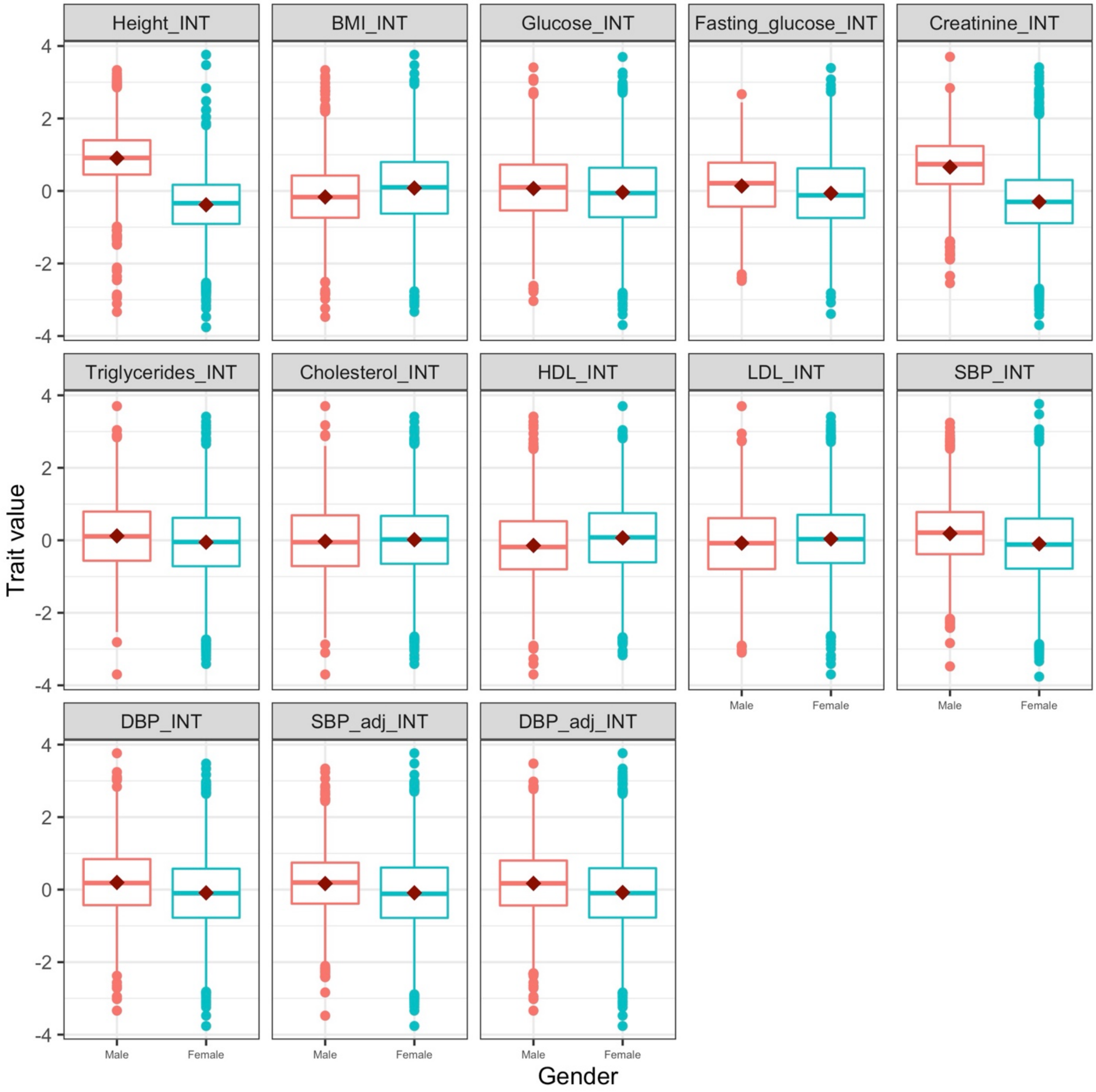


**Fig. S37. Complex trait variation by sex.** Box plots of trait values are shown for males (red) and females (blue).

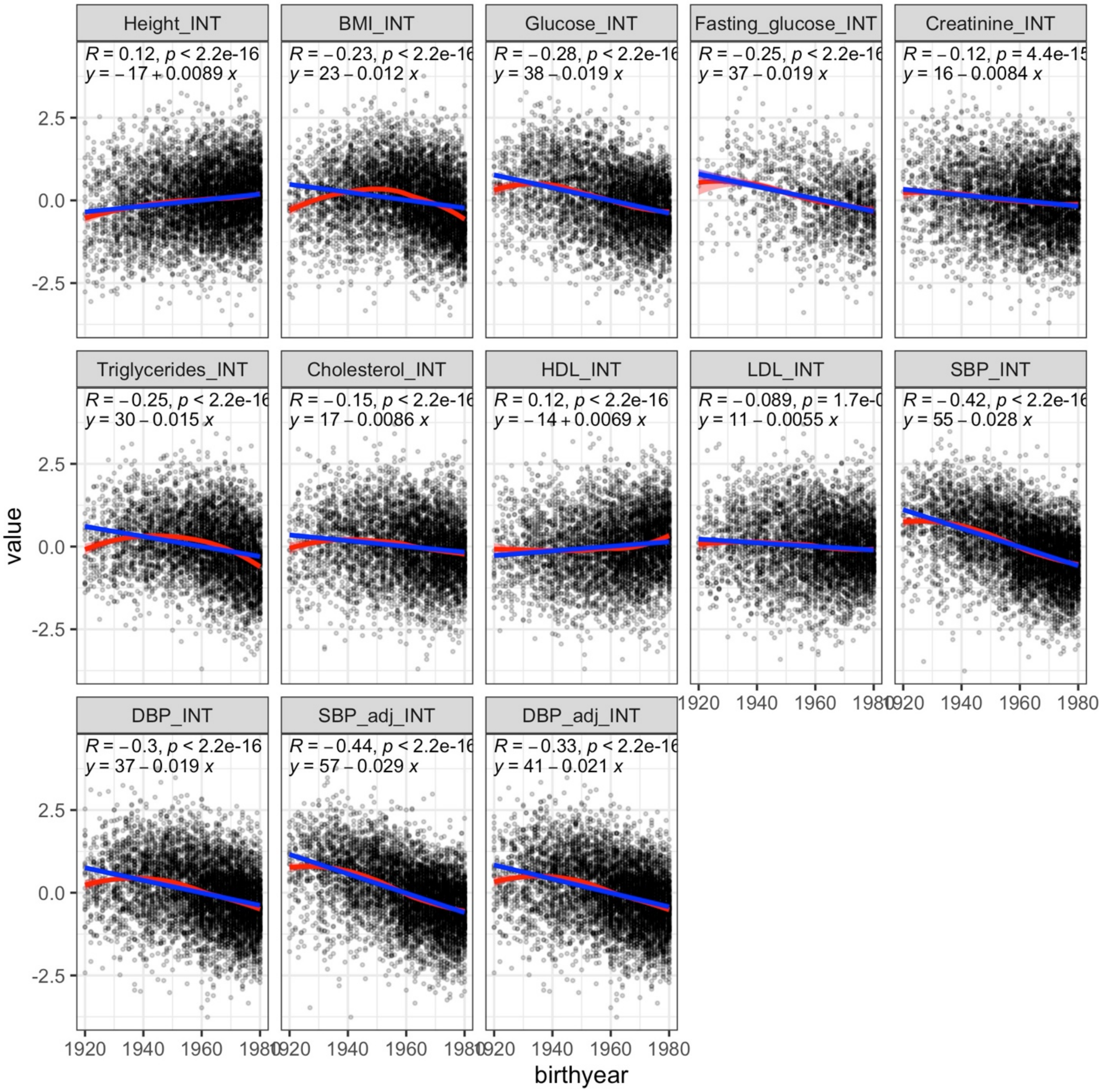


**Fig. S38. Complex trait variation by birth year.**

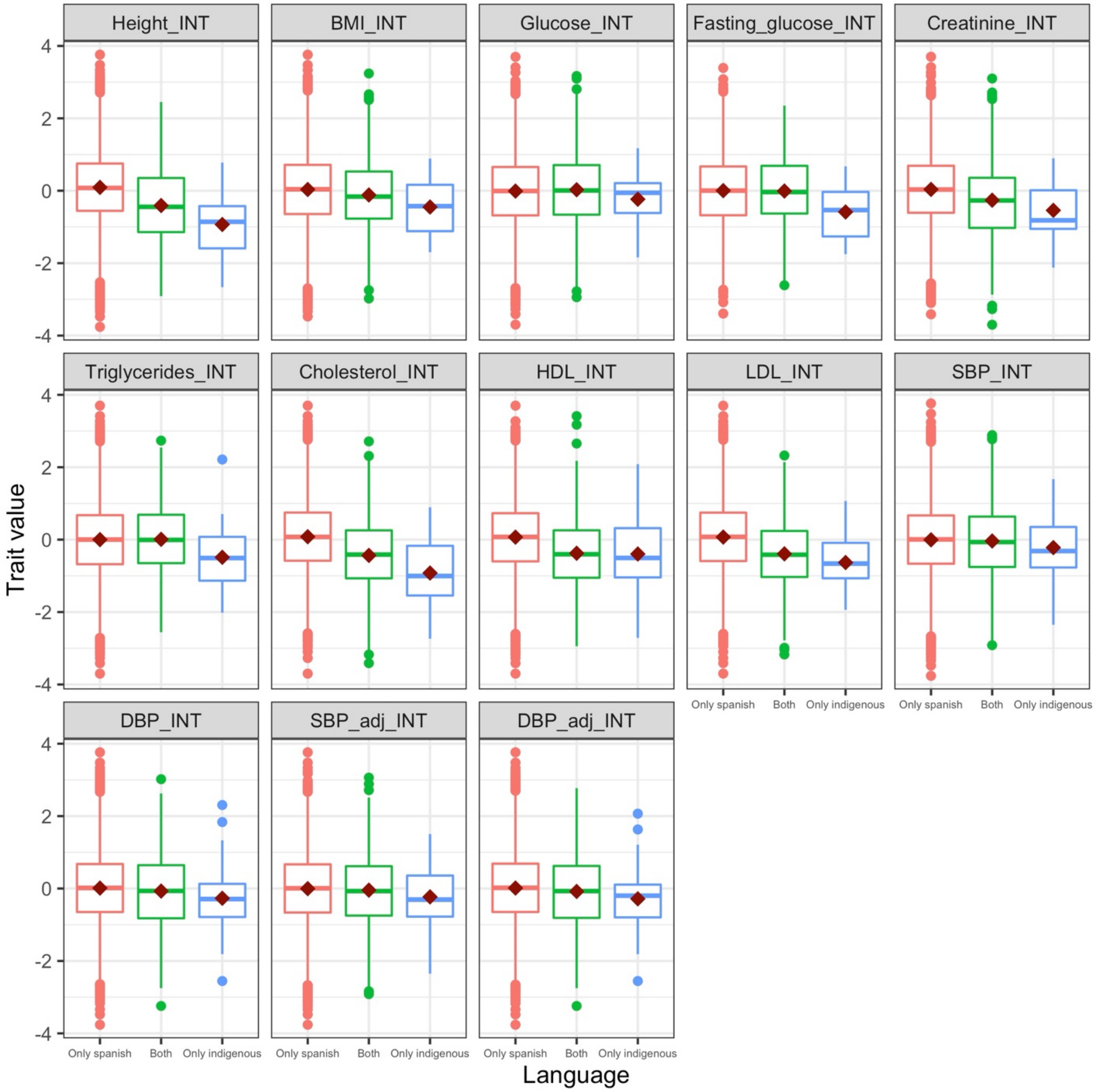


**Fig. S39. Complex trait variation by indigenous heritage.** Box plots of trait values are shown for individuals that speak only Spanish (red), that speak both Spanish and an indigenous language (green), and that speak only an indigenous language (blue).

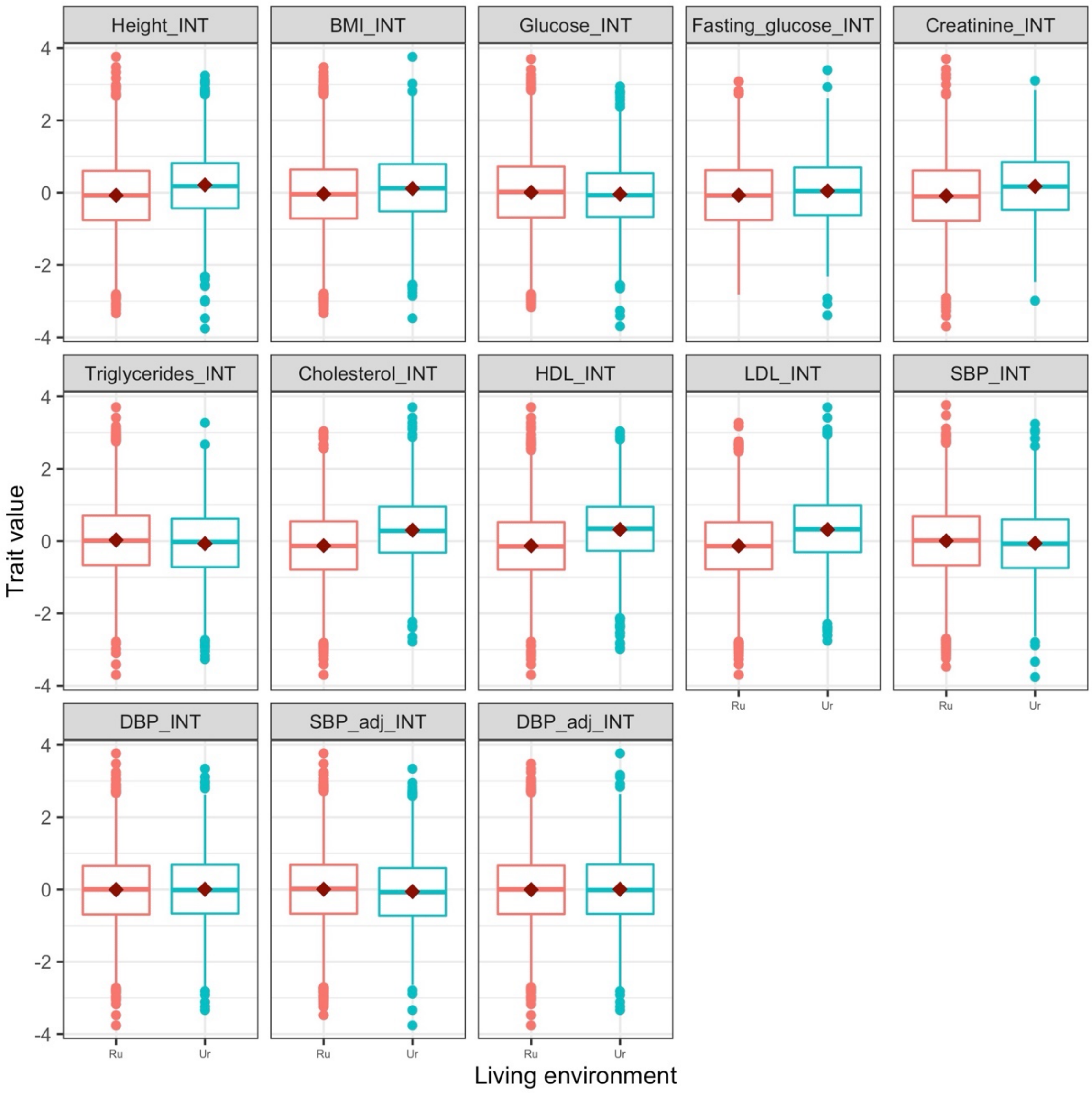


**Fig. S40. Complex trait variation by rural or urban living environment.** Box plots of trait values are shown for individuals that live in a rural area (red) and urban area (blue).

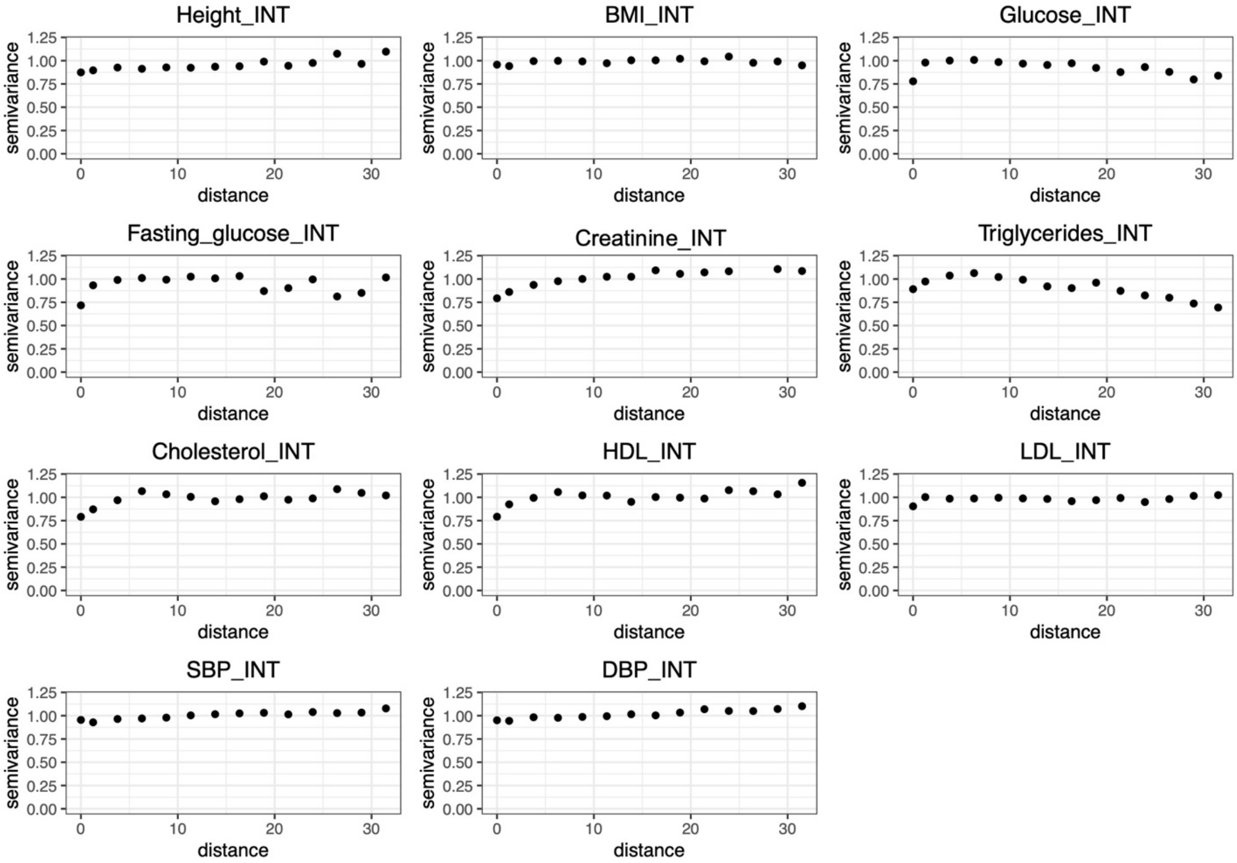


**Fig. S41. Variograms of complex traits.**

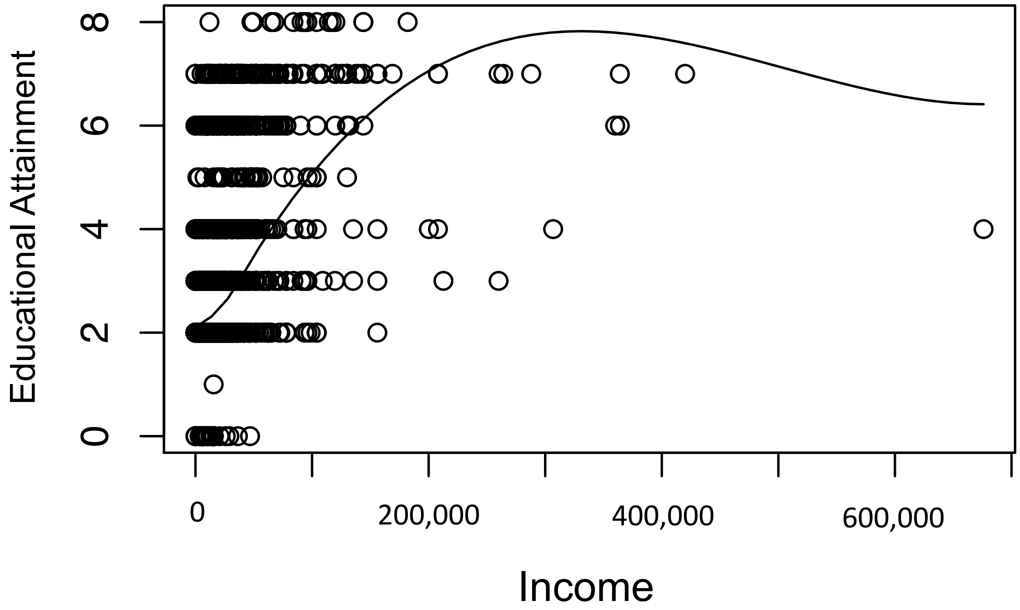


**Fig. S42. Correlation of educational attainment and income levels.**

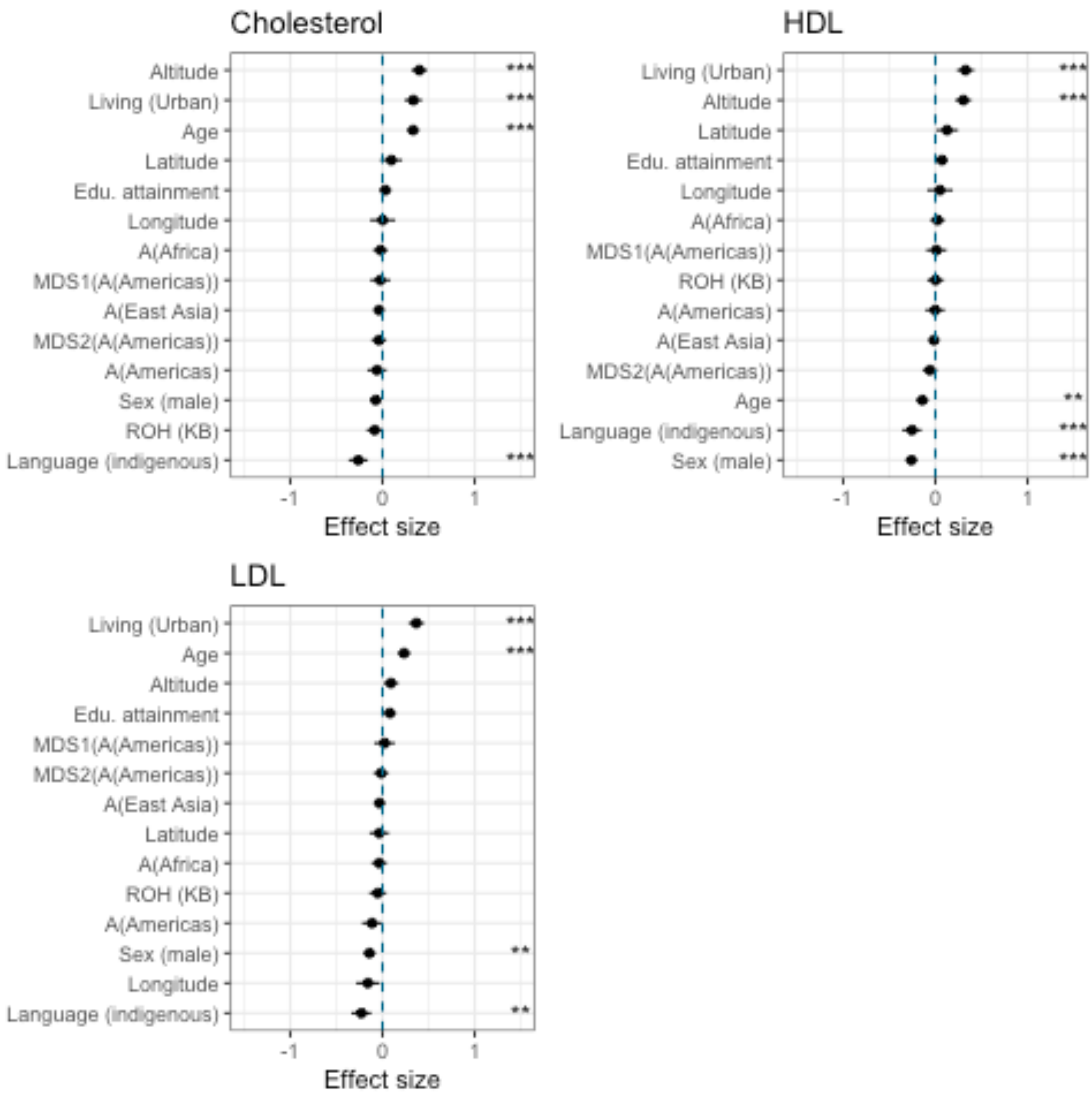


**Fig. S43. Mixed model results for Cholesterol related traits.** The plot shows effect size estimates and confidence intervals from a mixed model analysis. All quantitative predictors are centered and scaled by 2 standard deviations. Asterisks indicate significance of the effect of a predictor after Bonferroni correction (** $P < 10^{-5}$, *** $P < 10^{-6}$)

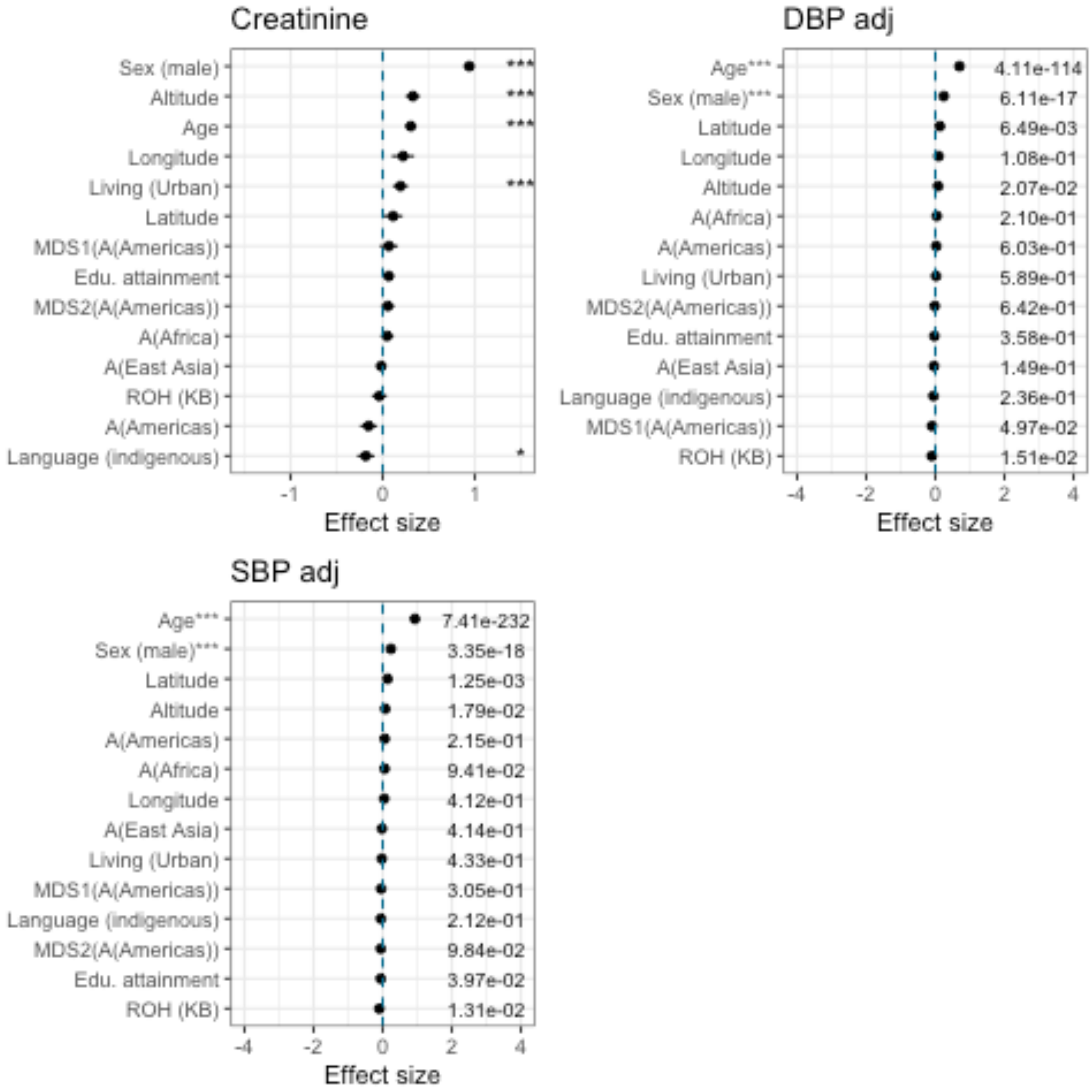


**Fig. S44. Mixed model results for Creatinine and blood pressure.** Blood pressure plots need to be updated to show asterisks instead of p-values. The plot shows effect size estimates and confidence intervals from a mixed model analysis. All quantitative predictors are centered and scaled by 2 standard deviations. Asterisks indicate significance of the effect of a predictor after Bonferroni correction (** $P < 10^{-5}$, *** $P < 10^{-6}$)

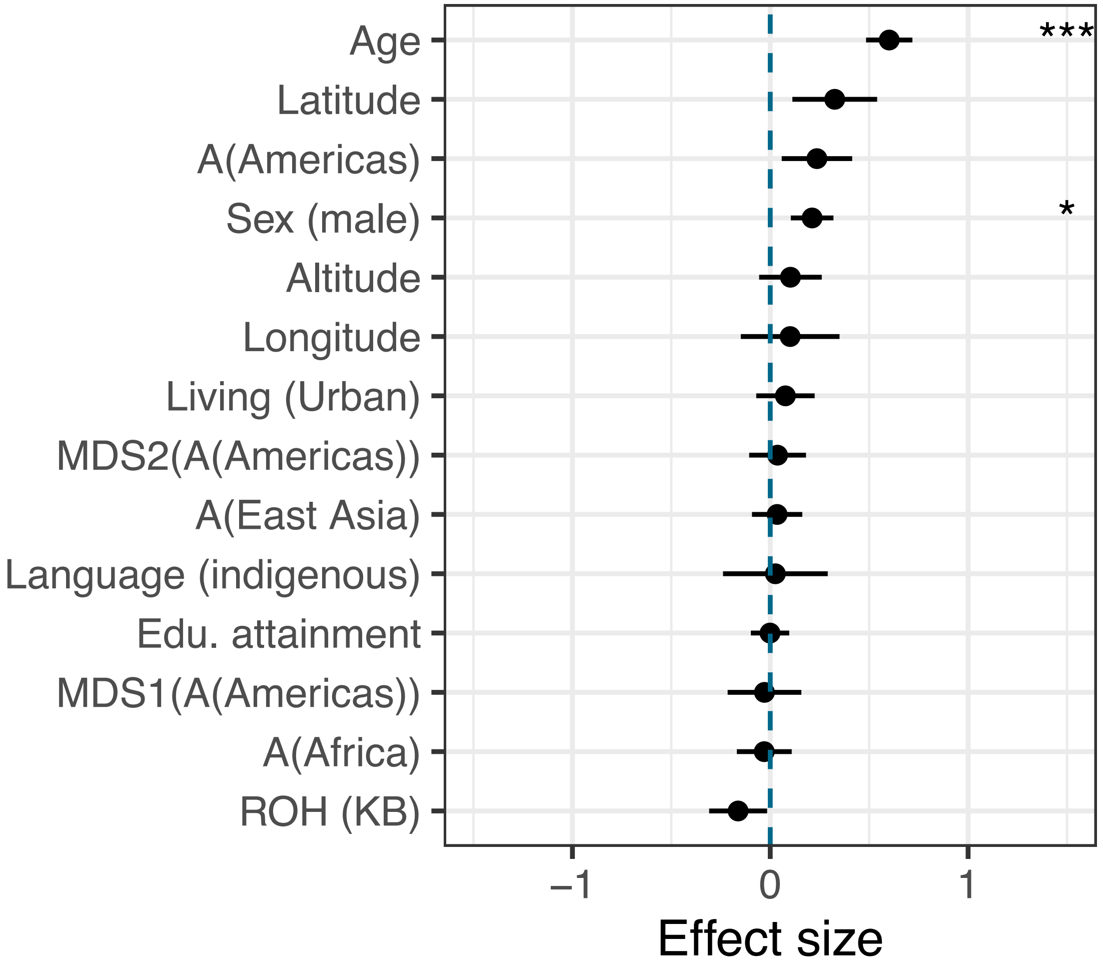


**Fig. S45. Mixed model results for fasting glucose.** The plot shows effect size estimates and confidence intervals from a mixed model analysis. All quantitative predictors are centered and scaled by 2 standard deviations. Asterisks indicate significance of the effect of a predictor after Bonferroni correction (** $P < 10^{-5}$, *** $P < 10^{-6}$).

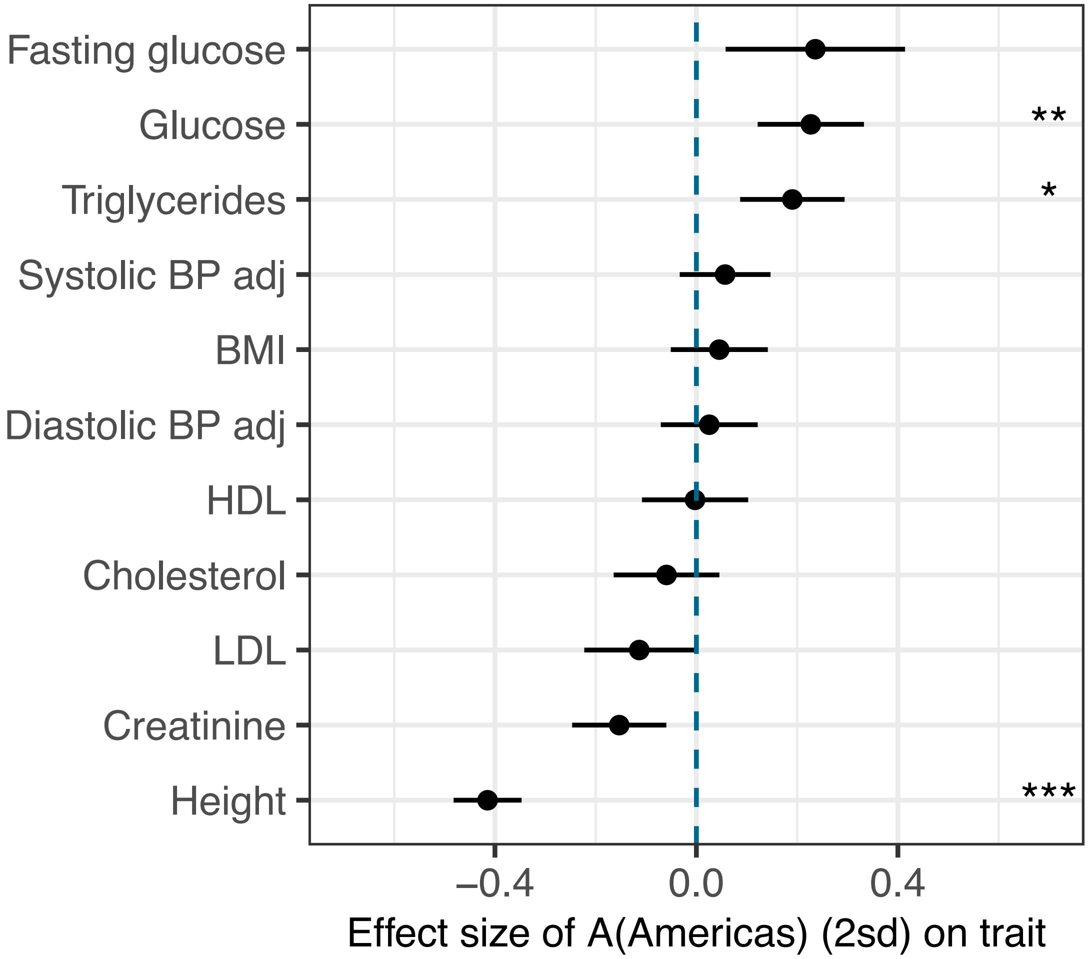


**Fig. S46. Mixed model results for ancestries from Central America.** The plot shows effect size estimates and confidence intervals from a mixed model analysis. A(Americas) predictor is centered and scaled by 2 standard deviations. Asterisks indicate significance of the effect of a predictor after Bonferroni correction (** $P < 10^{-5}$, *** $P < 10^{-6}$)

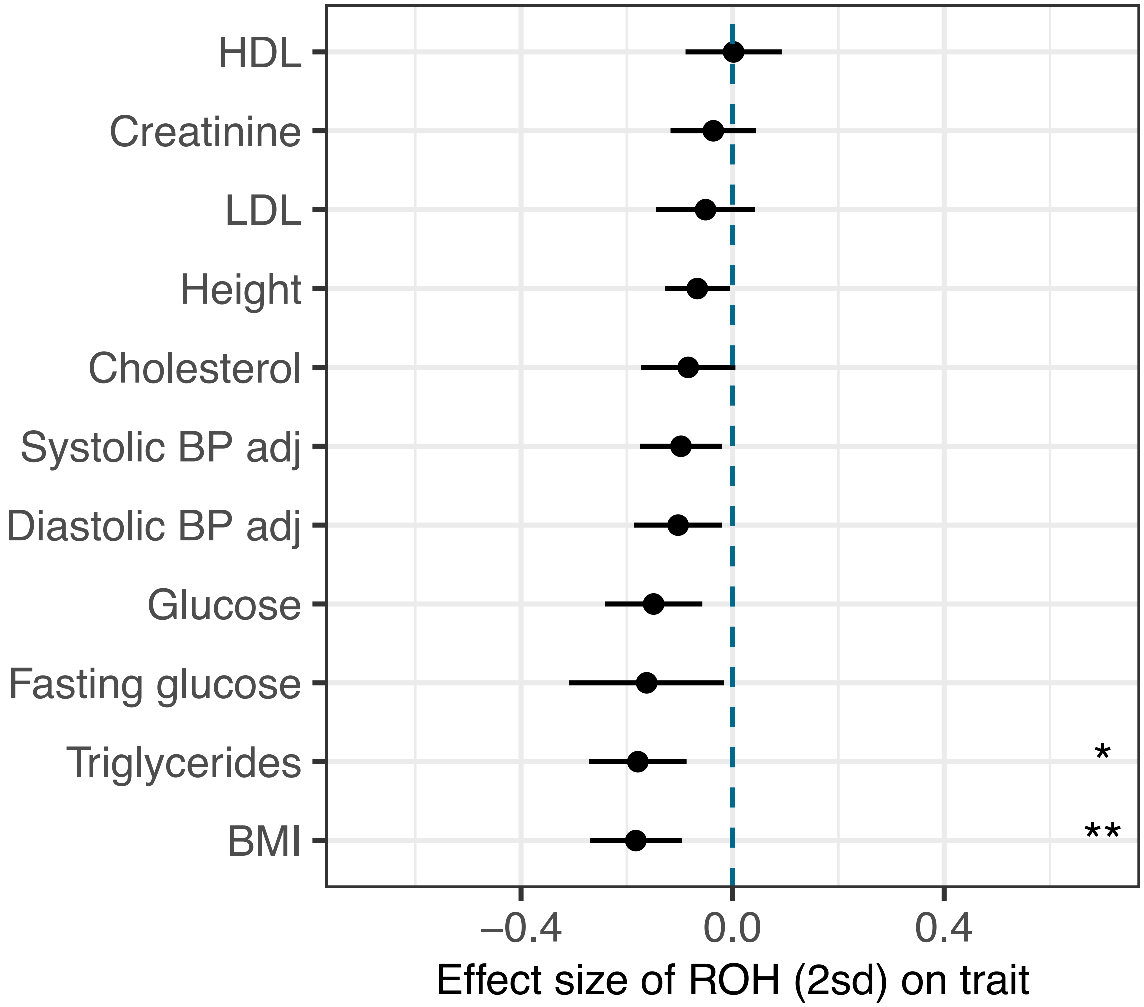


**Fig. S47. Mixed model results for sROH carried by an individual.** The plot shows effect size estimates and confidence intervals from a mixed model analysis. ROH predictor is centered and scaled by 2 standard deviations. Asterisks indicate significance of the effect of a predictor after Bonferroni correction (** $P < 10^{-5}$, *** $P < 10^{-6}$)

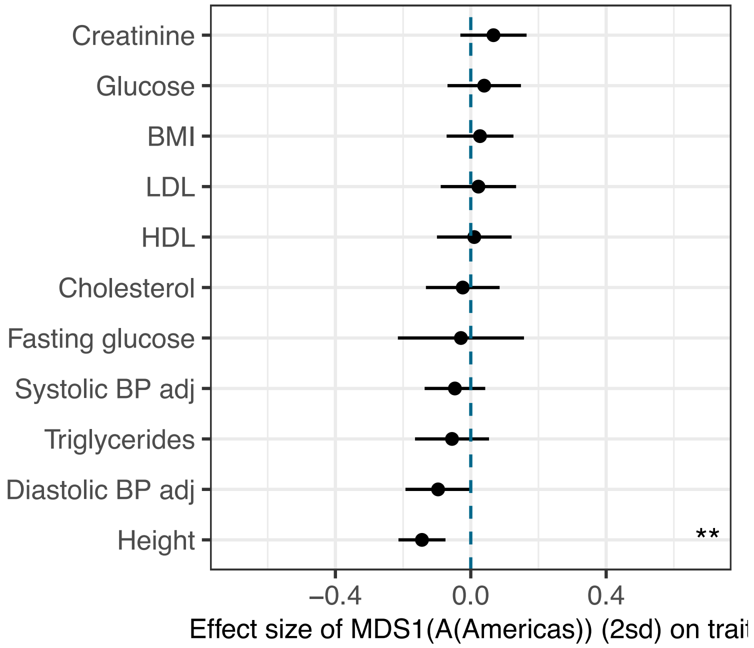


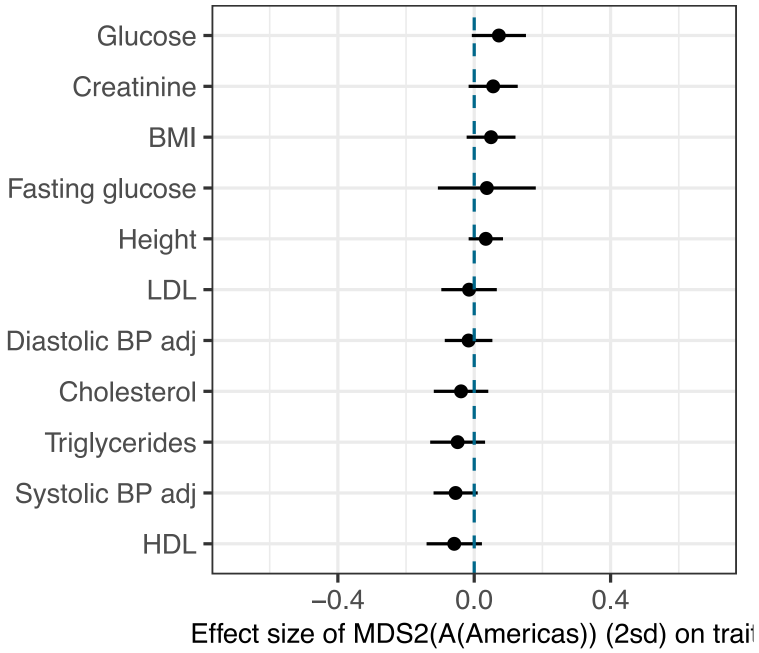


**Fig. S48. Mixed model results for two MDS axes reflecting genetic variation within Central America.** The plot shows effect size estimates and confidence intervals from a mixed model analysis. MDS axes are centered and scaled by 2 standard deviations. Asterisks indicate significance of the effect of a predictor after Bonferroni correction (** $P < 10^{-5}$, *** $P < 10^{-6}$)

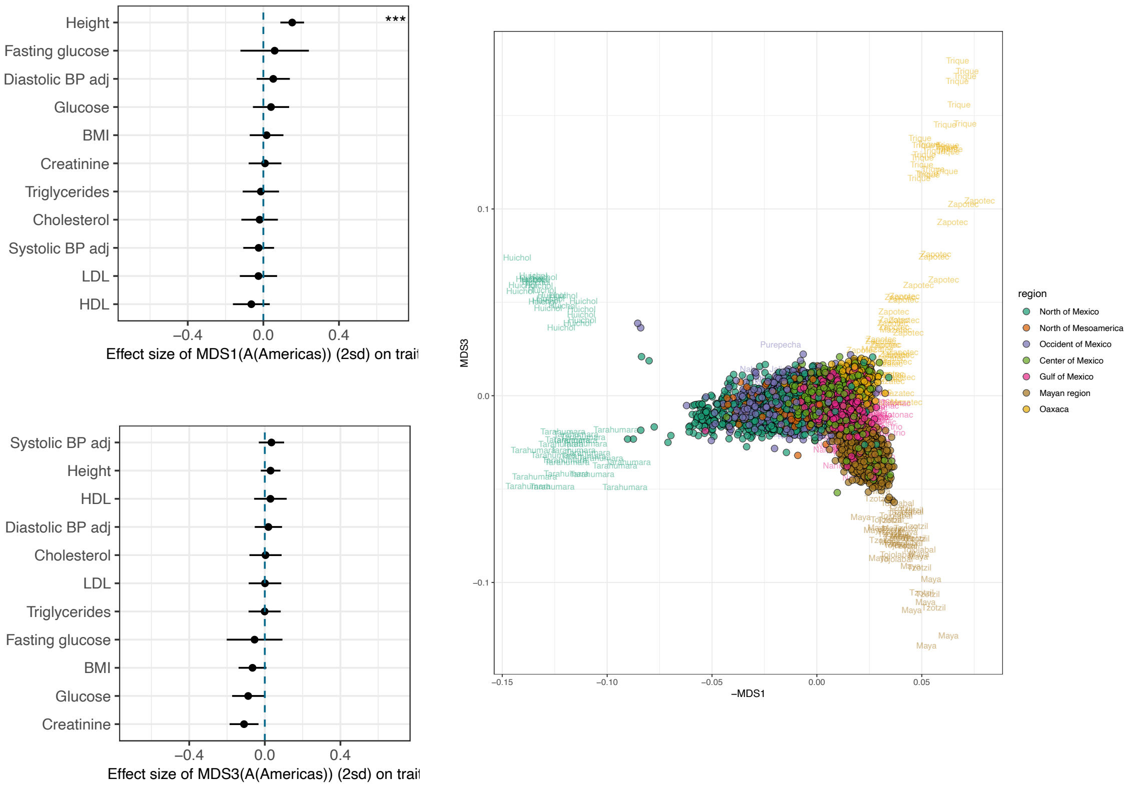


**Fig. S49. Mixed model results for two MDS axes reflecting genetic variation within Central America (using MAAS-MDS run on MXB and NMDP excluding Tojolabal individuals).** MDS 3 axis reflects variation among the groups from Oaxaca and from the Mayan regions. The plot shows effect size estimates and confidence intervals from a mixed model analysis. MDS axes are centered and scaled by 2 standard deviations. Asterisks indicate significance of the effect of a predictor after Bonferroni correction (** $P < 10^{-5}$, *** $P < 10^{-6}$)

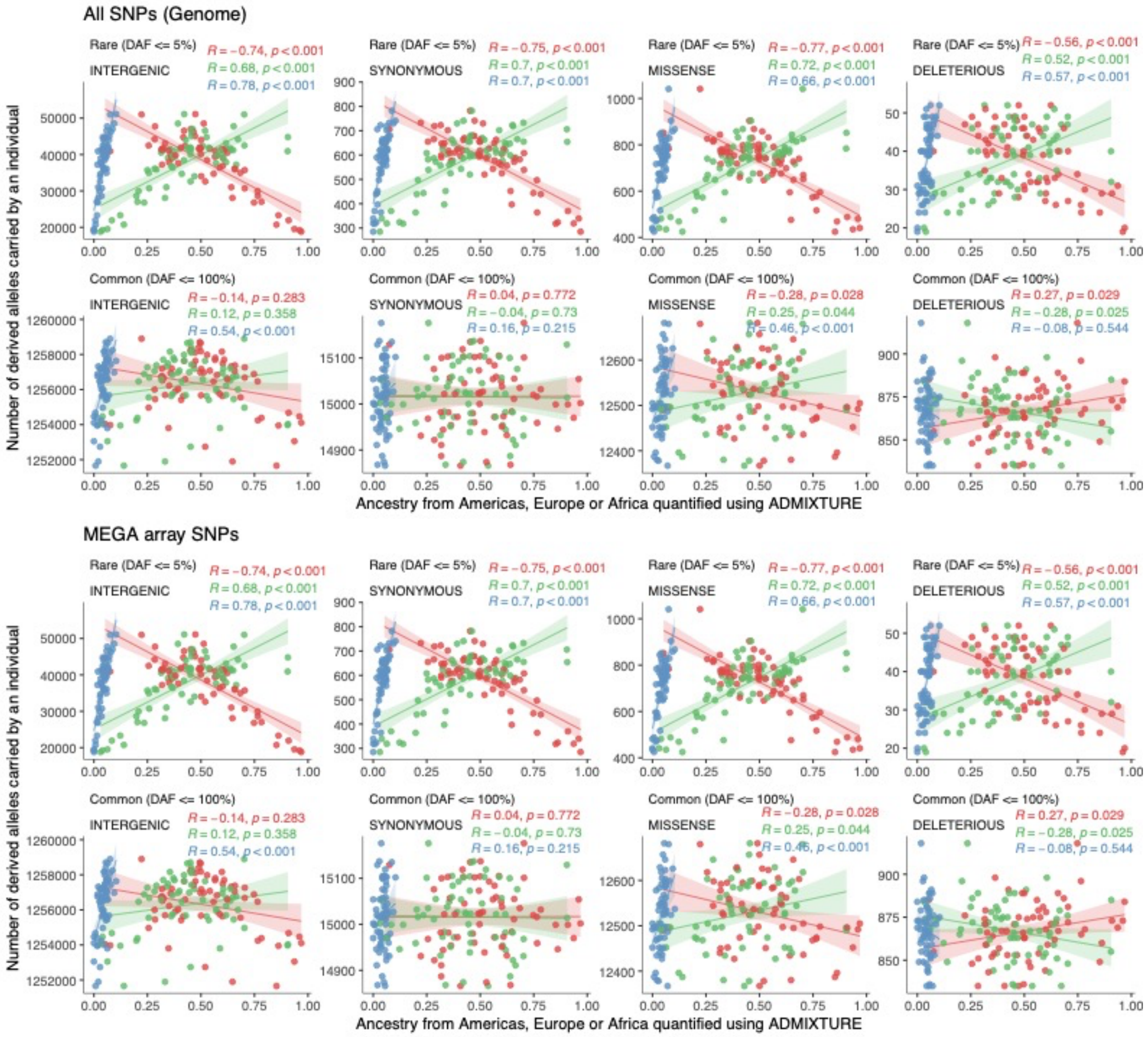


**Fig. S50. Mutation burden in 1000G MXL individuals as a function of their ancestries from Central America, Western Europe, and West Africa.** Colors show ancestries from Central America (red), Western Europe (green) or West Africa (blue). The top panel (first and second rows) shows mutation burden computed on All SNPs (Genome) while the bottom panel (third and fourth row) shows mutation burden computed on only the subset of MEGA array SNPs. A significant correlation between mutation burden and ancestry is seen for rare mutation burden in both the MEGA array SNPs and All SNPs (Genome) due to differential bottlenecking histories reflected in different ancestry proxies. In contrast, a significant correlation between mutation burden and ancestry is seen for total mutation burden only in the MEGA array SNPs, and not in All SNPs implying an ascertainment effect as the source of this signal. Thus, rare mutation burden shows a robust correlation with ancestry proxies, while total mutation burden is uncorrelated with ancestry proxies.

**Rural**
Spearman’s correlation = 0.075
p-value = 1.447 x 10^-6

**Urban**
Spearman’s correlation = 0.03
p-value = 0.2224

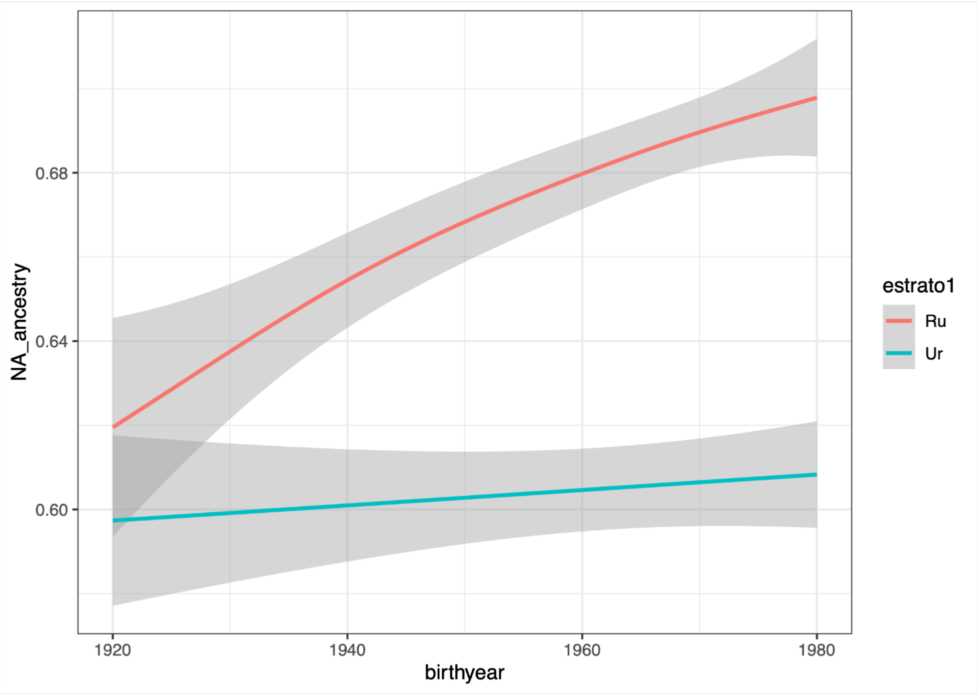


**Fig. S51. Ancestries from Central America as a function of birth year in rural and urban localities.**

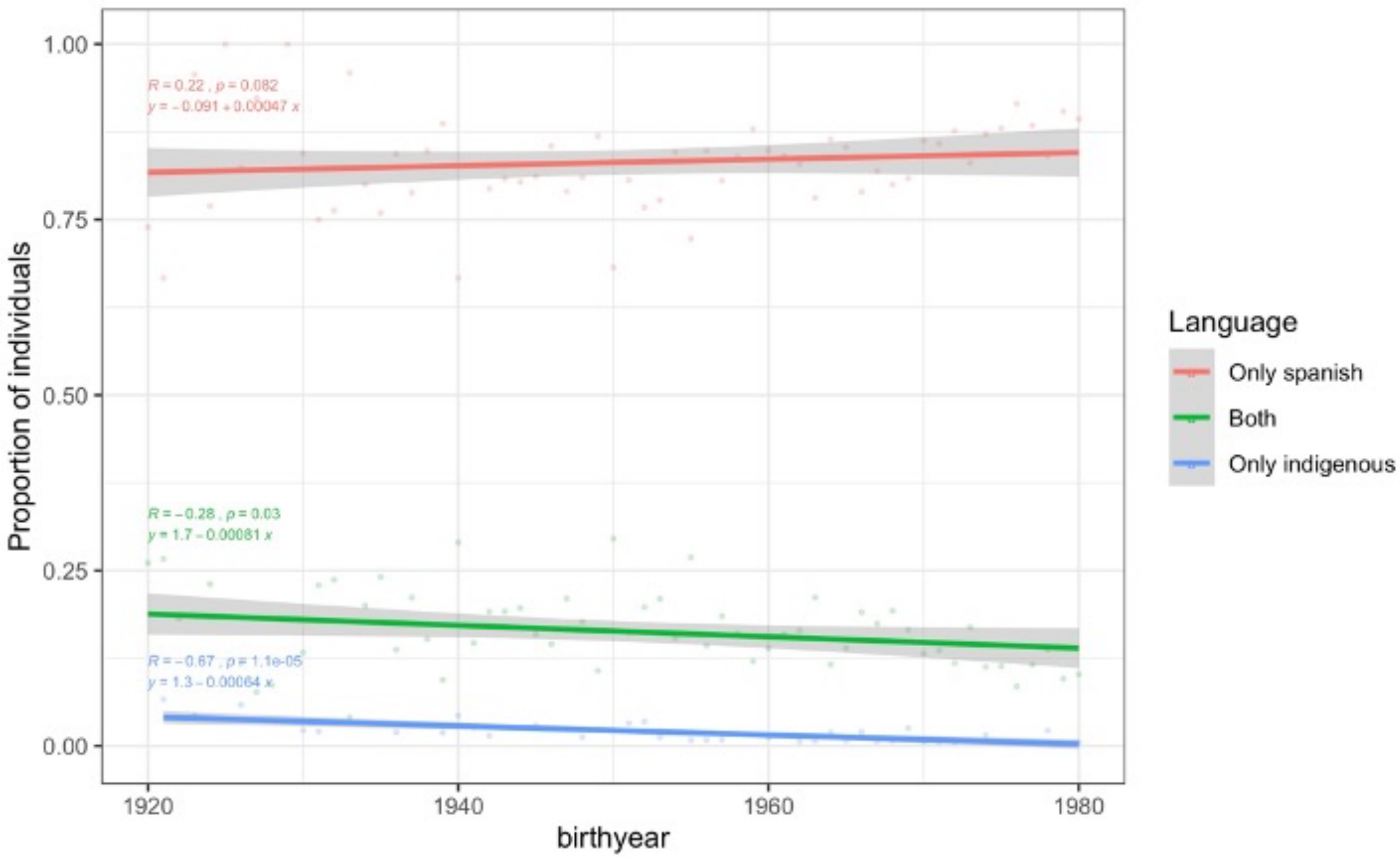


**Fig. S52. Proportion of individuals that speak Spanish, an indigenous language or both, as a function of birth year.**

**Supplementary Tables**

| Characteristic | Number | (%) |
|---|---|---|
| **Age** | | |
| **20-39** | **3295** | **55.13** |
| **40-59** | **1808** | **30.25** |
| **60-79** | **763** | **12.77** |
| **80+** | **111** | **1.86** |
| **Sex** | | |
| **Male** | **1820** | **30.45** |
| **Female** | **4157** | **69.55** |

**Table S1. Age and sex distribution in the Mexican Biobank.** This is the distribution for the QC'ed dataset used for downstream analyses.

| Anthropometric traits | Urine biochemical traits | Health-related traits | Familial disease history | Socioeconomic covariates | Health-related covariates |
|---|---|---|---|---|---|
| Height | pH | Blood pressure | Diabetes | Salary | Fertility |
| BMI | Leukocytes | Diabetes treatment | Hypertension | Indigenous heritage | Contraceptive use |
| | Nitrites | Diabetes diagnosis | Cardiac disease | Literacy | Alcohol use |
| | Proteins | Vaccination history | Inbreeding | Perceived health status | Tobacco use |
| | Glucose | History of paludism | | Access to healthcare | |
| | Ketones | History of Dengue | | Disability | |
| | Blood | Fasting blood glucose levels | | Access to safe water | |
| | Hemoglobin | Renal disease indicators | | Mosquito exposure | |
| | | Tuberculosis diagnosis | | Proximity to pollution sources | |

**Table S2. List of anthropometric, disease and lifestyle variables in the Mexican Biobank.** Additionally, a number of biochemical traits analyzed in this study were measured from blood samples.

| | Native | European | African | East Asian | South Asian |
|---|---|---|---|---|---|
| Baja California | 52.8 | 40.7 | 5 | 1.1 | 0 |
| Baja California Sur | 47.6 | 45 | 4.7 | 2.1 | 0.6 |
| Sonora | 52.9 | 41.1 | 3.4 | 2.1 | 0.5 |
| Chihuahua | 41.7 | 51.2 | 5.1 | 1.3 | 0.8 |
| Coahuila | 53.1 | 40.2 | 5.5 | 0.7 | 0 |
| Nuevo Leon | 46.9 | 47 | 4.8 | 0.7 | 0.6 |
| Tamaulipas | 57.8 | 36.1 | 4.7 | 0.8 | 0.5 |
| Sinaloa | 43.3 | 50.1 | 4.8 | 1.2 | 0.6 |
| Durango | 51 | 41.5 | 6.1 | 0.9 | 0.5 |
| Zacatecas | 50.1 | 43.3 | 5.3 | 0.8 | 0.6 |
| San Luis Potosi | 74 | 22.1 | 3.1 | 0.5 | 0 |
| Nayarit | 48.8 | 43.5 | 6.1 | 1.1 | 0.5 |
| Aguascalientes | 52.1 | 41.4 | 5.3 | 0.7 | 0.5 |
| Jalisco | 50.1 | 43.6 | 4.9 | 0.8 | 0.6 |
| Colima | 50.4 | 42 | 5.9 | 1 | 0.7 |
| Guanajuato | 53.7 | 40.6 | 4.6 | 0.7 | 0 |
| Queretaro | 64.6 | 31 | 3.5 | 0.7 | 0 |
| Veracruz | 73.8 | 19.8 | 5.2 | 0.8 | 0 |
| Michoacan | 59.9 | 34.5 | 4.3 | 0.7 | 0.6 |
| Hidalgo | 77 | 20.4 | 1.8 | 0.5 | 0 |
| Cd. Mexico | 67.9 | 28.1 | 2.9 | 0.7 | 0 |
| Edo. Mex | 69 | 27.4 | 2.4 | 0.8 | 0 |
| Tlaxcala | 79.5 | 18.4 | 1.3 | 0 | 0 |
| Guerrero | 69.9 | 21.7 | 5.4 | 2.3 | 0.7 |
| Puebla | 82.7 | 14.9 | 1.6 | 0 | 0 |
| Oaxaca | 89.9 | 7.8 | 1.7 | 0 | 0 |
| Morelos | 71.3 | 24.3 | 3.2 | 0.5 | 0.6 |
| Tabasco | 73.5 | 19.1 | 6.1 | 0.6 | 0.5 |
| Chiapas | 84 | 12.2 | 2.9 | 0 | 0.5 |
| Campeche | 78.8 | 16.1 | 3.8 | 0.6 | 0.5 |
| Yucatan | 79.3 | 17.2 | 2.4 | 0 | 0.7 |
| Quintana Roo | 77.1 | 18.2 | 3.4 | 0.7 | 0.6 |
| Average MXB | 65.6 | 29.1 | 4.01 | 0.82 | 0.50 |

**Table S3. Estimation of admixture proportions by state in MXB.** State with the highest ancestry from any source is highlighted.

| Cultural group | ID | Mesoamerican region | Latitude | Longitude |
|---|---|---|---|---|
| Seri | SER | North of Mexico | 29 | -112.15 |
| Tarahumara | TAR | North of Mexico | 27.75 | -107.17 |
| Tepehuano | Tep | North of Mesoamerica | 23.48 | -104.39 |
| Huichol | HUI | Occident de Mexico | 21.17 | -104.08 |
| Nahua_Jaliso | NAJ | Occident de Mexico | 19.5 | -103.5 |
| Purepecha | PUR | Occident de Mexico | 19.75 | -101.5 |
| Totonac | TOT | Gulf of Mexico | 20 | -97.8 |
| Nahua_Puebla | NXP | Gulf ofMexico | 19.97 | -97.62 |
| Nahua_trio | NFM | Gulf of Mexico | 19.93 | -97.62 |
| Nahua_Guerrero | NAG | Occident de Mexico | 17.89 | -99.13 |
| Trique | TRQ | Oaxaca | 17.18 | -97.95 |
| Zapotec | ZAP.N | Oaxaca | 17.41 | -96.69 |
| Zapotec | ZAP.S | Oaxaca | 17.23 | -96.23 |
| Mazatec | MAZ | Oaxaca | 18.33 | -96.33 |
| Tzotzil | TZT | Mayan region | 16.83 | -92.67 |
| Tojolabal | TOJ | Mayan region | 16.5 | -92 |
| Lacandon | LAC | Mayan region | 16.75 | -91.25 |
| Maya | MYA.Q | Mayan region | 19.58 | -88.58 |
| Maya | MYA.C | Mayan region | 20.37 | -90.05 |
| Maya | MYA.Y | Mayan region | 21.17 | -88.14 |

**Table S4. NMDP cultural groups and their designation into Mesoamerican regions.**

| | Native ancestry | African ancestry | European ancestry |
|---|---|---|---|
| nROH | 287675 | 429 | 23015 |
| Local ancestry (Mb) | 459820.79 | 2070.33 | 44006.19 |
| nROH/localancestry (MB) | 0.626 | 0.207 | 0.523 |

**Table S5. Number of ROH by local ancestry tracts.**
